# Affinity proteomic dissection of the human nuclear cap-binding complex interactome

**DOI:** 10.1101/2020.04.20.048470

**Authors:** Yuhui Dou, Svetlana Kalmykova, Maria Pashkova, Mehrnoosh Oghbaie, Hua Jiang, Kelly R. Molloy, Brian T. Chait, Michael P. Rout, David Fenyö, Torben Heick Jensen, Ilya Altukhov, John LaCava

**Author notes:** Equal contribution (Yuhui Dou, Svetlana Kalmykova, Maria Pashkova). Correspondence to IA and JL.

## Abstract

A 5’, 7-methylguanosine cap is a quintessential feature of RNA polymerase II-transcribed RNAs, and a textbook aspect of co-transcriptional RNA processing. The cap is bound by the cap-binding complex (CBC), canonically consisting of nuclear cap-binding proteins 1 and 2 (NCBP1/2). The CBC has come under renewed investigative interest in recent years due to its participation in RNA-fate decisions via interactions with RNA productive factors as well as with adapters of the degradative RNA exosome - including the proteins SRRT (a.k.a. ARS2) and ZC3H18, and macromolecular assemblies such as the nuclear exosome targeting (NEXT) complex and the poly(A) exosome targeting (PAXT) connection. A novel cap-binding protein, NCBP3, was recently proposed to form an alternative, non-canonical CBC together with NCBP1, and to interact with the canonical CBC along with the protein SRRT. The theme of post-transcriptional RNA fate, and how it relates to co-transcriptional ribonucleoprotein assembly is abundant with complicated, ambiguous, and likely incomplete models. In an effort to clarify the compositions of NCBP1-, 2-, and 3-related macromolecular assemblies, including their intersections and differences, we have applied an affinity capture-based interactome screening approach, where the experimental design and data processing have been modified and updated to identify interactome differences between targets under a range of experimental conditions, in the context of label-free quantitative mass spectrometry. This study generated a comprehensive view of NCBP-protein interactions in the ribonucleoprotein context and demonstrates the potential of our approach to benefit the interpretation of complex biological pathways.

## 1 Introduction

All RNAs transcribed by RNA polymerase II (RNAPII) are modified at the 5’-end early during transcription with an N7-methylguanosine (m7G) linked in a 5’-to-5’ orientation (1, 2). The resulting m7G-cap structure is bound by the nuclear cap-binding complex (CBC), a heterodimer of NCBP1 (CBP80) and NCBP2 (CBP20) (3). NCBP2 binds directly to the cap, albeit with relatively low affinity; its cap-binding affinity is significantly enhanced by its heterodimerization with NCBP1 (4, 5), which further serves as a binding platform for different proteins that influence the progression of RNAs (i.e. ribonucleoproteins; RNPs) towards productive or destructive fates. Through its diverse protein interactions, the CBC is known to modulate various activities of RNAPII transcripts. During transcription, the CBC interacts with P-TEFb and promotes transcription elongation (6); it also interacts with the U4/U6•U5 tri-snRNP to stimulate pre-mRNA splicing (3, 7). ARS2 (SRRT, Uniprot gene symbols preferentially used throughout) joins the CBC, forming the CBC-ARS2 (CBCA) complex, which influences the fate of multiple types of RNAs (8, 9). CBCA can interact with ZC3H18, which may in turn recruit the nuclear exosome targeting (NEXT) complex or the poly(A) tail exosome targeting (PAXT) connection, directing bound RNAs to decay via the RNA exosome (10, 11). On snRNAs and a few independently transcribed snoRNAs, the CBCA complex may interact with PHAX, forming the CBCAP complex which stimulates nuclear export of snRNAs and the movement of snoRNAs to nucleoli (9, 12–14). Within elongating (messenger) mRNPs, the CBC interacts with ALYREF in the ‘transcription/export’ (TREX) complex, promoting mRNA export (15).

NCBP3 (5, 16), previously coined c17orf85 or ELG (17–19), was recently proposed to form an alternative CBC with NCBP1, capable of substituting for NCBP2 and suppressing the mRNA export defect caused by loss of NCBP2 (16). Previous reports described the association of NCBP3 with mRNPs to be splicing-linked, exon junction complex (EJC) independent, and CBC-dependent (17); yet, NCBP3 has been grouped with the EJC and the TREX complex on the basis of protein-protein interaction studies (18–20). Most recently, NCBP3 was shown to interact in vitro with both CBC (via NCBP1) and SRRT, separately and as a ternary complex (5). The complex composed of CBC, SRRT, and NCBP3 was shown to be mutually exclusive with PHAX and proposed to be part of an RNA-fate decision tree - similar decision forks between CBC, NELF-E or SRRT (5), and between SRRT, PHAX or ZC3H18 (8) have also been reported. Based on the above mentioned studies, Figure 1A illustrates an abridged narrative for proposed NCBP1, NCBP2 and NCBP3 interactions.

**Figure 1:**
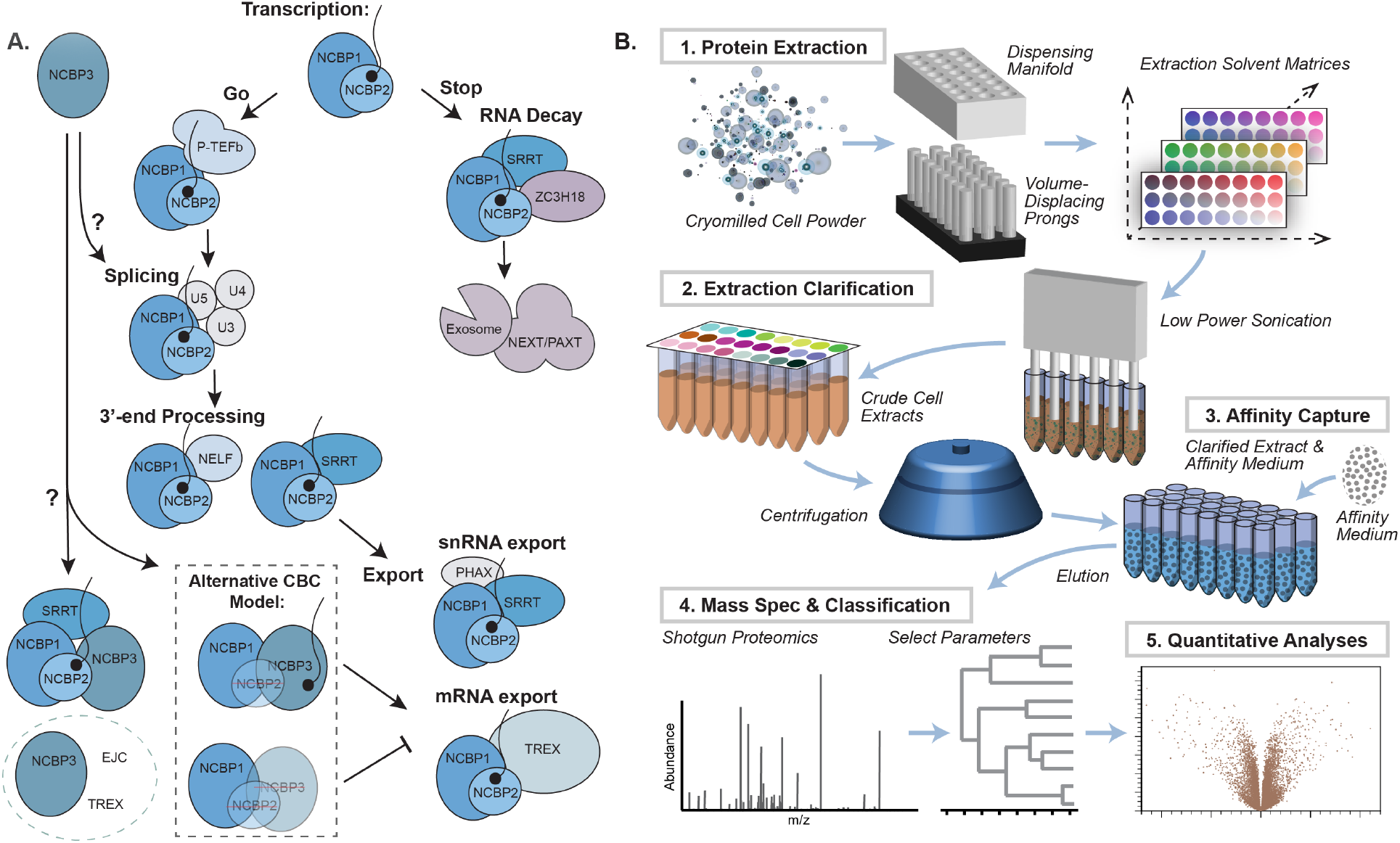
Summary of NCBP interactions and methodological approach. **A. NCBP interactions.** Right-side pathway: The canonical CBC consisting of NCBP1 and NCBP2 engage in early interactions guided by transcription, splicing, and 3’-end processing, to RNA export or decay (example interactors are shown for each summarized process; see main text for details). Left-side pathway: Putative NCBP3 functional interactions include: participation in splicing ((16–18, 20)); formation of an alternative CBC with NCBP1; and contributions to mRNA export (16). NCBP3 interacts with CBC and ARS2 (5, 16) and is affiliated with the EJC, and TREX complexes (16–18, 20). **B. Methodological approach.** Cryomilled cell powders are distributed with a dispensing manifold and macromolecules are extracted with 24 different extraction solutions (1). Brief sonication is applied to disperse and homogenize the extracts (2). After clarifying the extracts by centrifugation, affinity capture is performed (3) and protein eluates are subjected to MS analysis (4) and data processing (5).

Affinity capture, chiefly immunoprecipitation (IP), is arguably the most popular approach for characterizing target proteins’ physiological interacting partners. However, despite its common use, common implementations of this technique often suffer from shortcomings that include suboptimal parameterization of isolation conditions. Hence, significant improvements in performance may be obtained by customizing and optimizing the protocol at critical points (21, 22). To tap the full potential of the technique, we previously developed a platform to parallelize affinity capture and screen performance quality (23) - thus, this screen is a mode of quality assurance. Although numerous incremental protocol enhancements were built into this procedure (many described in (24)), the main advance is encompassed by the exploration of diverse protein extraction and capture conditions in a manner conceptually similar to crystallographic screening (25); details are illustrated in Figure 1B. In the present study, we retooled the screen to leverage label-free quantitative mass spectrometry (LFQ MS, e.g. reviewed in (26)). To expand our knowledge of the protein-protein interactions exhibited by NCBPs, we carried out a comparative analysis of proteins that co-IP with NCBP1, −2, and −3 from HeLa cell extracts under a variety of experimental conditions.

## 2 Results

### 2.1 Interaction screen and data analysis scheme

In our previous work, we primarily distinguished differences in the compositions of affinity captured protein complexes through the visual examination of Coomassie Brilliant Blue-stained SDS-polyacrylamide gels, followed by identification of the differentially enriched protein components by MALDI-TOF MS. This approach was successful on several multi-protein complexes we chose to examine (23), including those that form RNPs and/or metabolize RNA, and has enhanced our discovery potential on additional projects since (e.g. (27, 28)). The prior implementation, however, proved most suitable for the study of affinity enriched protein complexes that exhibited high enough yield of their individual constituent proteins to differentially detect differences by general staining, visually. We did not readily observe these characteristics when applying a 24-condition screen to NCBPs; exhibited for NCBP1 in Figure 2A. Thus to conduct a thorough study of NCBP interactomes, we modified and updated our procedure to: (i) pre-screen for experimental conditions that maximize sample diversity across a given space, (ii) accommodate control experiments and sufficient replicates for LFQ MS analysis of immunoprecipitated fractions, and (iii) provide bioinformatic resources useful for parsing and exploring interactome differences, statistically.

**Figure 2:**
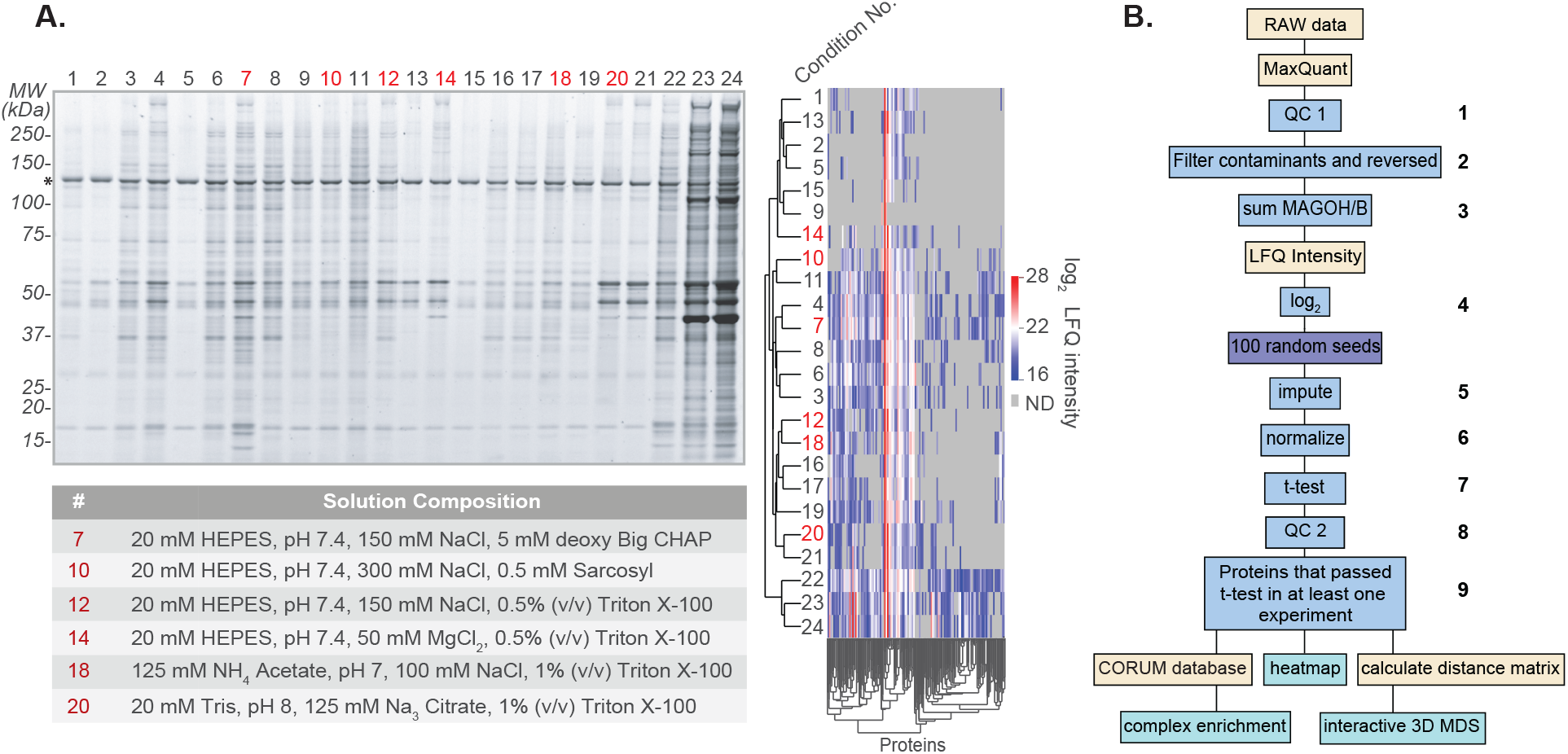
IP-MS pre-screen of NCBP1-LAP and depiction of data processing. **A. NCBP1-LAP pre-screen.** Coomassie stained SDS-polyacrylamide gel of co-IPs (left) and hierarchical clustering of the cognate MS data using log2 LFQ intensity (right). Gray coloring indicates a protein was not detected (ND). Six conditions, highlighted in red, were selected for subsequent quantitative screening; detailed solution compositions are listed in the table. **B. Bioinformatics pipeline.** After conducting an interactome screen on all three NCBPs and controls - using the conditions highlighted in panel A - the raw MS data were processed in MaxQuant (71) followed by post-processing (see Methods), summarized as follows: (1) inspected PTXQC quality control reports (72); (2) remove common contaminants and reversed protein sequences (FDR-control); (3) merged MS intensities for homologs MAGOH and MAGOHB; protein LFQ intensities were (4) log2 transformed, used to (5) impute missing values (multiple imputations with 100 random seeds), and (6) normalized to GFP intensities; (7) results from NCBP and control IPs were compared statistically and (8) imputation-derived false positives were removed; (9) proteins passing ANOVA were visualized by using complex enrichment (Figure 3A), heatmaps (Figure 3B), and multidimensional scaling (Figure 5).

After conducting an initial 24-condition IP / MS ‘pre-screen’ of NCBP1-LAP (localization and affinity purification tag; using anti-GFP affinity medium, see Methods), six conditions were chosen as exemplars for quantitative analysis (Figure 2A, red numbers). This was done in an effort to efficiently use the bandwidth of the screen while maximizing the protein-richness and breadth of interatomic differences. Thereafter 6-condition screens, with four replicates per condition, were applied to all LAP-tagged NCBPs and cognate controls (LAP-tag only) in anti-GFP IPs. Analysis of NCBP3-LAP demonstrated that the expression-level and yield of this protein did not reach comparable levels to NCBP1- and 2-LAP at the standard scale used in the screen; therefore, to more accurately assess the NCBP3-LAP interactome, we increased the scale and repeated IP-MS across four conditions, in three replicates each (see Methods). Once a complete experimental dataset for all three NCBPs was obtained, we applied the bioinformatic analyses outlined in Figure 2B. To enhance performance, we developed a novel approach to impute missing values in our data, a common challenge in LFQ MS (see Methods and Discussion). After statistical analysis was performed to determine which proteins were enriched in the NCBP-LAP IPs compared to controls, further analyses leveraging various visualization techniques were explored in an effort to reveal and emphasize differences in NCBP interactomes, presented below.

### 2.2 A complex-centric view of NCBP differences

To obtain a general, first-pass comparison of our samples, we explored the overlap between our NCBP data and protein complexes curated in the CORUM database (29). Figure 3A (‘complex enrichment plot’) displays a composite summary of the differences in protein complexes affinity captured with LAP-tagged NCBPs, including those illustrated in Figure 1A. The plot sacrifices details concerning the individual components of each complex in order to provide a functional summary of macromolecular differences.

**Figure 3:**
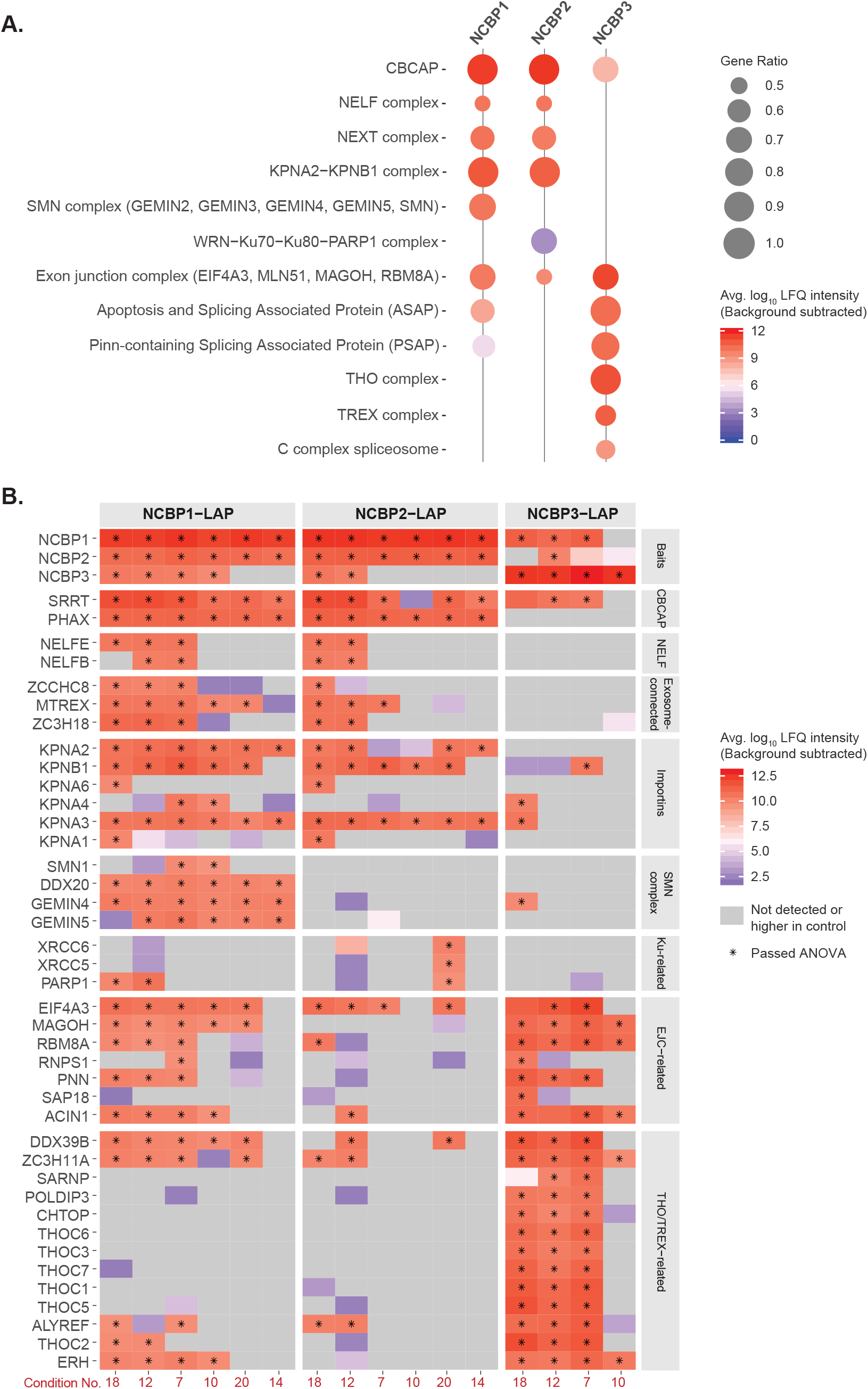
Complex- and protein-centric data visualizations. **A. Protein complex enrichment plot.** A composite representation of protein complexes observed across all IP-MS experiments, combined for each NCBP. Complexes were only represented in this plot if half or more of their constituent components passed statistical cut-offs (adjusted p-value ≤ 0.05 and log2 fold change ≥ 1) within the screen. This is represented by the ‘gene ratio.’ A minimum gene ratio of ≥ 0.5 was enforced for complexes with three or more components; for two-component complexes, both components must be present. Complex annotations were taken directly or modified from CORUM to avoid redundancy (29). The relative abundance of each complex captured is given as the control subtracted, average LFQ intensity of the constituent proteins (see Methods). Complex LFQ intensities are presented as their log10 values. A detailed interactive data interface, including results isolated from specific IP-MS experimental conditions, is available at: https://ncbps.shinyapps.io/complex_enrichment/. **B. Individual protein enrichments.** A summary heatmap with a focus on individual proteins, grouped based on selected CORUM complexes (manually inspected) and annotated protein functions. Control subtracted, average LFQ intensity of the quantified proteins are presented as their log10 values; negative or zero intensities are in grey. Proteins that passed the significance threshold (adjusted p-value ≤ 0.05 and log2 fold change ≥ 1) are marked with asterisks.

NCBP1 and 2 are largely described in the context of their cooperative activities in the CBC; represented here by e.g. CBCAP in sn(o)RNA transport (9), NELF (negative elongation factor) in 3’ end processing (30), NEXT in exosomal RNA decay (10), and the karyopherin import complex KPNA2-KPNB1 in the cytoplasmic-nuclear recycling of the CBC (31). However, we also observed potential independent/preferential protein complex associations for each: survival of motor neurons (SMN) complex, involved in snRNP biosynthesis and assembly (32), was co-enriched only with NCBP1. Some DNA-related complexes preferentially co-enriched with NCBP2, such as the WRN-Ku70-Ku80-PARP1 complex (33). Moving to NCBP3, we observed that it was neither appreciably associated with NELF (previously reported (16)) nor RNA exosome-related complexes (i.e. NEXT). Instead NCBP3 was more appreciably associated with EJC and EJC-affiliated complexes (ASAP, PSAP) and was even more contrastingly associated with THO (suppressors of the transcription defects of hpr1Δ mutants by over-expression) and TREX complexes as well as the spliceosomal C complex. The differential associations of NCBP1, 2, and 3 with EJC and THO/TREX are cross-validated and functionally dissected in a separate manuscript (Dou et al. submitted). Additional complexes are presented in a detailed interactive collection of enrichment plots online: https://ncbps.shinyapps.io/complex_enrichment/.

To reveal finer protein-level details, the heatmap, shown in Figure 3B was generated: NCBP1-LAP co-captures NCBP2 in all conditions tested and NCBP3 in four out of six conditions above our statistical cut-offs (adjusted p-value ≤ 0.05 and log2 transformed fold-change ≥ 1, see Methods; precise adjusted p-value and fold-change metrics can be found in the file: Supplemental_Data_pvalues_logfc.xlsx). NCBP2 co-captures NCBP3 in conditions 18 and 12, and NCBP3 captures NCBP2 in condition 12 only; although these proteins are also observed together in other IPs than those stated above, they do not surpass statistical cut-offs for co-enrichment in those conditions. Because NCBP2-LAP co-IPs NCBP3 and NCBP3-LAP reciprocally co-IPs NCBP2, in both cases along with NCBP1, these data support a model where NCBP3 association with NCBP1 (or, more precisely, NCBP1 affiliated macromolecules) is not mutually exclusive with NCBP2. NCBP1 and NCBP2 are always strongly co-associated, but their associations with NCBP3 appear to be less stable and the connection is lost in several conditions. All three NCBPs were associated with the protein SRRT, but only NCBP1 and NCBP2 were also observed to be associated with PHAX (confirming previous reports (5, 16)). An extended supplemental heatmap (Supplemental_heatmap.pdf) displays additional proteins, including spliceosomal proteins.

### 2.3 Re-examining the relative abundance of NCBP1:NCBP2

NCBP1 and NCBP2 were robustly co-associated across all NCBP1/2 IPs, yet their co-IP profiles also exhibited features distinguishing them from one another. Perhaps, outside of the context of the CBC, NCBP1 and/or NCBP2 participate in the formation of independent macromolecules. We reasoned that support, or opposition, for this idea may arise by examining the relative abundance of NCBPs present in their reciprocal IPs.

To address this question, we plotted NCBP MS-derived iBAQ intensity ratios, as displayed in Figure 4. iBAQ intensity ratios can be used as a proxy for protein proportions because they are normalized by the number of theoretically observable peptides for each protein (34); larger proteins encompass a greater number of observable peptides, smaller proteins, fewer. Examining the relative abundances of NCBPs revealed that the relationship between NCBP1 and NCBP2 was atypical. NCBP3 IPs illustrate why this is so: NCBP3 is more abundant than NCBP1 and NCBP2 when NCBP3 is the IP target (Figure 4, right side of plot). The target protein nearly always exhibits apparent superstoichiometry to its in vivo interaction partners in IP fractions; among other possibilities, in a well optimized IP this can usually be attributed to the antibody-antigen interaction being the highest affinity interaction in the mixture. This is also the reason why the same protein (initial target) then becomes substoichiometric in reciprocal IPs, that are commonly used to confirm IP-based interactions. In line with this expectation, and as mentioned earlier, NCBP3 is less abundant, or missing, in IPs targeting NCBP1 and NCBP2.

**Figure 4:**
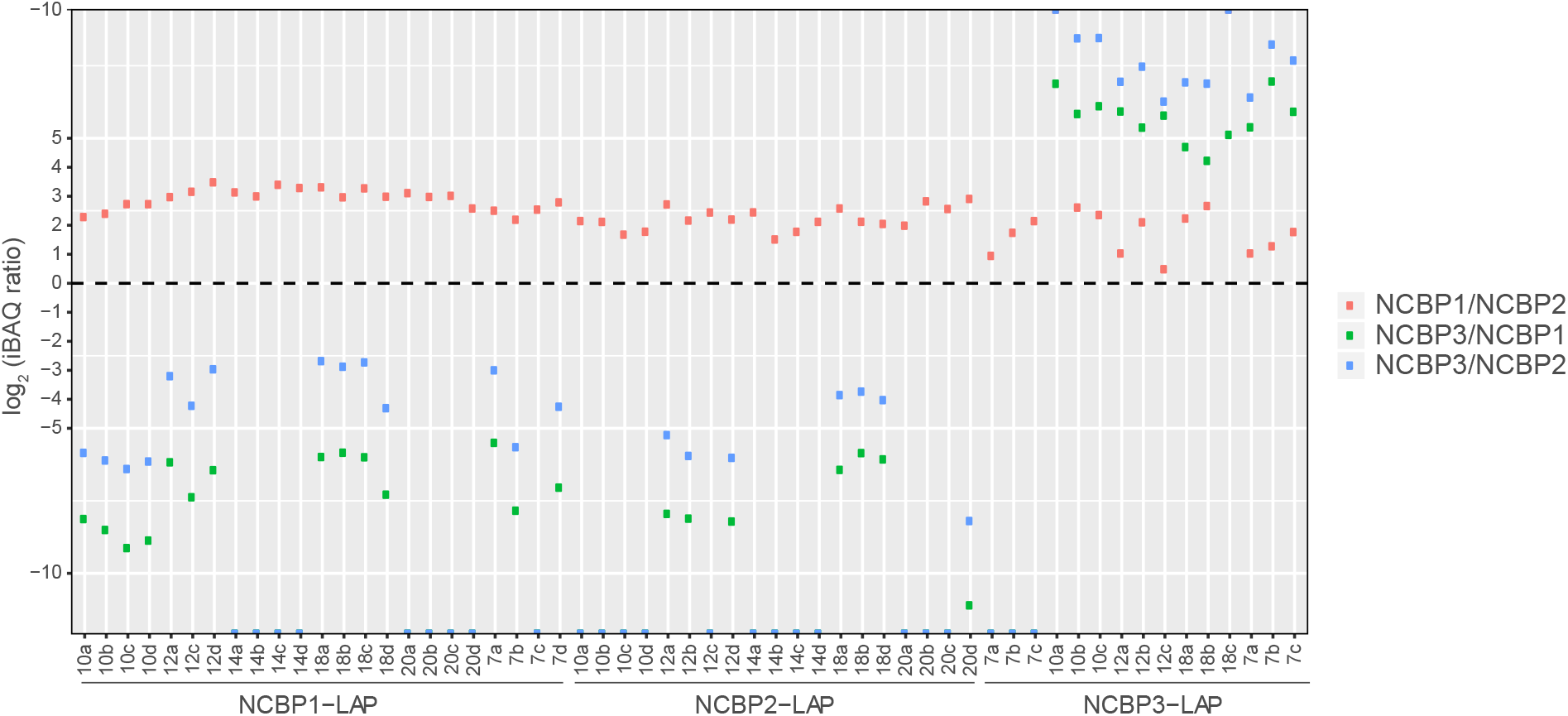
Relative copy numbers of NCBPs. **Ratios of NCBPs in individual IPs.** The log2 ratios of the NCBP iBAQ intensities were calculated for each IP experiment as a proxy for relative copy number - shown as NCBP1/NCBP2 (red), NCBP3/NCBP1 (green), NCBP3/NCBP2 (blue). NCBP1 has the highest copy number in NCBP1-LAP IPs and NCBP2-LAP IPs, while NCBP3 has the highest copy number in NCBP3 IPs.

When NCBP1 is the IP target, it also exhibits apparent superstoichiometry. However, NCBP2 exhibited a surprising trend as the target of IP: NCBP2 appears substoichiometric to NCBP1, exhibiting several-fold lower abundance. This trend is prevalent across all the NCBP2 IPs and suggests that while NCBP1/2 stably interact (the expected result), NCBP1 might be present in more than one copy per NCBP2 in an RNP context (see Discussion).

### 2.4 Contextualization of NCBPs in mRNP maturation

We sought to integrate and parse all the measured protein behaviors to gain a global view of protein interrelationships across IP targets and capture conditions. To achieve this we performed a multidimensional scaling (MDS) distance analysis and created an interactive 3D MDS plot (https://ncbps.shinyapps.io/3d_mds_app/; see Methods). Two representative snapshots of the plot are displayed in Figure 5. This analysis visually conveys the measured associations of the significantly different (p-value ≤ 0.05, log2 fold-change ≥ 1) proteins across the IP-MS experimental space: each experimental replicate contributes a dimension and MDS places each protein in a lower dimensional space that conserves distances between proteins as much as possible. In the present case, we had fifty-seven total dimensions before reduction, composed of twenty-three NCBP1-LAP IPs, twenty-two NCBP2-LAP IPs, and twelve NCBP3-LAP IPs. The output permitted us to compare and contrast the segregation of proteins that exhibited one or more statistically significant enrichments within the parameter space explored in the screen. The settings on the interactive plot allow users to hide or show different groups of proteins and also to calculate the MDS distance of proteins that passed from 1 to 12 statistical significance tests for co-enrichment from the full set of NCBP IPs.

**Figure 5:**
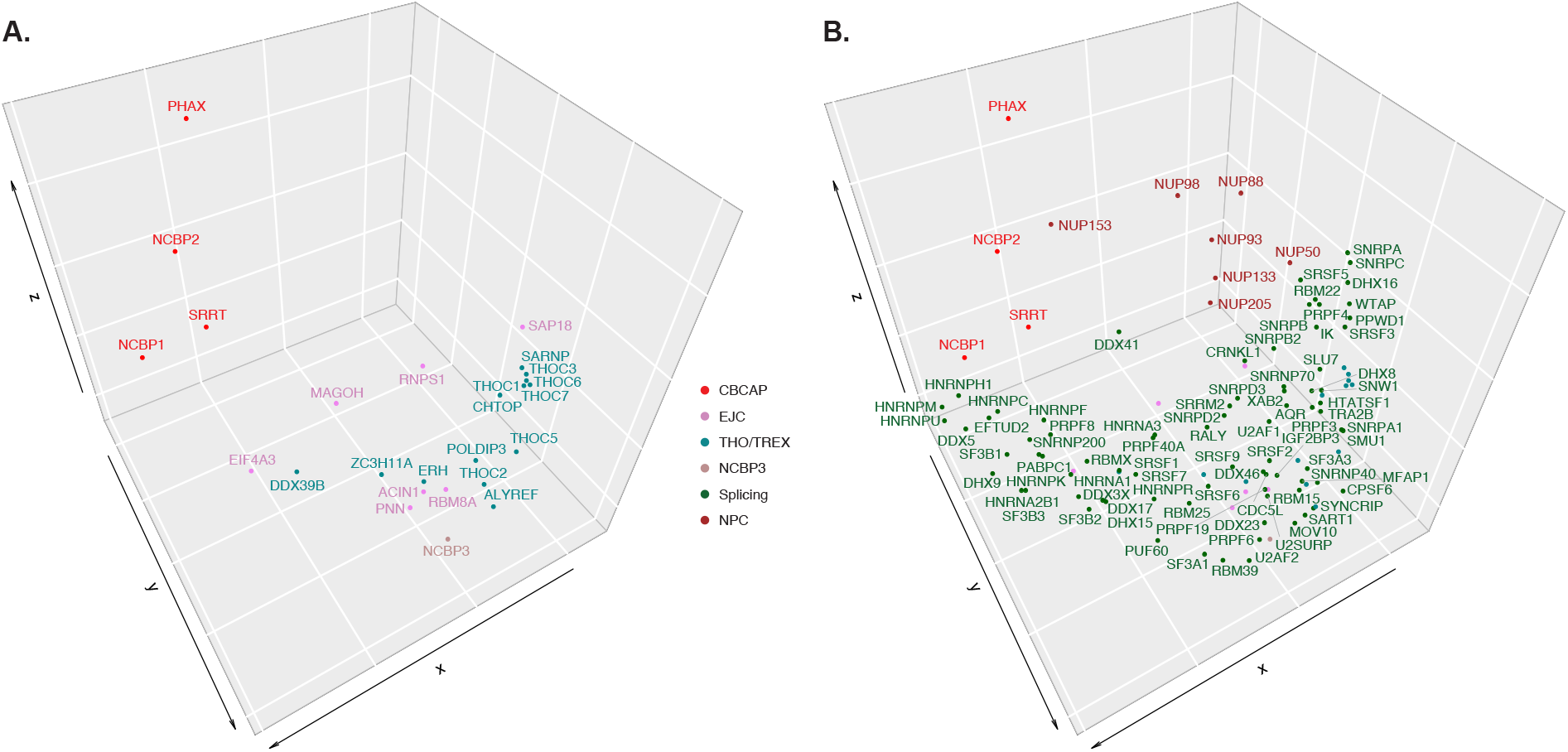
Global analysis contextualizes NCBP interactor relationships. **Multidimensional scaling (MDS) analysis of NCBP interactions.** Two orientations displaying select protein complexes from the 3D MDS are shown. See also, the full interactive plot: https://ncbps.shinyapps.io/3d_mds_app/. Panel (A) displays the segregation of the NCBPs, SRRT, PHAX, the EJC and TREX proteins. Panel (B) labels, CBCAP, spliceosomal, and NPC proteins with text; the nodes displayed for EJC and THO/TREX in panel (A) are also included in (B) for positional context, without labels to avoid crowding. These selected complexes can also be viewed in 3D context with rotation in the supplemental file: 3D_Animated_MDS.gif. The colour scheme for different groups of proteins is illustrated in the center of the plot.

Considering all IPs conducted in this study, only NCBP1/2 (CBC), SRRT, PHAX, KPNA3, and HNRNPUL2 passed our threshold for specific enrichment twelve separate times. CBC, SRRT, and PHAX are established interactors as part of the CBCA and CBCAP complexes (8–10). KPNA3 (importin subunit alpha-4) has been suggested to interact with both NCBP2 and NCBP3 (16). In our MDS analysis (see online plot), KPNA2, KPNA3 (importin-*α*) and KPNB1 (importin-*β*) cluster proximal to NCBP2 and NCBP1 (whereas NCBP3 clusters apart). The CBC-importin-*α* complex binds capped RNA in the nucleus, and the binding of importin-*β* stimulates the release of capped RNA from the CBC-importin-*α* complex in the cytoplasm (31, 35). We identified KPNA3, KPNA2, KPNB1, and RAN among these IP (see Figure 3B and Supplemental_heatmap.pdf), likely representing the different forms of CBC/RNP-importin associations. HNRNPUL2 has not previously been reported as an interactor of NCBPs, and very little information is available about this protein, but given these data, a functional role in coordination with CBC is likely.

Using the default settings (all groups, each protein must exhibit statistical significance for co-enrichment at least once) some observations appeared striking: (i) the CBC, SRRT, and PHAX segregate to a region of the graph that is otherwise relatively sparse of other nodes at a common extremity of the MDS plot. NCBP1/NCBP2/SRRT triangulate one another, with PHAX located on the opposite side of NCBP2 as NCBP1 and notably distal from the center of mass of the graph (Figure 5A); (ii) splicing proteins form an arch extending from the CBC towards the opposite extremity of the space, terminating near a cluster of nuclear pore complex (NPC) proteins (Figure 5B); and (iii) within this ‘splicing arch,’ located between the CBC and the NPC, EJC proteins interspersed with THO/TREX components; NCBP3 segregates nearly centrally within this EJC/THO/TREX spread (Figures 5A and B).

Considered directionally, these observations are remarkably consistent with the spatio-temporal features of the RNAPII-transcribed, mRNP maturation pathway. The binding of CBC-connected proteins represents one of the earliest steps in RNP assembly on 5’-capped, RNAPII transcripts (Figure 5A). The CBC proteins present at an interface with splicing proteins, a subsequent step in mRNP maturation (that typically occurs multiple times) - those most proximal to NCBP1 include HNRNPH1, HNRNPM, HNRNPU, HNRNPC, DDX5, and EFTUD2, followed by a myriad of other splicing proteins, spreading across a large, arching space (Figure 5B). This spread of proteins encompasses a collection of biochemically diverse and temporally staggered interactions, of which splicing is known to consist. The proteins we observed constitute an incomplete, cross-sectional composite of up to 79 out of 143 splicing factors curated by merging the CORUM categories ‘spliceosome C complex’ and ‘spliceosome’ - over 170 proteins have otherwise been classified as human spliceosome components (36–41). Towards the middle and end of the region demarcated by splicing proteins, EJC and TREX proteins appear next (text labels in Fig. 5A, overlap with splicing proteins in Fig. 5B); these are the proteins deposited at exon-exon junctions after splicing or shepherd mRNPs from transcription to export. First EIF4A3 and DDX39B appear; then MAGOH, which partitions adjacent to the splicing arch, roughly parallel to ZC3H11A, which lies within the arch; followed by ACIN1, PNN, ERH, NCBP3, and RBM8A partitioned in relative proximity within the crook of the arch, NCBP3 approximately at its apex. These are followed, among other EJC/TREX-linked proteins, by multiple THO complex components (also contributing to mRNP maturation and export; text labels in Fig. 5A, overlap with splicing proteins in Fig. 5B), reaching a terminal cluster along with various splicing factors. At this end of the arch, and opposite from the CBC, lies a cluster of NPC proteins (NUP50, NUP88, NUP93, NUP98, NUP133, NUP205; Fig. 5B), likely representing interactions exporting mRNPs from the nucleus. An exception to this is NUP153, which segregates closer to the opposite side of the arch (Fig. 5B): we speculate that this dichotomy reflects import of the CBC through the NPC, via docking to the nuclear basket protein Nup153 (42) (karyopherin / importin also cluster nearby, described above), and the final stages of nuclear export through the NPC via Nup98 (43, 44) and the cytoplasmic export platform that includes Nup88 (45).

## 3 Discussion

### 3.1 Interactome mapping by IP-MS

Among the most difficult challenges to interactome mapping by co-IP is the parameterization of optimal working conditions, enabling the transfer of target macromolecular complexes out of living cells and into the test tube. This is made more difficult by the fact that, while the target protein is known in advance, the macromolecular assemblies it forms in the cell are rarely known; and are highly unlikely to be known comprehensively, given the present state of interactome incompleteness (46, 47). When attempting to IP macromolecular complexes, the sample workup must afford at least the following two features to ensure the results will be informative: (i) the maintenance of at least some physiologically relevant macromolecular configurations and (ii) the simultaneous mitigation of accumulating post-lysis artifacts. Parameter optimization by parallelized IP screening confers access to these two features, representing a best practice that provides quality assurance to IP-MS studies (23). In the present study, we chose to apply this approach to chart NCBP interactomes (Figure 1A); these proteins join complex, dynamical RNPs in the cell, presenting a challenging use case. Upon initial screening, we found that the main differences between NCBP IPs were not sufficiently interpretable through the lens of SDS-PAGE / general protein staining alone. Although MS is most often used as an endpoint readout to verify sample composition, in this study we show that when SDS-PAGE protein banding patterns are not highly informative, MS, in conjunction with PAGE, is warranted for quality assurance and parameter selection (Figure 2A). Quantitative analysis using MS, along with appropriate controls, across the selected parameters can then be carried out with confidence (Figures 1B, 3, 4, and 5).

We chose to apply a popular LFQ MS based analysis using MaxQuant and custom post-processing in R (Figure 2B). Although our target proteins were ectopically expressed from within the host cell genome with a LAP-tag, we chose to use LFQ in-part because it is applicable to native biological sources where genomic tagging and/or metabolic labeling are not possible - modifying only the choice of control from what was used here. LFQ is also inexpensive but does come with some well-known shortcomings; one of which is the failure to detect common peptides (and thus proteins) between all samples analyzed. Although missing values are an intrinsic property of data dependent LFQ MS, it is notably problematic for the proteins that are enriched during the IP and not present in the cognate control. In a well-optimized IP, many (or most) of the proteins that co-IP with the experimental sample may not be detectably present in the control (24); but because proteins with missing values in the control cannot be expressed in terms of the commonly utilized metric ‘log2 fold-change’, imputation of small values to replace the missing values is a frequently used solution (e.g. reviewed in (48–50) and implemented in popular proteome analysis software). However, performance among imputation approaches varies. To improve the performance in our application, we developed a novel approach which takes advantage of information provided by the sample replicates (see Supplemental Methods, Section 5). Our algorithm resembles a decision tree that applies imputation in different ways depending on the degree to which it is needed. It outperformed the default imputation approach used in the popular Perseus software on the data produced in this study. Because imputation may suffer from instance-specific deficiencies, further trial and error testing is needed to determine the broader utility of our algorithm; based on its performance in this study we believe it holds great promise (we also applied it successfully in another LFQ MS study (51)).

### 3.2 NCBP interactome differences

With a well-performing bioinformatics pipeline in hand, we sought to mine the NCBP interactomes for missing information, potentially adding to our understanding of the macromolecules they form together and/or apart. We achieved this using three main visualizations: complex enrichment plots (Figure 3A), heatmaps (Figure 3B), and MDS (Figure 5). These analyses revealed highly similar protein associations exhibited by both NCBP1 and NCBP2, but also support possible moonlighting (52). NCBP1 IPs highly enriched components from the SMN complex, especially the SMN-independent intermediate containing GEMIN5, GEMIN4, and DDX20 (GEMIN3), but such enrichment was not observed in NCBP2 IPs, suggesting a stronger linkage of NCBP1 than NCBP2 to the snRNP maturation process (Figure 3). We also noticed some DNA-binding complexes preferentially co-enriched with NCBP2 (albeit only captured in one IP condition); these included the Ku-containing and PSF-p54-containing complexes (Figure 3 and https://ncbps.shinyapps.io/complex_enrichment/). This result may implicate NCBP2 in chromatin-associations that are distinct from NCBP1. There is scientific precedent for independent functional roles of NCBP1 and NCBP2: it has been shown that NCBP1 is expressed with roughly three-fold higher abundance than NCBP2 (16), supporting the hypothesis of some independent functions (52). Adding to that: individual depletion of NCBP1 and NCBP2 results in different effects on RNA export (16); in yeast, genetic deletion mutants of Cbp80p (NCBP1 homolog) and Cbp20p (NCBP2 homolog), respectively, share fewer than half of the resulting gene expression-level changes in common (55); and NCBP1, but not NCBP2, was co-enriched with eIF4E-bound transcripts (56), establishing NCBP2-independent, NCBP1-associated RNPs.

NCBP3 has been proposed capable of substituting for NCBP2 in the CBC and suppressing mRNA export defects caused by NCBP2 loss (16). Our findings do not contradict that proposal, but they show that NCBP1, NCBP2, and NCBP3 may normally all be present simultaneously within the same population of isolated RNPs - NCBP2 can co-IP NCBP3 and NCBP3 can co-IP NCBP2 (Figure 3B). If the existing cap-binding paradigm holds, then within mRNPs, NCBP3 may sometimes bind NCBP1/2 (canonical CBC) - fostering crosstalk with other mRNP maturation factors (discussed below). NCBP1 and NCBP2 are thought to be present in a 1:1 stoichiometry in the canonical CBC; yet we observed that NCBP1 may be present in apparent excess to NCBP2 (Figure 4) - importantly, this was also observed in IPs targeting NCBP2. This raises the possibility of multiple copies of NBCP1 in some NCBP1/2 containing macromolecules. Our current findings are not conclusive and warrant deeper follow-up analysis, including more precise and accurate quantitation by alternative methods. But if this observation holds true, our co-IPs of NCBP3 with NCBP2 and NCBP2 with NCBP3 may signify a mixture of interactions: NCBP1 and NCBP2, together in the context of the CBC, and NCBP1 and NCBP3 together in a distinct context, possibly more distal from the 5’-cap - the latter scenario would naturally increase the NCBP1:NCBP2 ratio in some mRNPs. We consider this possibility to be bolstered by knowledge of NCBP1s excess expression to NCBP2 and potential to engage RNPs apart from NCBP2 (16, 56).

Before NCBP3 was proposed to bind 5’-caps, its associations with mRNPs were instead linked to splicing and grouped with the EJC and TREX (17–20). Our studies reinforce these associations (Figure 3), placing NCBP3 at an interface between splicing, EJC, and TREX based on MDS analysis (Figure 5). All the evidence considered together, a parsimonious conjecture is that when NCBP3 is present, it (primarily) associates with EJC/TREX and may also (secondarily) interact with CBC via NCBP1 (when PHAX is not present). Notably, NCBP1 exhibits a somewhat intermediate level of EJC enrichment compared to NCBP2 (low EJC) and NCBP3 (high EJC), while NCBP3 associations are also strongly skewed toward THO/TREX.

### 3.3 MDS reconstructs ordered pathways from heterogeneous interactions

The MDS analysis provided key evidence to encourage our speculations regarding possible CBC and NCBP3 macromolecular organization. Our implementation is agnostic to how many times a protein was statistically significant, as long as it was significant at least once (statistical stringency is user selectable in the interactive MDS, https://ncbps.shinyapps.io/3d_mds_app/). This decision was made using the following rationale: in this study, we first formed our belief in a collection of in vivo interactions based on evidence, obtained from across the breadth of our screen, in the form of t-tests comparing cases and controls after IP-MS analysis. Thus having a large set of physiologically believable interactions, we then focussed on their in vitro behavioral patterns holistically. In a recent study, we showed that the co-partitioning of proteins across numerous IP-MS experiments revealed known and novel physical and function relationships, providing a basis for algorithmic clustering of putative macromolecules from heterogeneous mixtures (57). Although the affinity proteomics and algorithm design used here were distinctive in several details compared to the referenced prior study, comparable concepts apply. In short, it is informative for us to observe the co-behavior of all proteins that interact specifically, even when, in a given experiment they are not shown to be specific in that test tube.

We contend that the clustering of protein behaviors in our MDS plot provides the basis for a ‘pseudo-timeline’ of protein interactions along enriched pathways. MDS and other dimensionality-reduction techniques are already being used for conceptually comparable purposes e.g. in reconstructing the temporal progressions of cell states from transcriptional signatures (58–60). As described in the results section, the MDS graph (Figure 5) revealed a conspicuous and reassuring assortment of proteins that cluster together with other functionally-like proteins; the arrangement of the protein clusters related to mRNP maturation also occur in a manner that recapitulates the accepted composition and order of assembly of the mRNP pathway (61, 62). Several striking observations follow: (i) The CBC, SRRT, and PHAX segregate to a region of the graph that is otherwise sparse of other nodes at a common extremity of the MDS plot, with PHAX notably distal from the center of mass of the graph - this distance may be rationalized in light of our panel of results being enriched for mRNP processing factors - whereas PHAX connects with sn/snoRNP processing (12, 13); conducting PHAX IPs should contribute a new segment to the plot in this sparsely occupied space; (ii) Splicing proteins form an arch extending from the CBC towards the opposite extremity of the space, terminating in a cluster of NPC proteins; this is satisfyingly consistent with the reconstruction of a pseudo-timeline spanning cap-binding through processing and nuclear export of mRNAs; (iii) within the ‘splicing arch’, located between the CBC and NPC, EJC proteins interspersed with THO/TREX components appear; NCBP3 segregates nearly centrally within this EJC/THO/TREX spread; supporting the prior art and our contention that NCBP3 is, first and foremost, a physical and functional component of these complexes, rather than the CBC. If NCBP3 joins mRNPs after the canonical CBC has already formed, then the in vivo relevance of NCBP3’s ability to bind m7GTP in vitro (5) remains open for clarification (this binding will have already been satisfied by the CBC). One relevant scenario may be when NCBP3 substitutes for NCBP2, forming an alternative CBC: this may occur only (or primarily) upon NCBP2 loss or down regulation. If so, this activity may be mechanistically distinct from the splicing and mRNP maturation-connected NCBP3 narrative described by others (17–20), and reinforced based on the interactions observed here.

To avoid over-interpreting these data at an unjustified granularity, we are treating the result as largely descriptive. What is clear is that the interrelationships we observed broadly follow the interaction sequence and functional rationale believed to apply in vivo; it is notable that comparable though increasingly less complete assortment patterns of protein classes are discernible in the MDS plot when restricting the proteins analyzed only to those that have passed up to 4 t-tests (above which the number of nodes are few and the pattern begins to degrade). That said, these data emanate from a screen that explores macromolecule stability via their differing compositions, obtained under a multitude of biochemical challenges - the relative abundances of the proteins offers an opportunity for quantitative readout of their affinities, given experimental conditions. Interactomics as a discipline has shown it possible to authentically recapitulate macromolecular compositions by sampling a large number of target proteins by IP-MS (e.g. (63, 64)) - but the community also understands that there is still a great deal of missing information (46, 47). We contend that this information is missing, in large part, because the conditions of IP-MS are not optimized for the target macromolecules; as a result, in vivo interactions are rapidly shed during extraction and thus elude detection (23) - the present study extends the tools at our disposal to combat missing information.

## 4 Experimental Procedures

### 4.1 Preparation HeLa cells expressing LAP-tagged NCBPs and affinity capture

HeLa Kyoto cell lines stably expressing LAP-tagged proteins (NCBP1, 2, and 3) and control, “tag-only” (LAP-control), respectively, were provided as stably transfected cell pools by Ina Poser and Anthony Hyman (see Supplemental Methods, Section 1); these were engineered as previously described (65, 66), containing a TEV cleavage site, S peptides, a PreScission protease cleavage site and EGFP. For C-terminally tagged NCBPs, the LAP-tag DNA sequence is positioned in front of the stop codon; and for the LAP-control cells, the LAP tag DNA sequence is placed under control of the TUBG1 promoter (65). All cell pools provided were FACS sorted for EGFP-positive cells (forward scatter threshold = 5000). The sorted cell pools were cultured using standard techniques in DMEM supplemented with FBS and penicillin-streptomycin. Cell harvesting and cryomilling was carried out as previously described (24). 50 mg of cell powder (wet cell weight equivalent) was used per well (NCBP1, 2 and cognate controls) at 1:9 (w:v) in 24-wells of a 96-well plate; 300 mg of cell powder was used per affinity capture replicate of NCBP3 and cognate controls at 1:4 (w:v) when conducted individually in microfuge tubes. Screens were conducted as in (23), with modifications described in this study. Sonication was achieved using a QSonica Q700 equipped with an 8-tip microprobe (#4602), applied until material in multi-well plates was homogeneously resuspended as judged by visual inspection (4°C, 1 Amp, 30-40 sec [continuous]: 140 J per row on average); or using a QSonica S4000 equipped with a low-energy microprobe (#4717), applied individually in microfuge tubes (4°C, 2 Amp, 15 × 2 sec pulses [1 sec interval]: 50 J per sample). After centrifugal clarification of the extracts (10 min, 4°C, 20,000 RCF), affinity capture was achieved using 5 *μ*l of affinity medium slurry, conjugated with Llama *α*-GFP polyclonal antibody (67, 68), in multi-well screens, or 10 *μ*l of slurry in microfuge tubes. Affinity capture was allowed to proceed for 30 min at 4 °C with gentle mixing. Elution from the affinity medium was achieved using 1.1x LDS (ThermoFisher Scientific #NP0008). Protein extraction solutions used and obtained SDS-PAGE, protein staining results are presented in the file: Supplemental_Data_Tables.xlxs, and are also curated in an interactive, searchable form on http://copurification.org/.

### 4.2 IP-MS experimental design and data processing

The experimental schema is depicted in Figure 1B. 24 unique solutions were used for affinity capture pre-screening of NCBP1-LAP. We performed hierarchical clustering of the protein LFQ intensities obtained (Figure 2A); missing values were set to 0. To preserve a diverse parameter space for screening while freeing up bandwidth for replicates, we selected 6 extraction and capture solutions from across the breadth of the dendrogram while accounting for the quality of gel profiles (by visual inspection) and proteinaceous complexity (MS-based). These were used in quadruplicate IP-MS experiments for all affinity capture targets (using 24-wells of 96-well plates). Of these, at least three replicates passed initial gel-based quality control (QC) by visual inspection: those lanes exhibiting a relatively discrete pattern of sharply stained bands, a relative paucity of faint/fuzzy background staining, and a band that apparently corresponds to the target protein molecular mass, were moved forward to quantitative MS analyses. Subsequent computational QC of the MS data was applied using the R package PTXQC and in-house R scripts (see Data processing, below); NCBP3-LAP multi-well format IP-MS data did not pass, and was omitted (see Supplemental Methods, Section 2); poor quality was attributed primarily to low yield for this target protein at the 50 mg-scale used for multi-well IPs. To obtain usable IP-MS data for this target, the scale was increased to 300 mg and repeated in microfuge tubes, in four extraction and capture solutions and three replicates each. For multi-well and microfuge tube experiments, NCBP and LAP-control targets were captured under identical conditions, respectively. Accounting of all the samples and replicates used in this study is provided in the file: Supplemental_Data_Tables.xlxs. Statistical methods are described in detail, below.

### 4.3 Sample workup and mass spectrometry

Samples were reduced (DTT) and alkylated (iodoacetamide), and a fraction of the sample was subjected to standard SDS-PAGE, staining with Sypro Ruby, and CCD imaging (Fuji LAS-4000); the other fraction was run as a gel plug, Coomassie Blue stained, excised, subjected to in-gel tryptic digestion, and the peptides desalted and concentrated upon C18 resin (OMIX C18 pipette tips; Agilent #A57003100) essentially as previously described (69). For multi-well screens, 1/2 of the sample was used for imaging and MS respectively. For NCBP3 IPs done in microfuge tubes, 1/6 of the sample was used for imaging and the rest for MS. Standard SDS-PAGE was used for initial QC of sample composition and excision of select bands (based on imaging) for protein identification by tandem mass spectrometry; gel plugs were used to prepare whole IP fractions for LFQ MS. Summarized as follows: Samples produced by multi-well screening were run on either an Orbitrap Fusion or a Q Exactive Plus (Thermo Fisher Scientific). Dried peptide samples were resuspended in 10 *μ*L of 5% (v/v) methanol, 0.2% (v/v) formic acid in water; half was loaded on the LC column (Thermo Easy-Spray ES800). Peptides were ionized by electrospray at 1.8 – 2.1 kV following elution across a linear gradient (7 minutes for individual gel bands; 35-40 minutes for whole gel plugs) rising to 30% (v/v) acetonitrile. Solvent A was 0.1% (v/v) formic acid in water; Solvent B was 0.1% (v/v) formic acid prepared by combination with either 95% (v/v) or neat acetonitrile. Full MS scans were performed in profile mode, while fragmentation spectra were acquired in centroid mode with priority given to the most intense precursors. Dynamic exclusion was enabled to limit repeated sequencing of the same peptides. For NCBP3-LAP and control samples obtained by microfuge tube affinity capture: the dried peptide mix was reconstituted in a solution of 20 *μ*l of 2% (v/v) formic acid (FA) for MS analysis. 5 *μ*l of this solution was loaded with the autosampler directly onto a self-packed column, which was made from a 75 *μ*m ID PicoFrit column (New Objective, Woburn, MA) filled with 25 cm of 2.4 *μ*m Reprosil-Pur C18 AQ. Peptides were eluted at 200 nl/min from the column using an Eksigent NanoLC 415 with a 52 min gradient from 2% to 25% buffer B (0.1% (v/v) formic acid in acetonitrile); at which point the gradient was switched from 25% to 85% buffer B over 5 min and held constant for 3 min; finally, the gradient was changed from 85% buffer B to 98% buffer A (0.1% (v/v) formic acid in water) over 1 min, and then held constant at 98% buffer A for 15 more minutes. The application of a 3.5 kV distal voltage electrosprayed the eluting peptides directly into a Q Exactive HF mass spectrometer equipped with an EASY-Spray source (Thermo Scientific). Mass spectrometer-scanning functions and HPLC gradients were controlled by the Xcalibur data system (Thermo Scientific). The mass spectrometer was set to scan MS1 at 60,000 resolution with an AGC target set at 3×106. The scan range was m/z 375-2000. For MS2, resolution was set at 15,000 and AGC target at 2×105 with a maximum IT at 50 ms. The top 15 peaks were analyzed by MS2. Peptides were isolated with an isolation window of m/z 1.6 and fragmented at 27 CE. Minimum AGC target was at 8×103. Only ions with a charge state of 2 through 6 were considered for MS2. Dynamic exclusion was set at 15 sec. See Supplemental Methods, Section 3 for an ordered summary of instrument settings.

### 4.4 Data processing

Our computational proteomic pipeline is summarized in Figure 2B; supplementary information, figures, and data tables are indicated. Pre-processing of raw data was done in MaxQuant; data post processing, including statistical filtering and distance calculations, were done in R - all code is available at https://github.com/moghbaie/NCBP-pipeline. Peptide identification and quantitation was achieved using the MaxQuant v.1.6.5.0 software and a proteomic database comprised of a Uniprot human proteome (proteome:up000005640; reviewed:yes) with GFP added (97% identical to the EGFP sequence in the LAP-tag). The following abridged software settings were used: Include contaminants - True; PSM & Protein FDR - 0.01; quantify unmodified peptides and Oxidation (M), Acetyl (Protein N-term), Carbamidomethyl (C); Phospho (STY) was searched but not quantified, nor were unmodified counterpart peptides; iBAQ - True; iBAQ log fit - False; Match between runs - True; Decoy mode - revert; Include contaminants - True; Advanced ratios - False; Second peptides - True; Stabilize large LFQ ratios - True; Separate LFQ in parameter groups - True; Require MS/MS for LFQ comparisons - True; Razor protein FDR - True. The RAW and MaxQuant v.1.6.5.0 processed files are available for download via ProteomeXchange with identifier PXD016038. Data quality was initially assessed using the output from MaxQuant with the R package PTXQC (69) and in-house R scripts - examples detailed in Supplemental Methods, Section 2. Proteins marked as “contaminants” or “reverse” by MaxQuant were removed and intensities of the proteins MAGOH (UniProt ID P61326) and MAGOHB (UniProt ID Q96A72) were summed together in every experiment (see Supplemental Methods, Section 4). Only proteins which had “Peptide counts (razor+unique)” ≥ 2 were retained for analysis. Using the data obtained, protein intensities were log2-transformed in order to sample from a normal distribution during imputation (see Supplemental Methods, Section 5 for a full description with performance testing and results). Imputation for LFQ and iBAQ intensities was repeated 100 times (see below). To compare protein LFQ or iBAQ intensities between different experiments, they were normalized by the values for GFP obtained from within the same experiment (see Supplemental Methods, Section 6). For statistical testing, the intensities were rescaled by multiplying all values by a coefficient equal to the mean of the smallest and the largest intensity value before the normalization, restoring the original range of the data, log2-transformed intensities were used for further analysis.

Proteins were subjected to unpaired, two-sample t-tests between LAP-tagged-targets and LAP-only-controls for each set of IPs. Protein enrichment with the LAP-tagged NCBP target in each experiment was considered statistically significant if the Benjamini–Hochberg adjusted p-value was ≤ 0.05 and log2 transformed fold-change (log2FC; target/control) was ≥ 1. To mitigate imputation-induced sampling artifacts, only proteins that reached significance in ≥ 60 (of 100) imputation trials were retained for further consideration, and imputation induced false positives were removed (see Supplement Methods, Section 6). Next, all proteins reaching significance in at least one experiment were collated: these values were used in quantitative analyses. For every protein reaching statistical significance for co-enrichment with an NCBP target at least once, pairwise euclidean distances were calculated. For Figure 3 (complex enrichment plot and heatmap), background subtracted LFQ intensities were used to eliminate contributions by non-specific interactions. This was achieved by subtracting the mean LFQ intensities of proteins in control experiments from the LFQ intensities in the NCBP IPs, after which, the values were log10-transformed; the subtraction was applied to each experimental condition separately.

## 5 Availability of data and materials

The MS proteomics data have been deposited to the ProteomeXchange Consortium via the PRIDE (70) partner repository with the dataset identifier PXD016038. R code available at https://github.com/moghbaie/NCBP-pipeline.

## 6 Acknowledgments

Research reported in this publication was supported in-part by the National Institute of General Medical Sciences of the National Institutes of Health (NIH) including: grant R01GM126170 (to JL), grant P41GM103314 (to BTC), and grant P41GM109824 (National Center for Dynamic Interactome Research, to MPR.). The content is solely the responsibility of the authors and does not necessarily represent the official views of the National Institutes of Health. This research was also supported by the Independent Research Fund Denmark, the Novo Nordisk Foundation and the Lundbeck Foundation to THJ. We thank Drs. Emily Chen and Toni Koller for collecting NCBP3-LAP MS data, and Dr. Ina Poser and Prof. Tony Hyman for providing the LAP-tagged cell lines.

## 7 Author contributions

Yuhui Dou - methodology, validation, investigation, writing (original draft preparation, review and editing), visualization; Svetlana Kalmykova - software, formal analysis, data curation, writing (original draft preparation, review and editing), visualization; Maria Pashkova - software, formal analysis, data curation, writing (original draft preparation, review and editing), visualization; Mehrnoosh Oghbaie - methodology, software, formal analysis, data curation, visualization; Hua Jiang - investigation; Kelly R. Molloy - investigation; Brian T. Chait - resources, funding acquisition; Michael P. Rout - resources, funding acquisition, writing (review and editing), visualization; David Fenyö - supervision; Torben Heick Jensen - conceptualization, funding acquisition, writing (review and editing), supervision; Ilya Altukhov - methodology, writing (original draft preparation, review and editing), supervision, project administration; John LaCava - conceptualization, methodology, resources, data curation, writing (original draft preparation, review and editing), visualization, supervision, project administration, funding acquisition.

## Notes

### Competing Interest Statement

The authors have declared no competing interest.

## References

1. Shatkin,A.J. (1976) Capping of eucaryotic mRNAs. Cell, 9, 645–653.

2. Salditt-Georgieff,M., Harpold,M., Chen-Kiang,S. and Darnell,J.E. (1980) The addition of 5’ cap structures occurs early in hnRNA synthesis and prematurely terminated molecules are capped. Cell, 19, 69–78.

3. Izaurralde,E., Lewis,J., McGuigan,C., Jankowska,M., Darzynkiewicz,E. and Mattaj,I.W. (1994) A nuclear cap binding protein complex involved in pre-mRNA splicing. Cell, 78, 657–668.

4. Mazza,C., Ohno,M., Segref,A., Mattaj,I.W. and Cusack,S. (2001) Crystal structure of the human nuclear cap binding complex. Mol. Cell, 8, 383–396.

5. Schulze,W.M., Stein,F., Rettel,M., Nanao,M. and Cusack,S. (2018) Structural analysis of human ARS2 as a platform for co-transcriptional RNA sorting. Nat. Commun., 9, 1701.

6. Lenasi,T., Peterlin,B.M. and Barboric,M. (2011) Cap-binding protein complex links pre-mRNA capping to transcription elongation and alternative splicing through positive transcription elongation factor b (P-TEFb). J. Biol. Chem., 286, 22758–22768.

7. Pabis,M., Neufeld,N., Steiner,M.C., Bojic,T., Shav-Tal,Y. and Neugebauer,K.M. (2013) The nuclear cap-binding complex interacts with the U4/U6 U5 tri-snRNP and promotes spliceosome assembly in mammalian cells. RNA, 19, 1054–1063.

8. Giacometti,S., Benbahouche,N.E.H., Domanski,M., Robert,M.-C., Meola,N., Lubas,M., Bukenborg,J., Andersen,J.S., Schulze,W.M., Verheggen,C., et al. (2017) Mutually Exclusive CBC-Containing Complexes Contribute to RNA Fate. Cell Rep., 18, 2635–2650.

9. Hallais,M., Pontvianne,F., Andersen,P.R., Clerici,M., Lener,D., Benbahouche,N.E.H., Gostan,T., Vandermoere,F., Robert,M.-C., Cusack,S., et al. (2013) CBC-ARS2 stimulates 3’-end maturation of multiple RNA families and favors cap-proximal processing. Nat. Struct. Mol. Biol., 20, 1358–1366.

10. Andersen,P.R., Domanski,M., Kristiansen,M.S., Storvall,H., Ntini,E., Verheggen,C., Schein,A., Bunkenborg,J., Poser,I., Hallais,M., et al. (2013) The human cap-binding complex is functionally connected to the nuclear RNA exosome. Nat. Struct. Mol. Biol., 20, 1367–1376.

11. Meola,N., Domanski,M., Karadoulama,E., Chen,Y., Gentil,C., Pultz,D., Vitting-Seerup,K., Lykke-Andersen,S., Andersen,J.S., Sandelin,A., et al. (2016) Identification of a nuclear exosome decay pathway for processed transcripts. Mol. Cell, 64, 520–533.

12. Boulon,S., Verheggen,C., Jady,B.E., Girard,C., Pescia,C., Paul,C., Ospina,J.K., Kiss,T., Matera,A.G., Bordonné,R., et al. (2004) PHAX and CRM1 are required sequentially to transport U3 snoRNA to nucleoli. Mol. Cell, 16, 777–787.

13. Ohno,M., Segref,A., Bachi,A., Wilm,M. and Mattaj,I.W. (2000) PHAX, a mediator of U snRNA nuclear export whose activity is regulated by phosphorylation. Cell, 101, 187–198.

14. Izaurralde,E., Lewis,J., Gamberi,C., Jarmolowski,A., McGuigan,C. and Mattaj,I.W. (1995) A cap-binding protein complex mediating U snRNA export. Nature, 376, 709–712.

15. Cheng,H., Dufu,K., Lee,C.-S., Hsu,J.L., Dias,A. and Reed,R. (2006) Human mRNA export machinery recruited to the 5’ end of mRNA. Cell, 127, 1389–1400.

16. Gebhardt,A., Habjan,M., Benda,C., Meiler,A., Haas,D.A., Hein,M.Y., Mann,A., Mann,M., Habermann,B. and Pichlmair,A. (2015) mRNA export through an additional cap-binding complex consisting of NCBP1 and NCBP3. Nat. Commun., 6, 8192.

17. Merz,C., Urlaub,H., Will,C.L. and Lührmann,R. (2007) Protein composition of human mRNPs spliced in vitro and differential requirements for mRNP protein recruitment. RNA, 13, 116–128.

18. Hegele,A., Kamburov,A., Grossmann,A., Sourlis,C., Wowro,S., Weimann,M., Will,C.L., Pena,V., Lührmann,R. and Stelzl,U. (2012) Dynamic protein-protein interaction wiring of the human spliceosome. Mol. Cell, 45, 567–580.

19. Fan,J., Kuai,B., Wu,G., Wu,X., Chi,B., Wang,L., Wang,K., Shi,Z., Zhang,H., Chen,S., et al. (2017) Exosome cofactor hMTR4 competes with export adaptor ALYREF to ensure balanced nuclear RNA pools for degradation and export. EMBO J., 36, 2870–2886.

20. Dufu,K., Livingstone,M.J., Seebacher,J., Gygi,S.P., Wilson,S.A. and Reed,R. (2010) ATP is required for interactions between UAP56 and two conserved mRNA export proteins, Aly and CIP29, to assemble the TREX complex. Genes Dev., 24, 2043–2053.

21. Morris,J.H., Knudsen,G.M., Verschueren,E., Johnson,J.R., Cimermancic,P., Greninger,A.L. and Pico,A.R. (2014) Affinity purification-mass spectrometry and network analysis to understand protein-protein interactions. Nat. Protoc., 9, 2539–2554.

22. LaCava,J., Molloy,K.R., Taylor,M.S., Domanski,M., Chait,B.T. and Rout,M.P. (2015) Affinity proteomics to study endogenous protein complexes: pointers, pitfalls, preferences and perspectives. BioTechniques, 58, 103–119.

23. Hakhverdyan,Z., Domanski,M., Hough,L.E., Oroskar,A.A., Oroskar,A.R., Keegan,S., Dilworth,D.J., Molloy,K.R., Sherman,V., Aitchison,J.D., et al. (2015) Rapid, optimized interactomic screening. Nat. Methods, 12, 553–560.

24. LaCava,J., Jiang,H. and Rout,M.P. (2016) Protein Complex Affinity Capture from Cryomilled Mammalian Cells. J. Vis. Exp., 10.3791/54518.

25. Jancarik,J. and Kim,S.H. (1991) Sparse matrix sampling: a screening method for crystallization of proteins. J. Appl. Crystallogr., 24, 409–411.

26. Armean,I.M., Lilley,K.S. and Trotter,M.W.B. (2013) Popular computational methods to assess multiprotein complexes derived from label-free affinity purification and mass spectrometry (AP-MS) experiments. Mol. Cell. Proteomics, 12, 1–13.

27. Ma,Y., Silveri,L., LaCava,J. and Dokudovskaya,S. (2017) Tumor suppressor NPRL2 induces ROS production and DNA damage response. Sci. Rep., 7, 15311.

28. Winczura,K., Schmid,M., Iasillo,C., Molloy,K.R., Harder,L.M., Andersen,J.S., LaCava,J. and Jensen,T.H. (2018) Characterizing ZC3H18, a Multi-domain Protein at the Interface of RNA Production and Destruction Decisions. Cell Rep., 22, 44–58.

29. Giurgiu,M., Reinhard,J., Brauner,B., Dunger-Kaltenbach,I., Fobo,G., Frishman,G., Montrone,C. and Ruepp,A. (2019) CORUM: the comprehensive resource of mammalian protein complexes-2019. Nucleic Acids Res., 47, D559–D563.

30. Narita,T., Yung,T.M.C., Yamamoto,J., Tsuboi,Y., Tanabe,H., Tanaka,K., Yamaguchi,Y. and Handa,H. (2007) NELF interacts with CBC and participates in 3’ end processing of replication-dependent histone mRNAs. Mol. Cell, 26, 349–365.

31. Dias,S.M.G., Wilson,K.F., Rojas,K.S., Ambrosio,A.L.B. and Cerione,R.A. (2009) The molecular basis for the regulation of the cap-binding complex by the importins. Nat. Struct. Mol. Biol., 16, 930–937.

32. Chari,A., Golas,M.M., Klingenhäger,M., Neuenkirchen,N., Sander,B., Englbrecht,C., Sickmann,A., Stark,H. and Fischer,U. (2008) An assembly chaperone collaborates with the SMN complex to generate spliceosomal SnRNPs. Cell, 135, 497–509.

33. Li,B., Navarro,S., Kasahara,N. and Comai,L. (2004) Identification and biochemical characterization of a Werner’s syndrome protein complex with Ku70/80 and poly(ADP-ribose) polymerase-1. J. Biol. Chem., 279, 13659–13667.

34. Smits,A.H., Jansen,P.W.T.C., Poser,I., Hyman,A.A. and Vermeulen,M. (2013) Stoichiometry of chromatin-associated protein complexes revealed by label-free quantitative mass spectrometry-based proteomics. Nucleic Acids Res., 41, e28.

35. Görlich,D., Kraft,R., Kostka,S., Vogel,F., Hartmann,E., Laskey,R.A., Mattaj,I.W. and Izaurralde,E. (1996) Importin provides a link between nuclear protein import and U snRNA export. Cell, 87, 21–32.

36. Will,C.L. and Lührmann,R. (2011) Spliceosome structure and function. Cold Spring Harb. Perspect. Biol., 3.

37. Bertram,K., Agafonov,D.E., Dybkov,O., Haselbach,D., Leelaram,M.N., Will,C.L., Urlaub,H., Kastner,B., Lührmann,R. and Stark,H. (2017) Cryo-EM Structure of a Pre-catalytic Human Spliceosome Primed for Activation. Cell, 170, 701–713.e11.

38. Bertram,K., Agafonov,D.E., Liu,W.-T., Dybkov,O., Will,C.L., Hartmuth,K., Urlaub,H., Kastner,B., Stark,H. and Lührmann,R. (2017) Cryo-EM structure of a human spliceosome activated for step 2 of splicing. Nature, 542, 318–323.

39. Zhang,X., Yan,C., Zhan,X., Li,L., Lei,J. and Shi,Y. (2018) Structure of the human activated spliceosome in three conformational states. Cell Res., 28, 307–322.

40. Zhan,X., Yan,C., Zhang,X., Lei,J. and Shi,Y. (2018) Structures of the human pre-catalytic spliceosome and its precursor spliceosome. Cell Res., 28, 1129–1140.

41. Zhan,X., Yan,C., Zhang,X., Lei,J. and Shi,Y. (2018) Structure of a human catalytic step I spliceosome. Science, 359, 537–545.

42. Makise,M., Mackay,D.R., Elgort,S., Shankaran,S.S., Adam,S.A. and Ullman,K.S. (2012) The Nup153-Nup50 protein interface and its role in nuclear import. J. Biol. Chem., 287, 38515–38522.

43. Griffis,E.R., Altan,N., Lippincott-Schwartz,J. and Powers,M.A. (2002) Nup98 is a mobile nucleoporin with transcription-dependent dynamics. Mol. Biol. Cell, 13, 1282–1297.

44. Blevins,M.B., Smith,A.M., Phillips,E.M. and Powers,M.A. (2003) Complex formation among the RNA export proteins Nup98, Rae1/Gle2, and TAP. J. Biol. Chem., 278, 20979–20988.

45. Fernandez-Martinez,J., Kim,S.J., Shi,Y., Upla,P., Pellarin,R., Gagnon,M., Chemmama,I.E., Wang,J., Nudelman,I., Zhang,W., et al. (2016) Structure and Function of the Nuclear Pore Complex Cytoplasmic mRNA Export Platform. Cell, 167, 1215–1228.e25.

46. Tompa,P. and Rose,G.D. (2011) The Levinthal paradox of the interactome. Protein Sci., 20, 2074–2079.

47. Menche,J., Sharma,A., Kitsak,M., Ghiassian,S.D., Vidal,M., Loscalzo,J. and Barabási,A.-L. (2015) Disease networks. Uncovering disease-disease relationships through the incomplete interactome. Science, 347, 1257601.

48. Webb-Robertson,B.-J.M., Wiberg,H.K., Matzke,M.M., Brown,J.N., Wang,J., McDermott,J.E., Smith,R.D., Rodland,K.D., Metz,T.O., Pounds,J.G., et al. (2015) Review, evaluation, and discussion of the challenges of missing value imputation for mass spectrometry-based label-free global proteomics. J. Proteome Res., 14, 1993–2001.

49. Välikangas,T., Suomi,T. and Elo,L.L. (2018) A comprehensive evaluation of popular proteomics software workflows for label-free proteome quantification and imputation. Brief. Bioinformatics, 19, 1344–1355.

50. Tyanova,S., Temu,T., Sinitcyn,P., Carlson,A., Hein,M.Y., Geiger,T., Mann,M. and Cox,J. (2016) The Perseus computational platform for comprehensive analysis of (prote)omics data. Nat. Methods, 13, 731–740.

51. Ardeljan,D., Wang,X., Oghbaie,M., Taylor,M.S., Husband,D., Deshpande,V., Steranka,J.P., Gorbounov,M., Yang,W.R., Sie,B., et al. (2020) LINE-1 ORF2p expression is nearly imperceptible in human cancers. Mob. DNA, 11, 1.

52. Gonatopoulos-Pournatzis,T. and Cowling,V.H. (2014) Cap-binding complex (CBC). Biochem. J., 457, 231–242.

53. Lee,C.-G., Hague,L.K., Li,H. and Donnelly,R. (2004) Identification of toposome, a novel multisubunit complex containing topoisomerase IIalpha. Cell Cycle, 3, 638–647.

54. Du,Y.-C., Gu,S., Zhou,J., Wang,T., Cai,H., Macinnes,M.A., Bradbury,E.M. and Chen,X. (2006) The dynamic alterations of H2AX complex during DNA repair detected by a proteomic approach reveal the critical roles of Ca(2+)/calmodulin in the ionizing radiation-induced cell cycle arrest. Mol. Cell. Proteomics, 5, 1033–1044.

55. Baron-Benhamou,J., Fortes,P., Inada,T., Preiss,T. and Hentze,M.W. (2003) The interaction of the cap-binding complex (CBC) with eIF4G is dispensable for translation in yeast. RNA, 9, 654–662.

56. Rufener,S.C. and Mühlemann,O. (2013) eIF4E-bound mRNPs are substrates for nonsense-mediated mRNA decay in mammalian cells. Nat. Struct. Mol. Biol., 20, 710–717.

57. Taylor,M.S., Altukhov,I., Molloy,K.R., Mita,P., Jiang,H., Adney,E.M., Wudzinska,A., Badri,S., Ischenko,D., Eng,G., et al. (2018) Dissection of affinity captured LINE-1 macromolecular complexes. elife, 7.

58. Kouno,T., de Hoon,M., Mar,J.C., Tomaru,Y., Kawano,M., Carninci,P., Suzuki,H., Hayashizaki,Y. and Shin,J.W. (2013) Temporal dynamics and transcriptional control using single-cell gene expression analysis. Genome Biol., 14, R118.

59. Trapnell,C., Cacchiarelli,D., Grimsby,J., Pokharel,P., Li,S., Morse,M., Lennon,N.J., Livak,K.J., Mikkelsen,T.S. and Rinn,J.L. (2014) The dynamics and regulators of cell fate decisions are revealed by pseudotemporal ordering of single cells. Nat. Biotechnol., 32, 381–386.

60. Ji,Z. and Ji,H. (2016) TSCAN: Pseudo-time reconstruction and evaluation in single-cell RNA-seq analysis. Nucleic Acids Res., 44, e117.

61. Wickramasinghe,V.O. and Laskey,R.A. (2015) Control of mammalian gene expression by selective mRNA export. Nat. Rev. Mol. Cell Biol., 16, 431–442.

62. Gehring,N.H., Wahle,E. and Fischer,U. (2017) Deciphering the mRNP Code: RNA-Bound Determinants of Post-Transcriptional Gene Regulation. Trends Biochem. Sci., 42, 369–382.

63. Hein,M.Y., Hubner,N.C., Poser,I., Cox,J., Nagaraj,N., Toyoda,Y., Gak,I.A., Weisswange,I., Mansfeld,J., Buchholz,F., et al. (2015) A human interactome in three quantitative dimensions organized by stoichiometries and abundances. Cell, 163, 712–723.

64. Huttlin,E.L., Ting,L., Bruckner,R.J., Gebreab,F., Gygi,M.P., Szpyt,J., Tam,S., Zarraga,G., Colby,G., Baltier,K., et al. (2015) The bioplex network: A systematic exploration of the human interactome. Cell, 162, 425–440.

65. Poser,I., Sarov,M., Hutchins,J.R.A., Hériché,J.-K., Toyoda,Y., Pozniakovsky,A., Weigl,D., Nitzsche,A., Hegemann,B., Bird,A.W., et al. (2008) BAC TransgeneOmics: a high-throughput method for exploration of protein function in mammals. Nat. Methods, 5, 409–415.

66. Cheeseman,I.M. and Desai,A. (2005) A combined approach for the localization and tandem affinity purification of protein complexes from metazoans. Sci. STKE, 2005, pl1.

67. Cristea,I.M. and Chait,B.T. (2011) Conjugation of magnetic beads for immunopurification of protein complexes. Cold Spring Harb. Protoc., 2011, pdb.prot5610.

68. Domanski,M., Molloy,K., Jiang,H., Chait,B.T., Rout,M.P., Jensen,T.H. and LaCava,J. (2012) Improved methodology for the affinity isolation of human protein complexes expressed at near endogenous levels. BioTechniques, 0, 1–6.

69. Jiang,H., Taylor,M.S., Molloy,K.R., Altukhov,I. and LaCava,J. (2019) Identification of RNase-sensitive LINE-1 Ribonucleoprotein Interactions by Differential Affinity Immobilization. Bio Protoc, 9.

70. Perez-Riverol,Y., Csordas,A., Bai,J., Bernal-Llinares,M., Hewapathirana,S., Kundu,D.J., Inuganti,A., Griss,J., Mayer,G., Eisenacher,M., et al. (2019) The PRIDE database and related tools and resources in 2019: improving support for quantification data. Nucleic Acids Res., 47, D442–D450.

71. Tyanova,S., Temu,T. and Cox,J. (2016) The MaxQuant computational platform for mass spectrometry-based shotgun proteomics. Nat. Protoc., 11, 2301–2319.

72. Bielow,C., Mastrobuoni,G. and Kempa,S. (2016) Proteomics quality control: quality control software for maxquant results. J. Proteome Res., 15, 777–787.

