## Supplemental Methods for "Affinity proteomic dissection of the human nuclear cap-binding complex interactome"

### Table of Contents


**Table of Contents** 1

**Supplemental Methods, Figures, and Text** 3
- Found in separate files: 3

**Supplemental Methods, Section 1** 3
- Cell line BAC information 3

**Supplemental Methods, Section 2** 4
- NCBP3-LAP screen QC 4

**Supplemental Methods, Section 3** 6
- MS method details 6
  - NCBP1-LAP - 24 different extraction solutions 6
    - Gel Bands 6
    - Gel Plugs 7
  - NCBP1-LAP and LAP-control - 4x6 different extraction solutions 7
    - Gel Plugs 7
  - NCBP2-LAP, NCBP3-LAP, and LAP-control - 4x6 different extraction solutions 8
    - Bands 8
    - Gel Plugs 8
  - NCBP3-LAP and LAP-control - 3x4 different extraction solutions (test tubes) 9
    - Gel Bands 9
    - Gel Plugs 10

**Supplemental Methods, Section 4** 10
- Merging MAGOH and MAGOHB 10

**Supplemental Methods, Section 5** 11
- Imputation of missing values 11
  - Perseus algorithm reproduction in R 12
  - New imputation algorithm 14
    - Method 3.1 15
    - Method 3.2 15
    - Method 3.3 15
    - Method 4.1 16
    - Method 4.2 16
    - Method 4.3 16
    - Method 4.4 16
    - Method 4.5 17

Method 4.6 17
Method 4.7 17
Performance estimation methods 18
Correlation between replicates 19
ROC curves 19
Root Mean Square Error 20
Selection of MNAR imputation method 21
Correlation heatmap 21

**Supplemental Methods, Section 6** **22**
GFP Normalization 22
Statistical filtering 23

**References** **23**

#### Supplemental Methods, Figures, and Text

Found in separate files:

1. Supplemental_Data_Tables.xlxs
    a. Book 1: Legend and replicate information for MS analysis
    b. Book 2: 24 conditions NCBP1-LAP pre-screen
    c. Book 3: 4x6 conditions NCBP1-LAP and LAP-control, multi-well
    d. Book 4: 4x6 conditions NCBP2-LAP and LAP-control, multi-well
    e. Book 5: 4x6 conditions NCBP3-LAP and LAP-control, multi-well (omitted from this study)
    f. Book 6: 3x4 conditions NCBP3-LAP and control, microfuge tubes
2. Supplemental_Data_pvalues_logfc.xlsx
3. Supplemental_heatmap.pdf
4. 3D_Animated_MDS.gif
5. Supplemental_Data_replicate_correlation_plots.pdf
6. RMSE_NRMSE_plots.zip
7. GFP_normalization.pdf
8. FN_proteins_never_passed_ttest.csv
9. FN_proteins_passed_ttest_once.csv

#### Supplemental Methods, Section 1

##### Cell line BAC information

| Cell line | Protein name | BAC ID | Tagging cassette |
|---|---|---|---|
| CTRL-LAP | (LAP-tag) | CTD-3000G10 | N-terminal |
| NCBP1-LAP | NCBP1 | RP11-17A21 | C-terminal |
| NCBP2-LAP | NCBP2 | HS.E139.A23 | C-terminal |
| NCBP3-LAP | NCBP3 | RP11-118B14 | C-terminal |

#### Supplemental Methods, Section 2

##### NCBP3-LAP screen QC

Several factors indicated that the NCBP3-LAP multi-well screen did not yield sufficient quality data for integrative analysis, motivating us to exclude that data and repeat sample production at a larger scale - the following are example indicators. We observed the location of the target protein in the distribution of all protein intensities. The intensity of target protein in the NCBP3-LAP multi-well data was lower and exhibited more variation than the other dataset (**Fig. S1**).

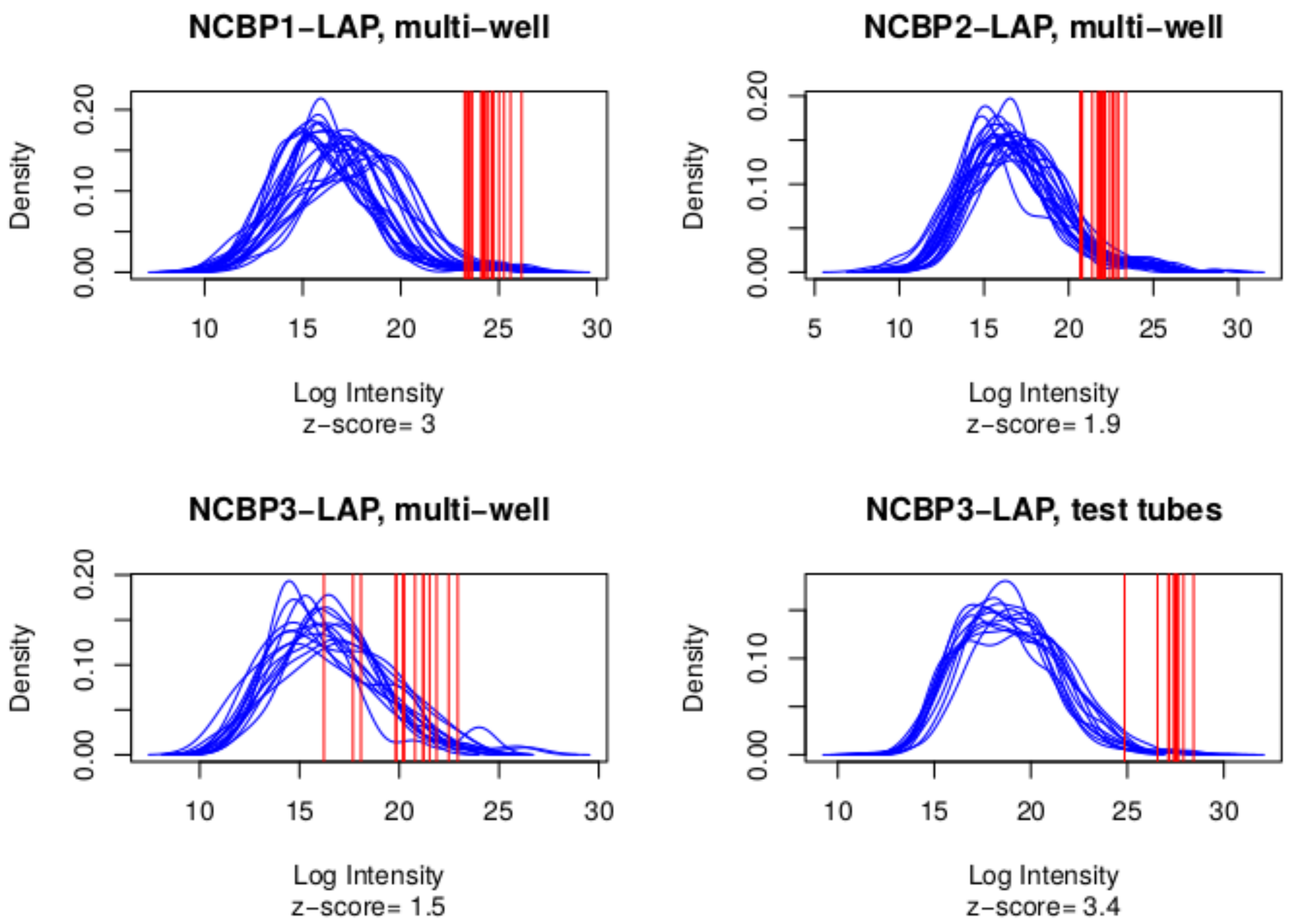


**Figure S1** - Distribution of protein iBAQ intensities in different experiments. Red lines show the location of target protein intensities. Multiple lines in each plot account for multiple conditions and replicates. Z-score: mean((target intensity - mean intensity) / sd(intensity)).

We observed that the contaminant abundance was in general higher in the NCBP3-LAP multi-well experiments compared to the others (**Fig. S2**). Other experiments exhibited a greater degree of dispersion across the different samples. We also observed that both before and after imputation replicates in these experiments had less correlation with each other than the other experiments (**Section 5, Fig. S11**).

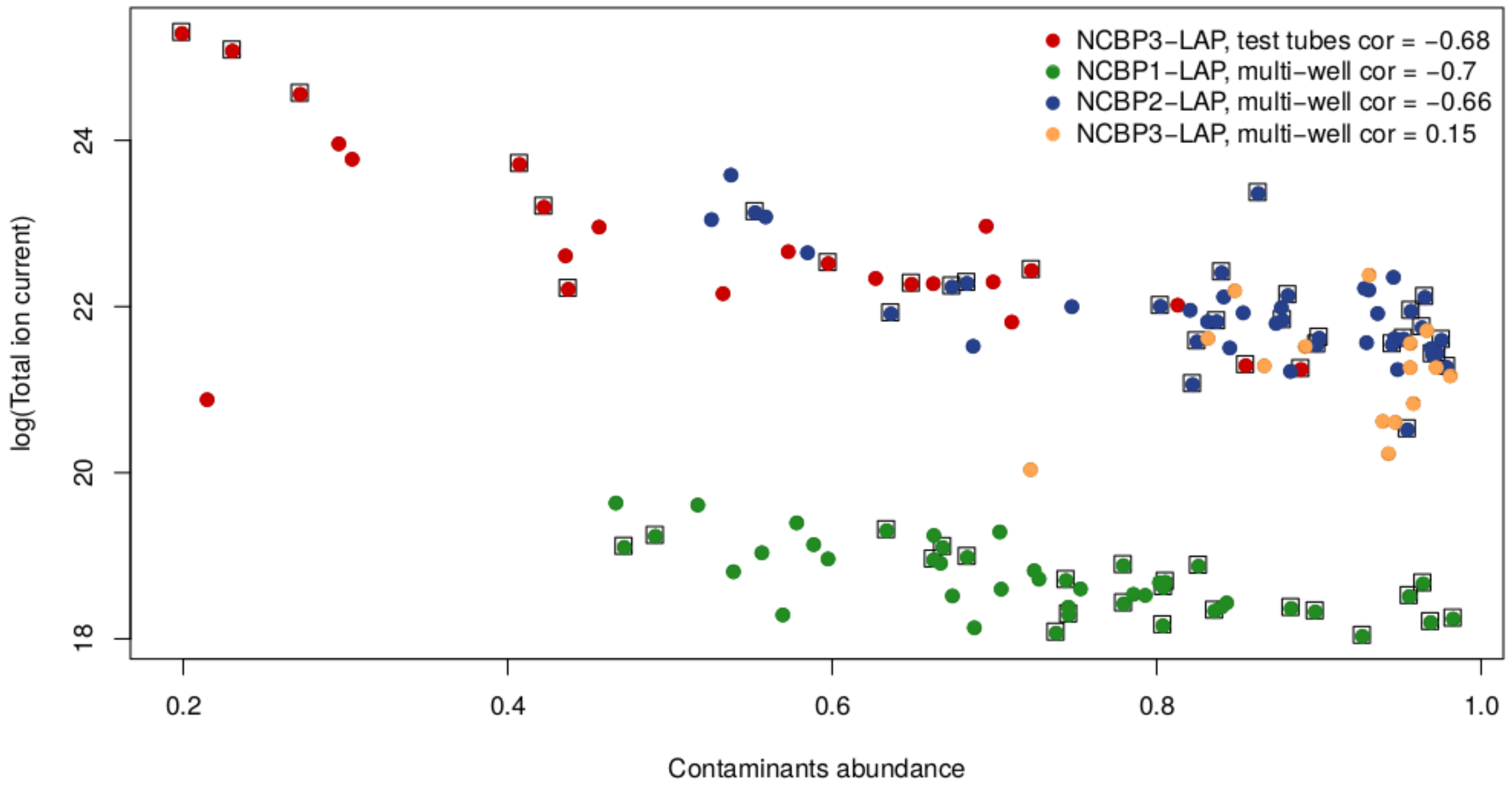


**Figure S2** - Dependence between Total ion current (ln(x), Y-axis) and contaminants abundance (X-axis) across the experiments. Contaminants abundance = \frac{\sum{contaminant LFQ intensity}}{\sum{all LFQ intensity}}. The control experiments are marked by black squares; NCBP2-LAP and NCBP3-LAP done in multi-well used the same control (blue dot with black square) . NCBP1 samples were run on an Orbitrap Fusion mass spectrometer whereas others were run on a Q Exactive Plus.

To assess the consequences of omitting the NCBP3-LAP multi-well data, we compared two distance matrices (before and after removing the data) with Mantel's test using 1000 permutations, which showed that the two matrices had correlation = 0.96 with p-value = 0.001 (**Fig. S3**). Thus, we concluded that these data did not significantly contribute to our understanding of NCBP protein-proteins relationships, and could be omitted.

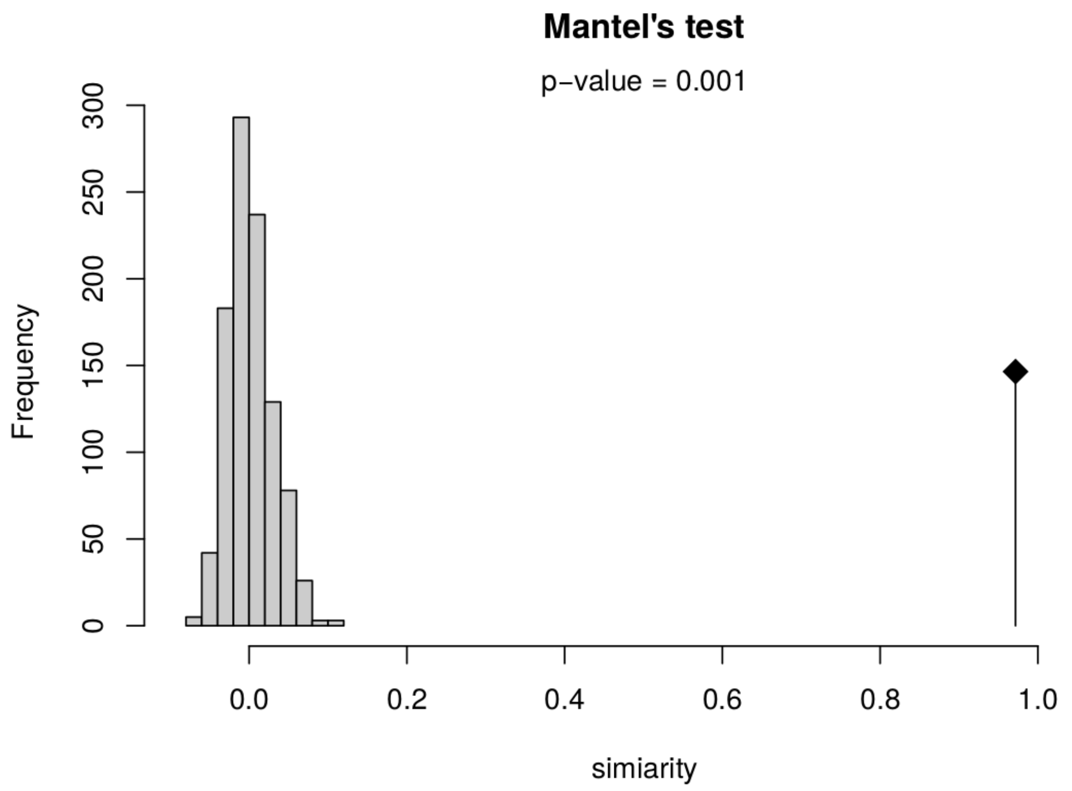


**Figure S3** - Comparison of the distance matrix with and without the NCBP3-LAP multi-well data. On the left is the density estimate of the permutation distribution. The vertical line on the right is the observed Z-statistic. The Z-statistic for the Mantel test is equal to the sum of the pairwise product of the lower triangles of the permuted matrices, for each permutation of rows and columns.

#### Supplemental Methods, Section 3

##### MS method details

###### NCBP1-LAP - 24 different extraction solutions

*Gel Bands*

Instrument: Orbitrap Fusion

Solvent B: 0.1% formic acid in acetonitrile

Gradient: 5-30%  solvent B in 7 min

1 second cycle

*MS1*

Resolution: 60k

range: 350-1700 m/z

100 ms max injection time

5e5 ion accumulation target

*MS2*

Detection: ion trap

4 s dynamic exclusion

Fragmentation: CID with 35% collision energy

200 ms max injection time

2.2e4 ion accumulation target

##### *Gel Plugs*

Instrument: Q Exactive Plus

Solvent B: 0.1% formic acid in acetonitrile

Gradient: 4-30%  solvent B in 40 min

*MS1*

Resolution: 70k

range: 350-1500 m/z

500 ms max injection time

3e6 ion accumulation target

*MS2*

Resolution: 17.5k

15 s dynamic exclusion

Fragmentation: HCD with 28% normalized collision energy

200 ms max injection time

1e5 ion accumulation target

Top 10 most abundant precursors selected for MS2

#### NCBP1-LAP and LAP-control - 4x6 different extraction solutions

##### *Gel Plugs*

Instrument: Orbitrap Fusion

Solvent B: 0.1% formic acid in acetonitrile

Gradient: 2-30%  solvent B in 35 min

4 second cycle

*MS1*

Resolution: 120k

range: 300-1500 m/z

500 ms max injection time

2e5 ion accumulation target

*MS2*

Detection: ion trap

15 s dynamic exclusion

Fragmentation: CID with 35% collision energy

200 ms max injection time

2.2e4 ion accumulation target

NCBP2-LAP, NCBP3-LAP, and LAP-control - 4x6 different extraction solutions

*Bands*

Instrument: Q Exactive Plus

Solvent B: 0.1% formic acid in 95% acetonitrile

Gradient: 2-30%  solvent B in 7 min

*MS1*

Resolution: 70k

range: 300-1700 m/z

100 ms max injection time

1e6 ion accumulation target

*MS2*

Resolution: 17.5k

4 s dynamic exclusion

Fragmentation: HCD with 24% normalized collision energy

200 ms max injection time

2e5 ion accumulation target

Top 5 most abundant precursors selected for MS2

*Gel Plugs*

Instrument: Q Exactive Plus

Solvent B: 0.1% formic acid in 95% acetonitrile
Gradient: 2-32%  solvent B in 35 min
*MS1*
Resolution: 70k
range: 350-1500 m/z
500 ms max injection time
3e6 ion accumulation target
*MS2*
Resolution: 17.5k
15 s dynamic exclusion
Fragmentation: HCD with 24% normalized collision energy
100 ms max injection time
2e5 ion accumulation target
Top 20 most abundant precursors selected for MS2

NCBP3-LAP and LAP-control - 3x4 different extraction solutions (test tubes)

*Gel Bands*

Instrument: Q Exactive HF
Solvent B: 0.1% (v/v) formic acid in acetonitrile
Gradient: 5-30% solvent B in 20 min
*MS1*
Resolution: 60k
range: 375-2000 m/z
100 ms max injection time
3e6 ion accumulation target
*MS2*
Resolution: 15k
10 s dynamic exclusion
Fragmentation: HCD with 27% normalized collision energy
100 ms max injection time
1e5 ion accumulation target
Top 10 most abundant precursors selected for MS2

*Gel Plugs*

Instrument: Q Exactive HF

Solvent B: 0.1% (v/v) formic acid in acetonitrile

Gradient: 2-25% solvent B in 53 min

*MS1*

Resolution: 60k

range: 375-2000 m/z

100 ms max injection time

3e6 ion accumulation target

*MS2*

Resolution: 15k

15 s dynamic exclusion

Fragmentation: HCD with 27% normalized collision energy

50 ms max injection time

2e5 ion accumulation target

Top 15 most abundant precursors selected for MS2

#### Supplemental Methods, Section 4

##### Merging MAGOH and MAGOHB

```
MAGOHB            1 MAVASDFYLRYYVGHKGKFGHEFLEFEFRPDGKLRYANNSNYKNDVMIRK     50
                      :.||||||||||||||||||||||||||||||||||||||||||||||
MAGOH             1 --MESDFYLRYYVGHKGKFGHEFLEFEFRPDGKLRYANNSNYKNDVMIRK     48

MAGOHB           51 EAYVHKSVMEELKRIIDDSEITKEDDALWPPPDRVGRQELEIVIGDEHIS    100
                    ||||||||||||||||||||||||||||||||||||||||||||||||||
MAGOH            49 EAYVHKSVMEELKRIIDDSEITKEDDALWPPPDRVGRQELEIVIGDEHIS     98

MAGOHB          101 FTTSKIGSLIDVNQSKDPEGLRVFYYLVQDLKCLVFSLIGLHFKIKPI    148
                    ||||||||||||||||||||||||||||||||||||||||||||||||
MAGOH            99 FTTSKIGSLIDVNQSKDPEGLRVFYYLVQDLKCLVFSLIGLHFKIKPI    146
```

**Figure S4.** Sequence alignment of MAGOH and MAGOHB: four amino acids difference at the N-terminus.

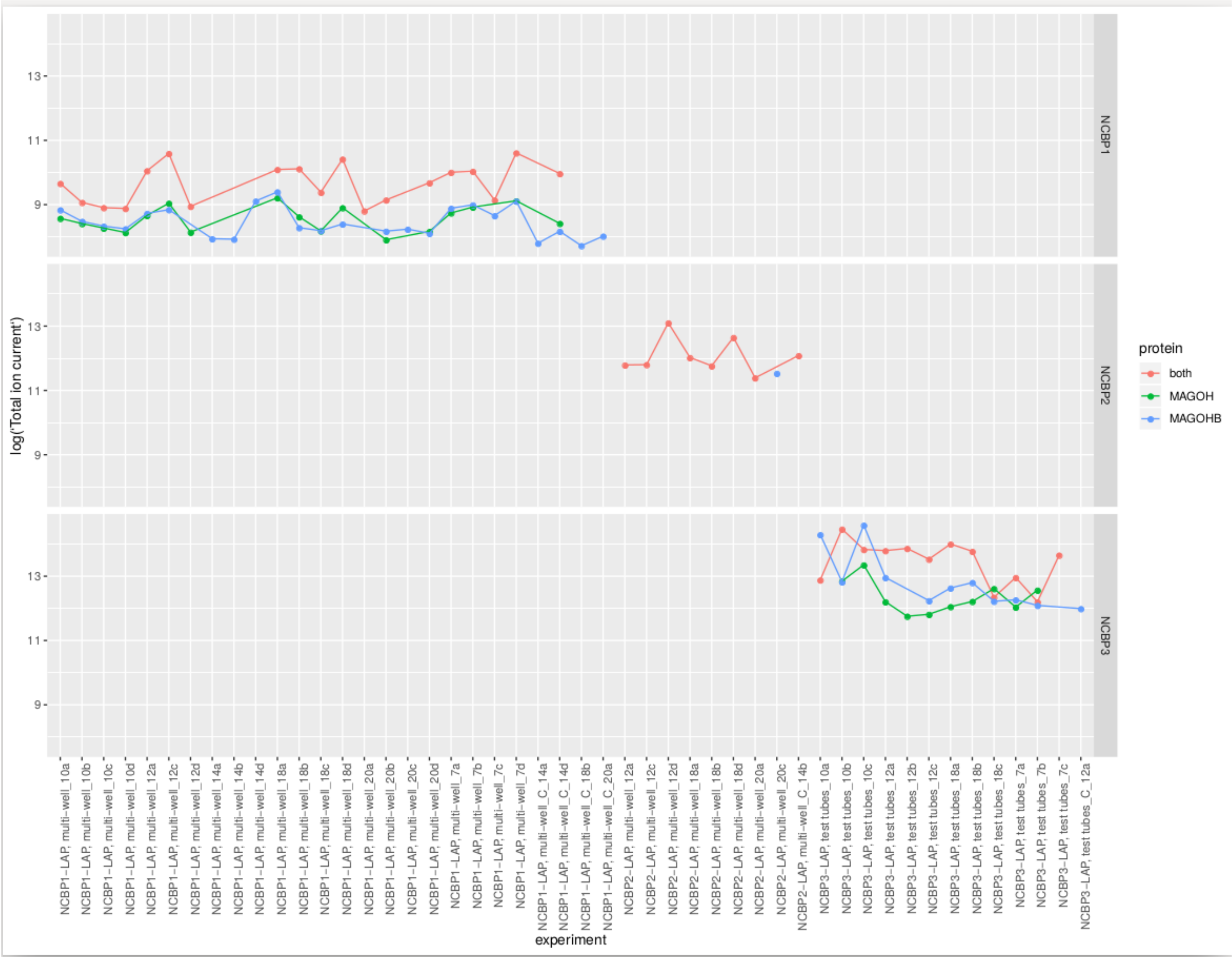


**Figure S5.** Total ion current (TIC, ln(x); Y-axis)) of MAGOH and MAGOHB peptides in different experiments (X-axis). The red color shows the mean TIC of peptides which can be denoted both to MAGOH and MAGOHB. The green and the blue colors show TICs for the unique peptides. In NCBP1 experiments the intensities of MAGOH and MAGOHB unique peptides are nearly the same. In NCBP2 experiments these proteins were not well distinguished. In some NCBP3 experiments the MAGOHB unique peptide was more prominent.

#### Supplemental Methods, Section 5

##### Imputation of missing values

The data included missing values. Zero values can be explained either as the real absence of the protein in the sample or as a failure to observe polypeptides due to the sampling characteristics of data dependent, shotgun mass spectrometry analyses. These missing values interfere with the calculation of $\log_2$FC and statistical comparative analysis; imputation is a way of dealing with incomplete data. Imputation is not without challenges, for example, to avoid introducing artifacts into the data; this is an area of active ongoing research in mass spectrometry data processing (1–3). We developed a custom imputation algorithm to increase performance with our data and benchmarked our method against the imputation algorithm used by the Perseus software (4).

##### Perseus algorithm reproduction in R

To compare new methods with the Perseus algorithm, it was reproduced in R and consisted of the following steps: (1) calculate the mean ($\mu$) and standard deviation (sd) of $\log_2$ intensities of all non-zero proteins for every replicate; (2) replace every missing value by number, sampled from normal distribution with the following parameters: $sd_{new} = sd * 0.3$ ; $\mu_{new} = \mu - 1.8 * sd$. To be sure that the Perseus algorithm implemented in R authentically reproduces the original Perseus algorithm, the imputed values were compared for NCBP3-LAP, microfuge tube data. Most of the values are intensities which were originally non-zero and are the same for both methods. The imputed values (**Fig. S6**) are very similar between the original and reproduced Perseus method, the deviations are limited to $\pm 2$ and are connected with random sampling.

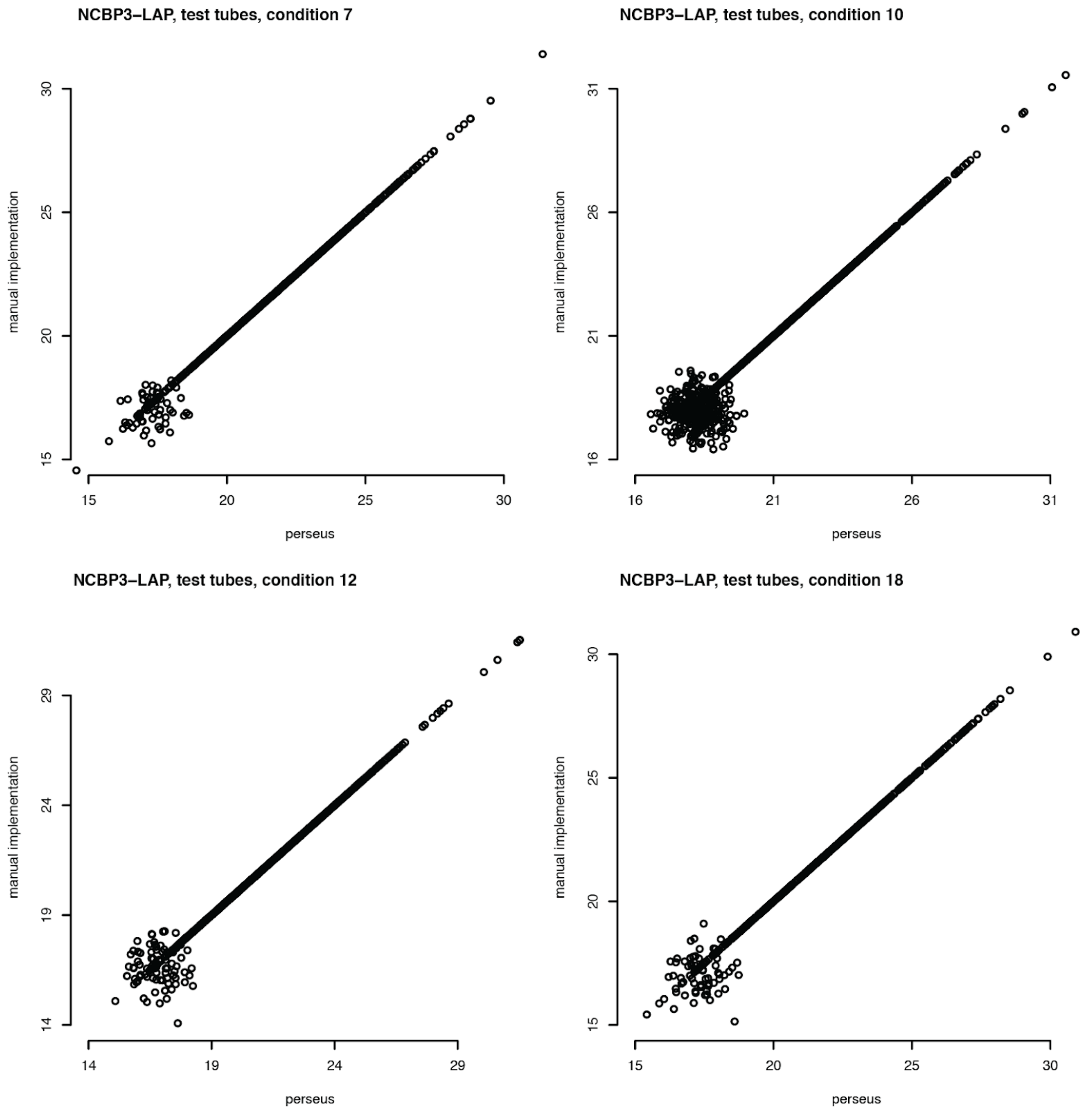


**Figure S6.** Comparison of Perseus and our implementation of the Perseus imputation algorithm for four example NCBP-LAP samples, as labeled. On the x-axis are values output from the Perseus software. On the y-axis are values output from the Perseus algorithm reproduced in R.

New imputation algorithm

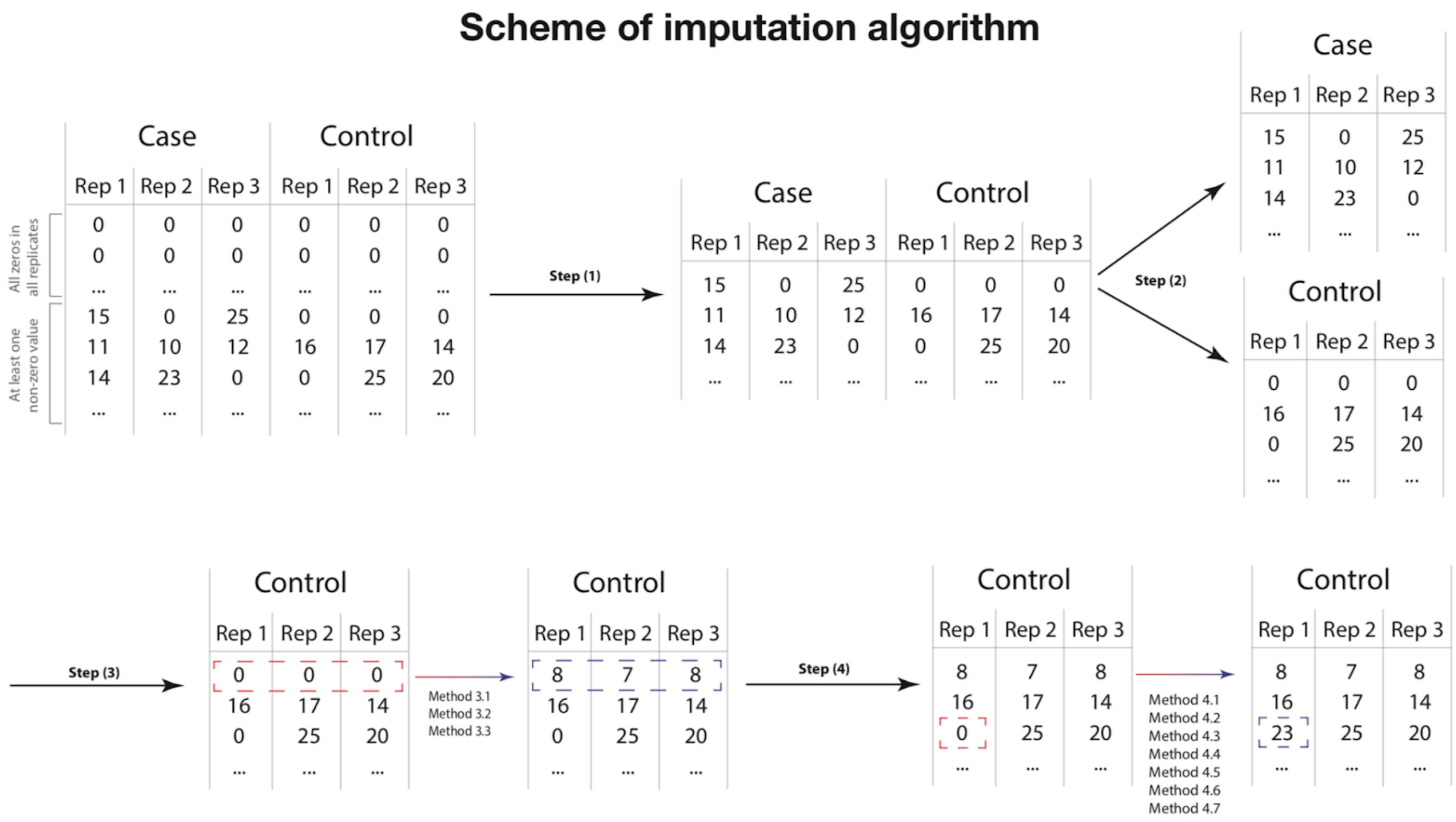


**Figure S7.** In general the new imputation algorithm has the following steps:

1. separate experiments by groups (each group includes the experimental [case] and control IPs produced under identical conditions) and process each group separately;
2. remove proteins not identified both in cases and controls, process case and control replicates separately;
3. impute small values for proteins that have zeros in all case or control replicates;
4. impute values for proteins that have zero intensities in some case or control replicates;

The main advance included within our algorithm was the use of data from three or four replicates for each case and control IP. The 3$^{rd}$ step in the list above is used to impute values which are thought to be real zeros but needed to have some small value for the $\log_2$FC calculation during ANOVA: multiple failures to observe data across numerous replicates indicates a likely true zero (at the level of detection applied). The 4$^{th}$ step in the list above was used to restore missing values when some non-zero values exist among the replicates and includes information from non-zero replicates: occasional detection (or failure thereof) may be due to MS sampling and fluctuations around the detection baseline (especially for low abundance proteins), rather than true absence.

We have developed and tested three distinct methods for the 3rd step and seven different methods for the 4th step.

Assuming an analysis that yields data including zero values in **all** *case* **or** *control* replicates for some proteins, 3rd step imputation methods address these zero values and include the following:

*Method 3.1*

For each replicate, the mean and standard deviation of non-zero protein intensities are counted. New intensities for each missing value in each replicate are sampled from a uniform distribution:

**(I)** $$Int_{new} = Unif\{start = mean(Int_{replica}) - 3 * sd(Int_{replica}), end = mean(Int_{replica}) - 2 * sd(Int_{replica})\}$$

where $Int_{new}$ is a new imputed intensity, $Int_{replica}$ is a vector of all protein intensities in the replicate and $Unif\{start, end\}$ is a uniform distribution. The parameters 2 and 3 were chosen as standard parameters from the Perseus algorithm.

*Method 3.2*

The replicate with the least number of zero values is chosen and the mean and standard deviation of the non-zero proteins intensities are calculated. New intensities for the zero-value proteins in this replicate are sampled from uniform distribution (see (I), above). The values for the other replicates are imputed further in the 4th step.

*Method 3.3*

For each replicate mean and standard deviation of non-zero protein intensities are calculated. New intensities for each missing value in each replicate are sampled from a normal distribution:

**(II)** $$Int_{new} = \mathcal{N}\{\mu = mean(Int_{replica}) - 1.8 * sd(Int_{replica}), sd = 0.3 * sd(Int_{replica})\}$$

where $\mathcal{N}\{\mu, sd\}$ is a normal distribution. This method reproduces how the Perseus algorithm treats missing values.

4th step methods impute values to replace zeros when proteins receive some zero **and** non-zero values in *case* **and/or** *control* replicates. These include the following:

a. Build distribution of deltas for all non zero proteins, where delta:

**(III)** $$\Delta = \frac{Int_{rep1} - Int_{rep2}}{mean(Int_{rep1}, Int_{rep2})}$$

b. Calculate $\mu_\Delta, sd_\Delta$

c. Calculate new delta and new intensity:

*Method 4.1*

**(IV)** $$\Delta_{new} = \mathcal{N}\{\mu = \mu_\Delta, sd = \frac{\sqrt{2} * sd_\Delta}{mean(corr)}\},$$

where $mean(corrs)$ is a mean correlation between vector of all proteins intensities of imputed replicates and replicates which don't have zeros in current imputed protein intensity, $Int_{other}$ are intensities of the protein in non-zero replicates. The assumption which motivates this parameter is that higher value imputed replicates should correlate with non-zero replicates. If correlation is in general very low, the delta of intensity for the imputed value will be sampled from normal distribution with large standard deviation and vice versa.

After that the intensities to be imputed are calculated using this formula:

**(V)** $$Int_{new} = mean(Int_{other} * \mid 1 + \Delta_{new} \mid).$$

*Method 4.2*

**(VI)** $$\Delta_{new} = \mathcal{N}\{\mu = \mu_\Delta, sd = \frac{\sqrt{2} * sd_\Delta}{mean(corr)} * NumberOfZeros\},$$

where $NumberOfZeros$ - number of replicates in which current imputed protein has missing values. The hypothesis is that the higher number of zeros there are for the protein, the lower should be imputed intensity. New intensity is calculated using **(V).**

*Method 4.3*

Delta new is calculated using **(IV)**

**(VII)** $$Int_{new} = mean(Int_{other} * \mid 1 + \Delta_{new} \mid *(1 - \log_{10} NumberOfZeros))$$

*Method 4.4*

Delta new is calculated using **(VI)**

**(VIII)** $$Int_{new} = mean(Int_{other} * \mid 1 + \Delta_{new} \mid * \frac{1 - NumberOfZeros}{TotalNumber})$$

*Method 4.5*

**(IX)** $$\Delta_{new} = \mathcal{N}\{\mu = \mu_{\Delta}, sd = \frac{\sqrt{2} * sd_{\Delta}}{mean(corr)} * (1 + \log_{10} NumberOfZeros)\}$$

New intensity is calculated using **(V)**

*Method 4.6*

$$\Delta_{new} = \mathcal{N}\{\mu = \mu_{\Delta}, sd = \frac{sd_{\Delta}}{\sqrt{2} * mean(corr)}\}$$

New intensity is calculated using **(V)**

*Method 4.7*

For outliers (proteins, which have zero values in all but one replicate) this method accommodates one of the three methods for imputation of all zero replicates. For other proteins method 4.6 is used.

#### Performance estimation methods

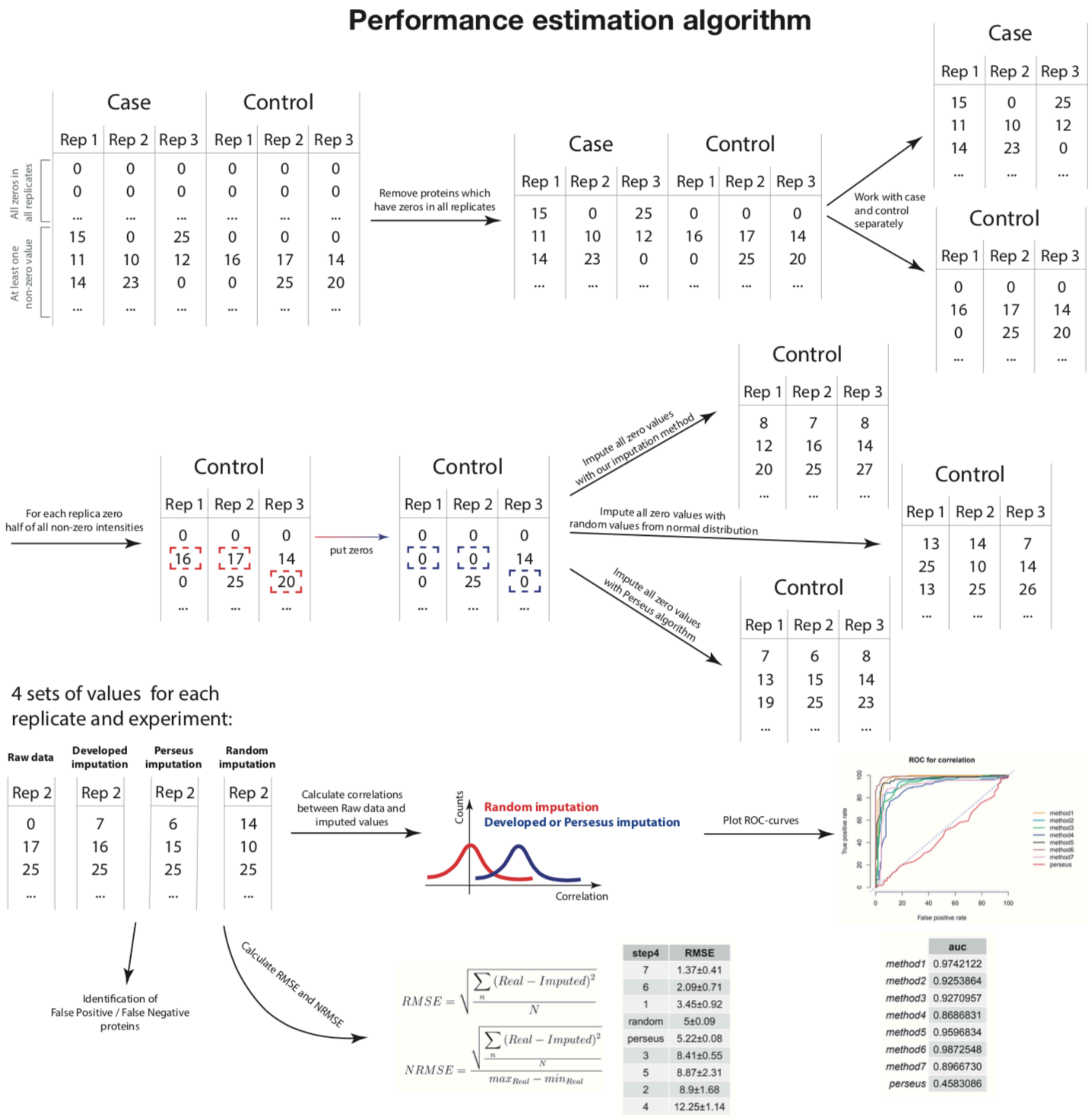


**Figure S8.** To select the best imputation method and to compare our methods with existing alternatives we estimated the algorithms performances using the data produced in this study. Three methods of efficiency control were used which measured distinct qualities of the imputation: correlation between replicates, ROC-AUC (area under the curve), and RMSE (root mean square error).

##### Correlation between replicates

Each imputation method was applied to the whole dataset. For every group of cases and controls, the imputed values for proteins from a replicate were plotted against the non-zero initial values for the same proteins in other replicates. The plots are provided in **Supplemental_Data_replicate_correlation_plots.pdf**. Every page in the document represents one group of experiments (cases or controls). On each page the first three columns represent method 3.1, method 3.2, method 3.3 respectively for the imputation of intensities for proteins which have zero values in all replicates; the 4th column is a correlation matrix of all replicates after imputation using method 3.1. Rows represent methods from step 4. These plots were used to select between methods 3.1 - 3.3.

##### ROC curves

ROC-curves were built according to the following algorithm:

1. Take all the proteins' intensities that are >0 in at least one replicate
2. For each replicate convert half of all non-zero intensities to zeros
3. Impute all zero values with our imputation methods
4. Impute all zero values with random values from normal distribution (mean and sd taken from the distribution of all intensities in these replicates).
5. Calculate two correlations for each replicate
    a. The correlation of methodically imputed data with original data
    b. The correlation of randomly imputed data with original data
    c. These correlations were counted using only those values that were deleted during step 2 of this algorithm.
6. Make distributions for correlations with methodically and randomly imputed data
7. Plot ROC-curves from these distributions (**Fig. S9**)

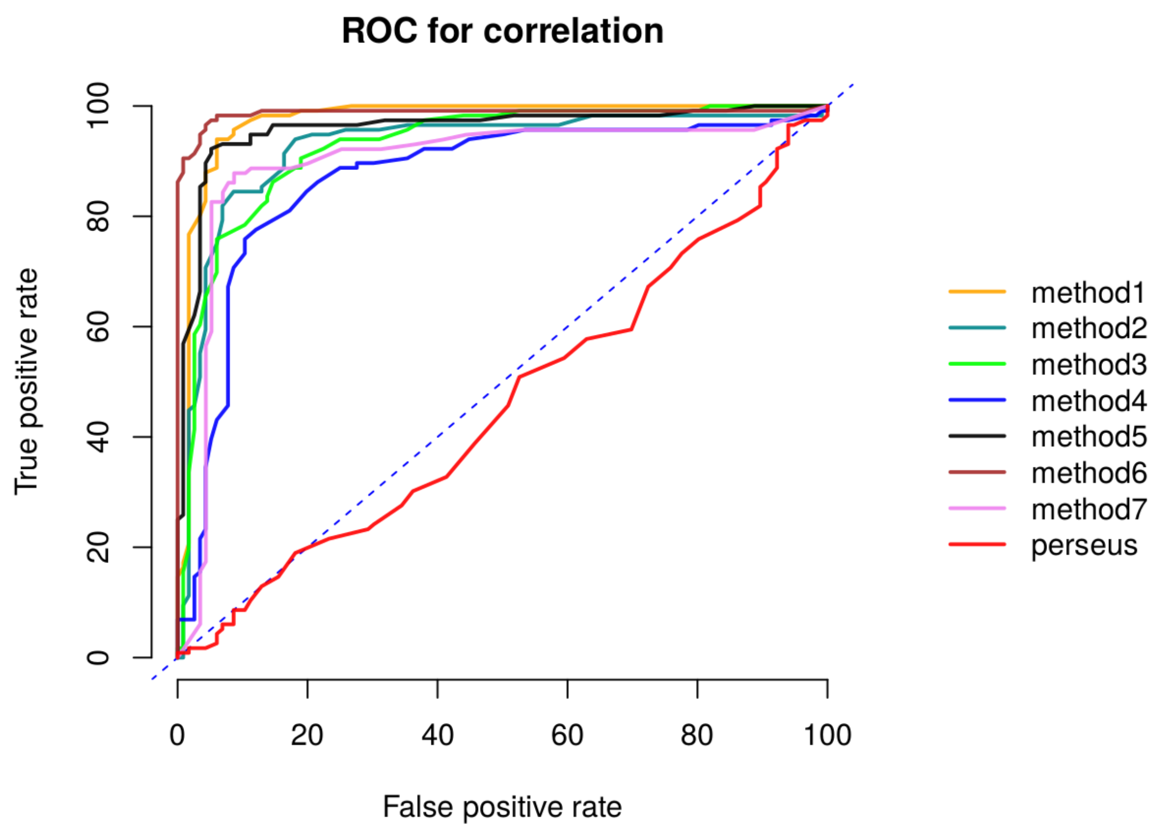


| method | AUC |
|---|---|
| method6 | 0.99±0.05 |
| method1 | 0.97±0.09 |
| method5 | 0.95±0.09 |
| method7 | 0.93±0.08 |
| method3 | 0.92±0.11 |
| method2 | 0.91±0.1 |
| method4 | 0.87±0.11 |
| perseus | 0.5±0.13 |

**Figure S9.** LFQ intensity ROC curves for different imputation methods with AUC; 97.5% confidence interval from 10 trials.

#### Root Mean Square Error

Another metric we used for performance estimation was RMSE.

$$RMSE = \sqrt{\frac{\sum_n (Real - Imputed)^2}{N}}$$ - root-mean-square error

$$NRMSE = \frac{\sqrt{\frac{\sum_n (Real - Imputed)^2}{N}}}{max_{Real} - min_{Real}}$$ - normalized root-mean-square error

For every method, RMSE was assessed as follows: some values from real data were set to zero, as described in the ROC-curve section (see Steps 1-2). RMSE was calculated as square root from the mean of squared errors of imputed intensities for every protein in every experiment. The procedure was repeated 15 times to estimate 97.5% confidence intervals for every method's RMSE. Method 7 produced the lowest RMSE at 1.37 ±0.41, followed by method 6, 1, random imputation, Perseus, and others. For every method NRMSE was calculated: the values of NRMSE were computed for every experiment. After that they were plotted, with x-axis representing the experiment, y-axis showing the NRMSE value (see files, indicated below). Both RMSE and NRMSE were calculated only for values imputed by method 4.1- 4.6 and Perseus, but not by method 3.1- 3.3. RMSE was not used to evaluate imputation of proteins which had zeros in all replicates; it was calculated for the “random” method, when the values were sampled from a uniform distribution. The plots showing RMSE/NRMSE results and additional plots with results by ROC/AUC are located in the file **RMSE_NRMSE_plots.zip**

#### Selection of MNAR imputation method

For a selection of the best MNAR (missing not at random) imputation algorithms (step 3) we have calculated the number of false positives (FP) and false negatives (FN) proteins. FPs are proteins that passed the t-test in a case/control comparison but had one or zero non-zero values among case replicates before imputation. FNs are proteins that did not pass the case/control t-test despite having two or more non-zero intensity values among cases and only one or zero non-zero values among controls. Counts of FP and FN are presented in the figure below **(Fig. S10**); the best result (low FPs and low FNs) was obtained using method 3.1 for MNAR imputation.

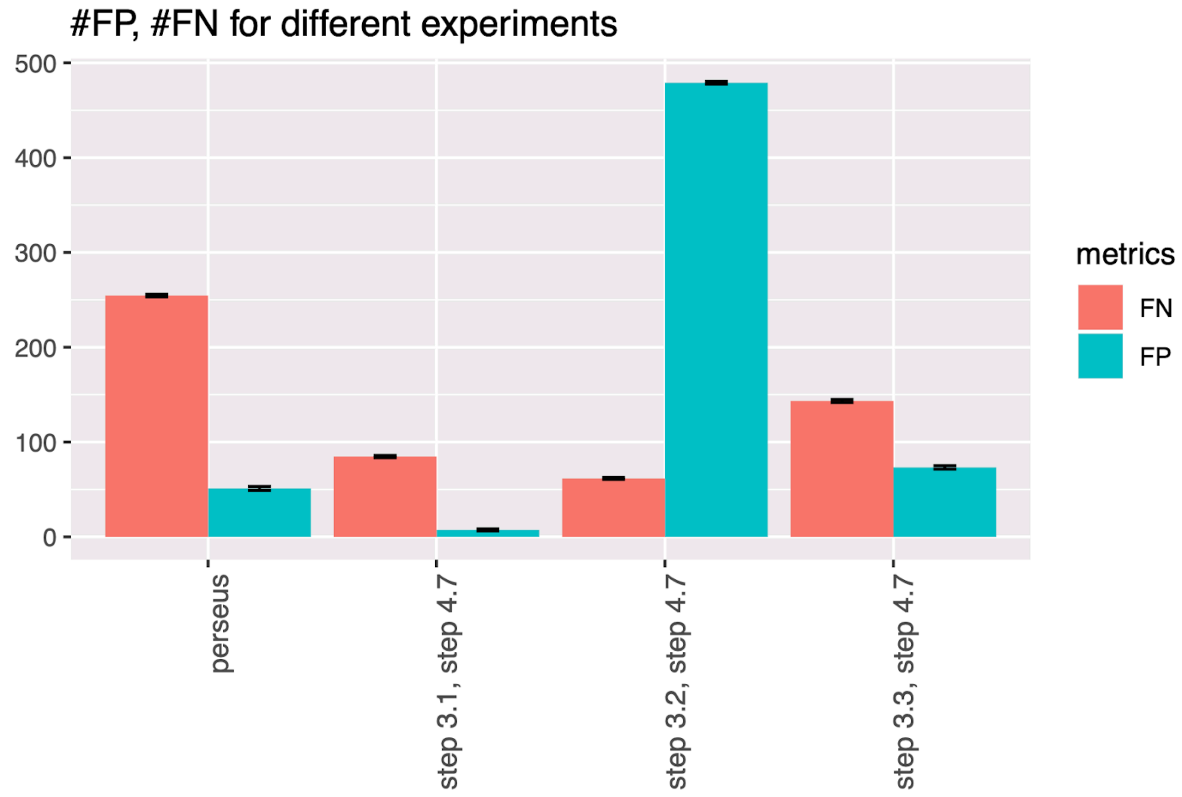


**Figure S10.** Bar plot of FPs and FNs generated by different imputation approaches. We calculated 97.5% confidence intervals for 20 runs of the procedure.

#### Correlation heatmap

Applying our imputation algorithm improved the correlations between replicates and across groups, shown below (**Fig. S11**).

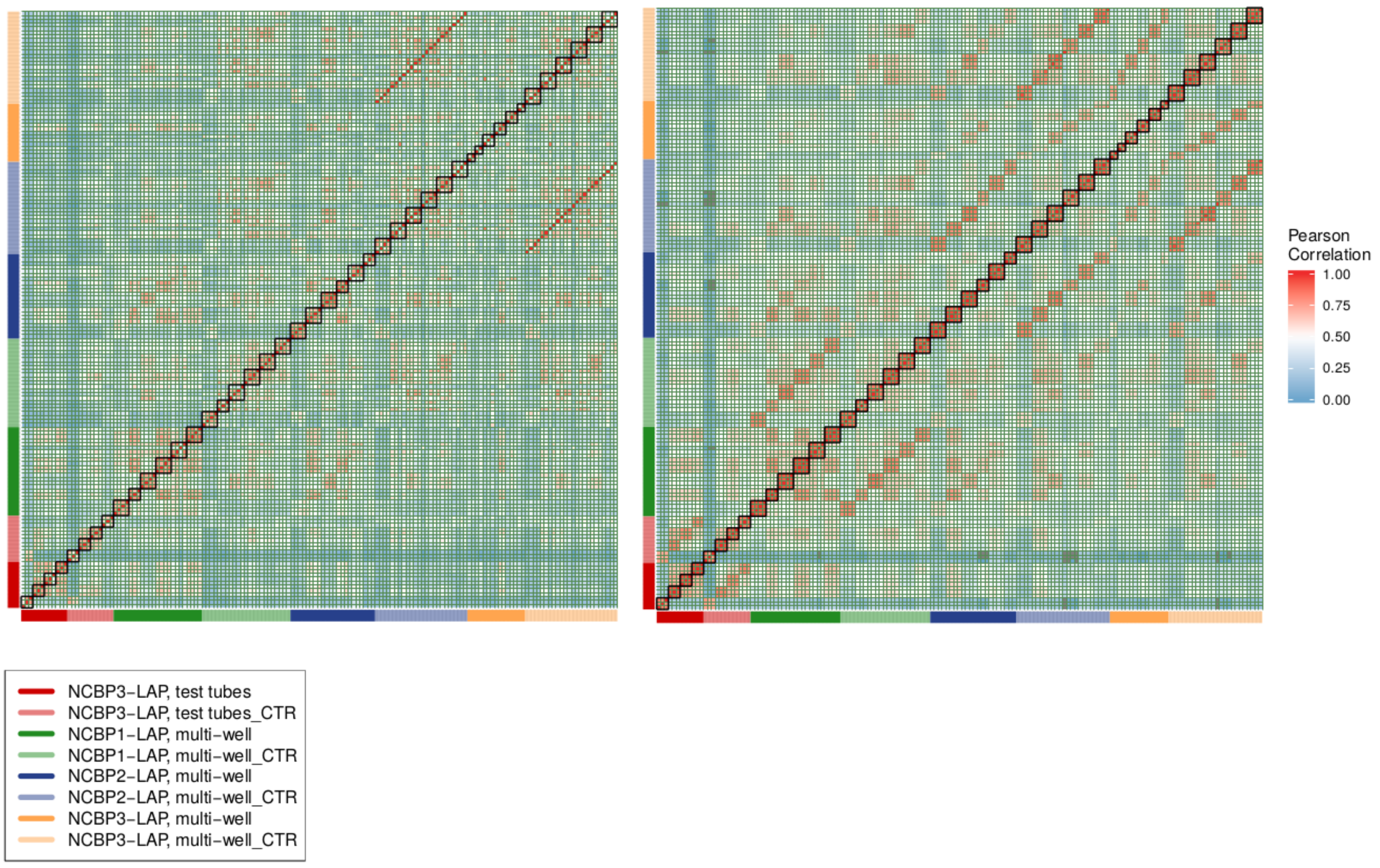


**Figure S11.** Correlation heatmap of LFQ intensities before and after imputation. On the x and y axes, the experiments are shown, the color distinguishes IP, bright colors mean cases and matched faded colors mean controls (see legend). Black borders on the diagonal denote replicate groups for different experimental conditions.

#### Supplemental Methods, Section 6

##### GFP Normalization

See file: **GFP_normalization.pdf**, which shows the distribution of selected protein LFQ intensities across experiments before normalization and after. Each color represents one protein. The grey color denotes the minimal and maximal intensity in every experiment. GFP was chosen for this transformation to enable fair cross samples comparisons because the LAP-tag was the target of the IPs using anti-GFP antibodies. Unlike NCBP proteins, which fluctuate across conditions and controls, GFP was explicitly present in every experiment as a consequence of the design, and it was (as expected) close to maximum intensity in all the experiments. Apparent fluctuations in the yield of NCBPs and their interactors due to fluctuations in GFP yield could be thus compensated for.

After normalization, we observed that the intensity fluctuations of the target proteins were smoothed.

##### Statistical filtering

P-values and $log_2$FC values for 100 runs of the imputation algorithm (described in **Section 5**) were obtained and used to produce a table with mean protein intensity p-values and $log_2$FC, at 97.5% confidence intervals, which were used in further analysis for proteins that passed statistical significance in ≥ 60 of the 100 runs (voting). After voting, no imputation induced FPs remained; FNs were removed from the analysis and provided as supplemental files for manual inspection in two separate classes: FNs that passed a t-test in a different experimental conditions, and FNs that never passed a t-test (FN_proteins_never_passed_ttest.csv, FN_proteins_passed_ttest_once.csv; see Selection of MNAR imputation method, **Figure S10**, for the description of FP and FN**)**. Hence, we view this procedure as stringent FP control.

#### References

1. Webb-Robertson,B.-J.M., Wiberg,H.K., Matzke,M.M., Brown,J.N., Wang,J., McDermott,J.E., Smith,R.D., Rodland,K.D., Metz,T.O., Pounds,J.G., *et al.* (2015) Review, evaluation, and discussion of the challenges of missing value imputation for mass spectrometry-based label-free global proteomics. *J. Proteome Res.*, **14**, 1993–2001.

2. Välikangas,T., Suomi,T. and Elo,L.L. (2018) A comprehensive evaluation of popular proteomics software workflows for label-free proteome quantification and imputation. *Brief. Bioinformatics*, **19**, 1344–1355.

3. Lazar,C., Gatto,L., Ferro,M., Bruley,C. and Burger,T. (2016) Accounting for the Multiple Natures of Missing Values in Label-Free Quantitative Proteomics Data Sets to Compare Imputation Strategies. *J. Proteome Res.*, **15**, 1116–1125.

4. Tyanova,S., Temu,T., Sinitcyn,P., Carlson,A., Hein,M.Y., Geiger,T., Mann,M. and Cox,J. (2016) The Perseus computational platform for comprehensive analysis of (prote)omics data. *Nat. Methods*, **13**, 731–740.
