## Supplemental Figures and Tables for "Affinity proteomic dissection of the human nuclear cap-binding complex interactome": Supplemental_Data_replicate_correlation_plots.pdf

NCBP1−LAP, multi−well, condition 7

cases

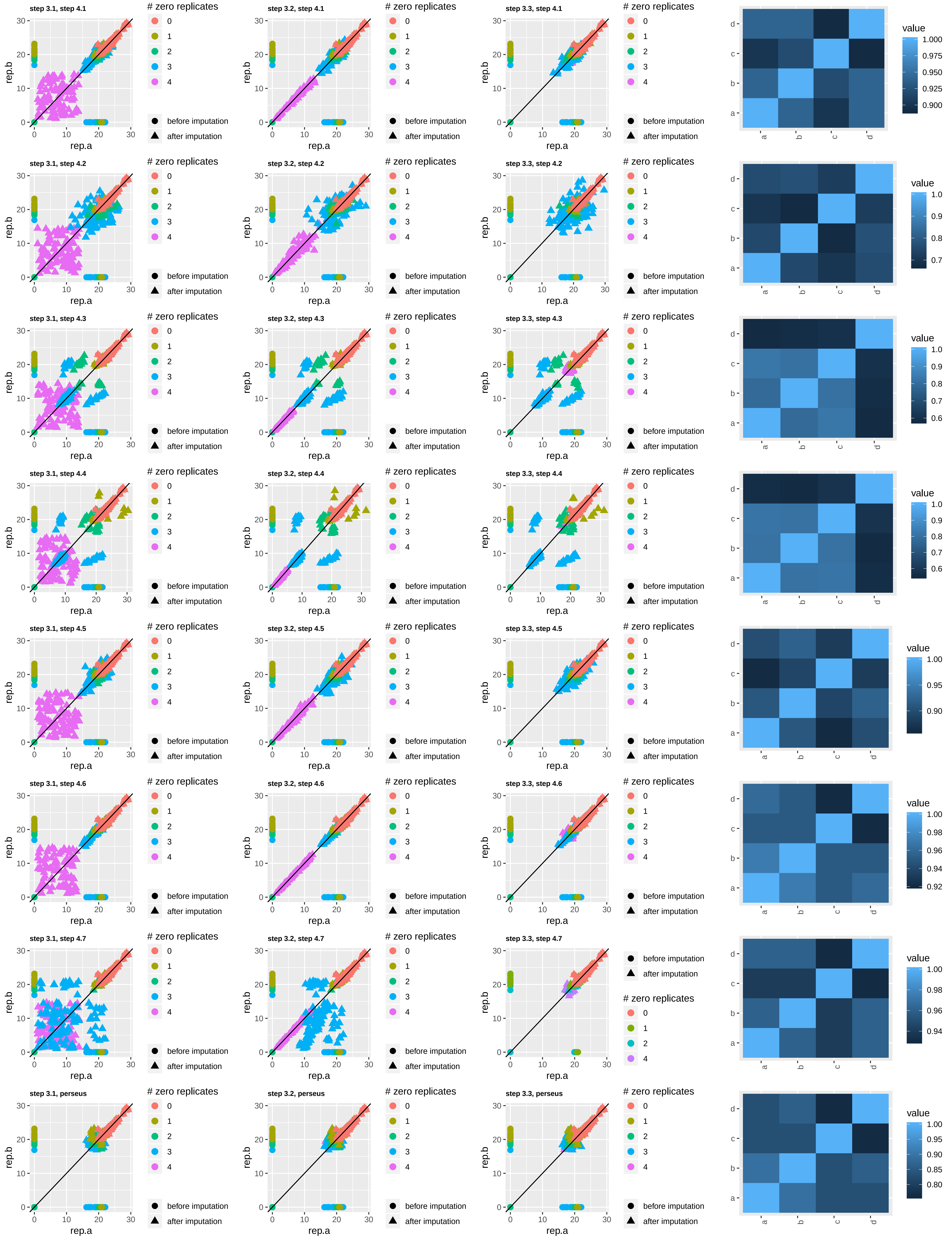

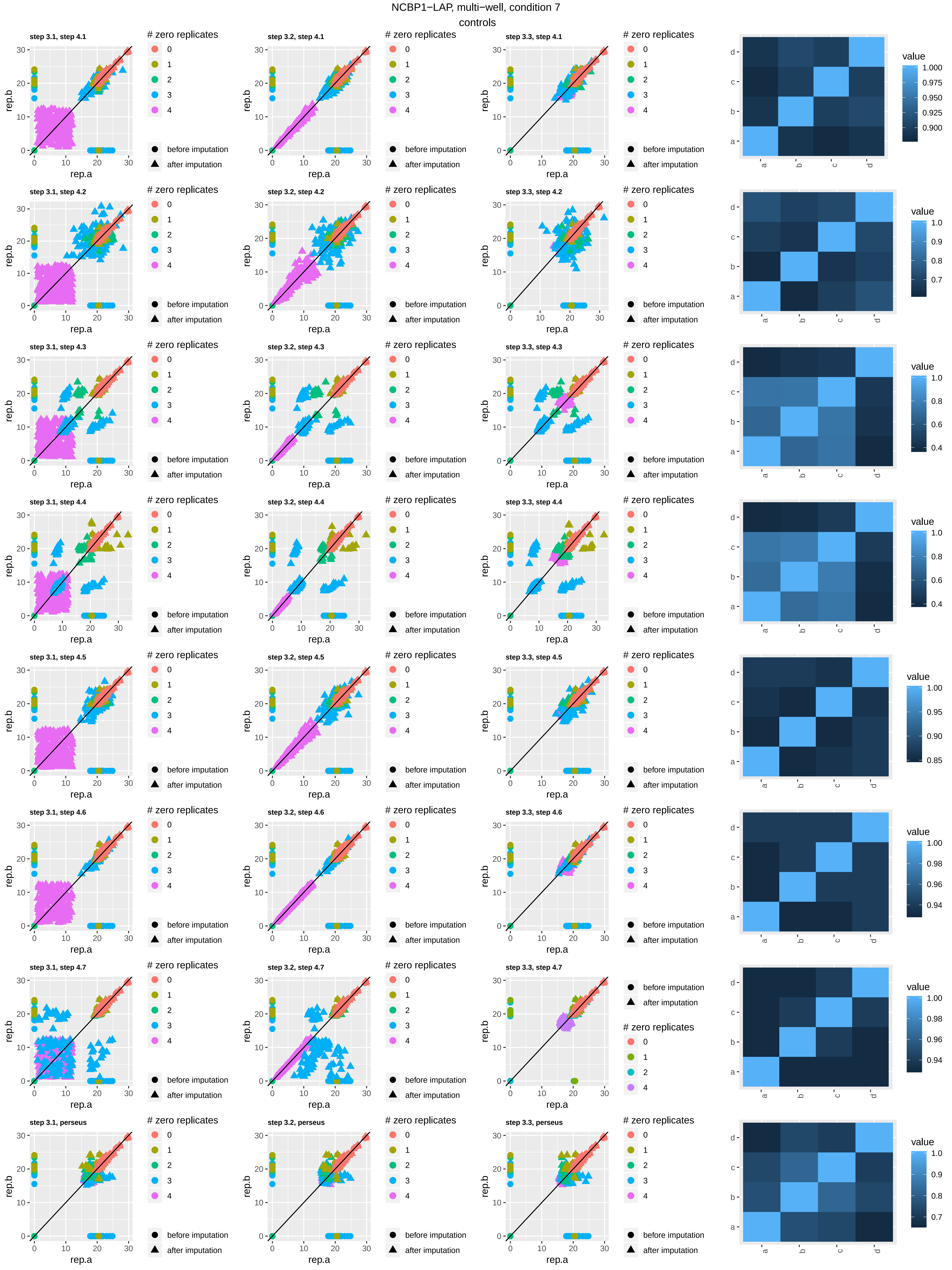

NCBP1−LAP, multi−well, condition 7
controls
step 3.1, step 4.1
step 3.2, step 4.1
step 3.3, step 4.1
step 3.1, step 4.2
step 3.2, step 4.2
step 3.3, step 4.2
step 3.1, step 4.3
step 3.2, step 4.3
step 3.3, step 4.3
step 3.1, step 4.4
step 3.2, step 4.4
step 3.3, step 4.4
step 3.1, step 4.5
step 3.2, step 4.5
step 3.3, step 4.5
step 3.1, step 4.6
step 3.2, step 4.6
step 3.3, step 4.6
step 3.1, step 4.7
step 3.2, step 4.7
step 3.3, step 4.7
step 3.1, perseus
step 3.2, perseus
step 3.3, perseus
rep.a
rep.b
### zero replicates
0
1
2
3
4
before imputation
after imputation
value
a
b
c
d

NCBP1−LAP, multi−well, condition 10

cases

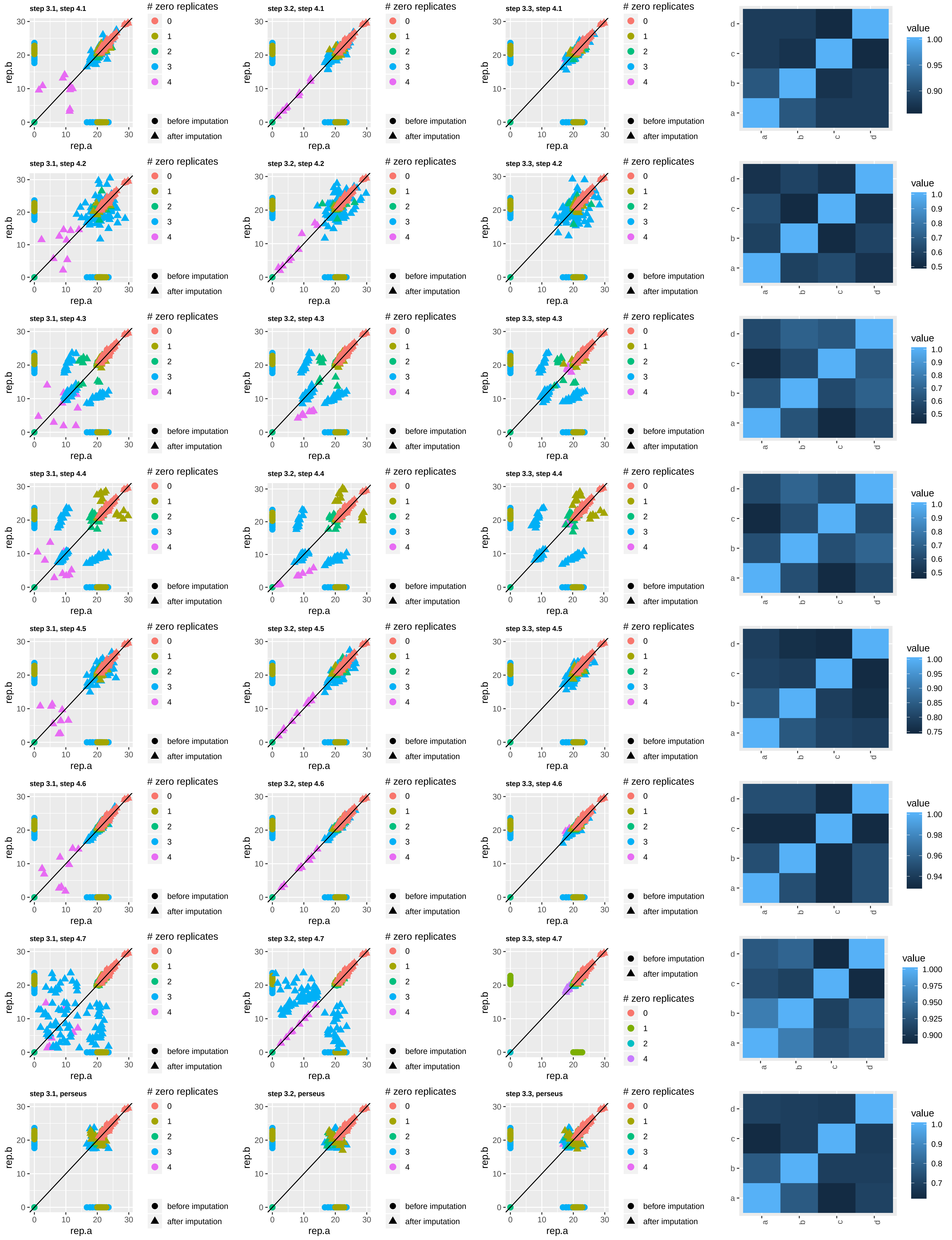

NCBP1−LAP, multi-well, condition 10
controls

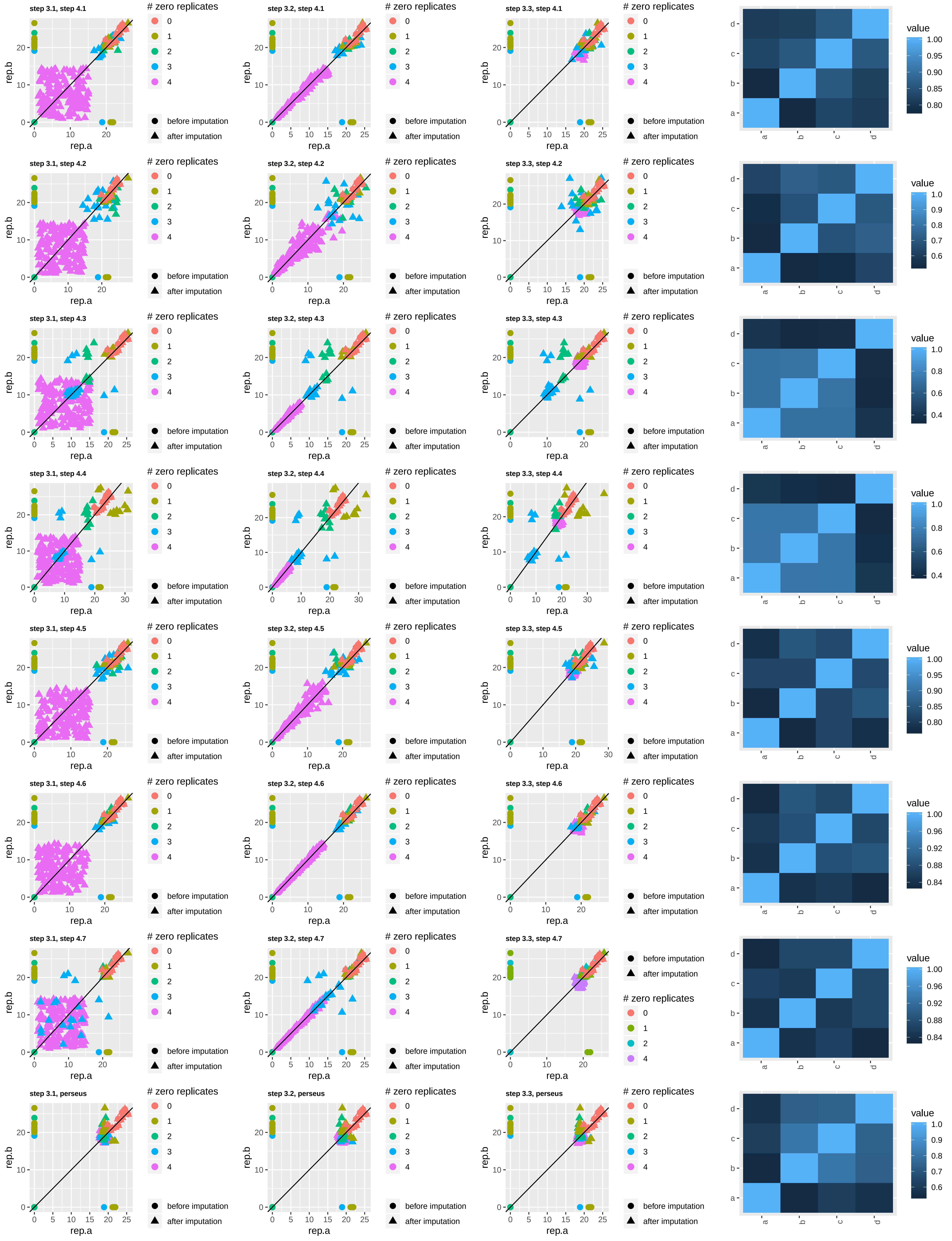

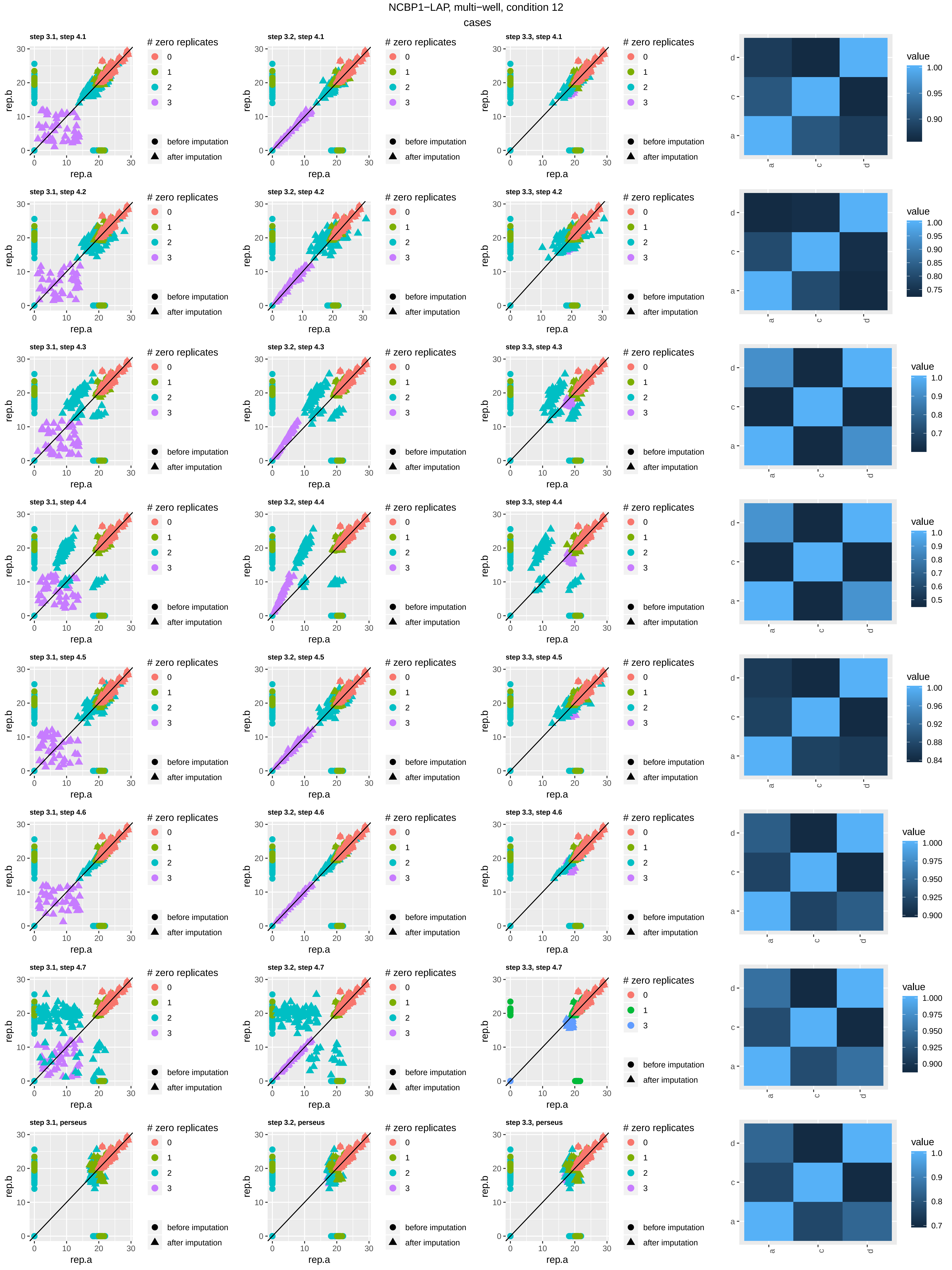

NCBP1−LAP, multi−well, condition 12
cases
step 3.1, step 4.1
step 3.2, step 4.1
step 3.3, step 4.1
step 3.1, step 4.2
step 3.2, step 4.2
step 3.3, step 4.2
step 3.1, step 4.3
step 3.2, step 4.3
step 3.3, step 4.3
step 3.1, step 4.4
step 3.2, step 4.4
step 3.3, step 4.4
step 3.1, step 4.5
step 3.2, step 4.5
step 3.3, step 4.5
step 3.1, step 4.6
step 3.2, step 4.6
step 3.3, step 4.6
step 3.1, step 4.7
step 3.2, step 4.7
step 3.3, step 4.7
step 3.1, perseus
step 3.2, perseus
step 3.3, perseus
rep.a
rep.b
### zero replicates
0
1
2
3
before imputation
after imputation
value
a
c
d

NCBP1−LAP, multi−well, condition 12

controls

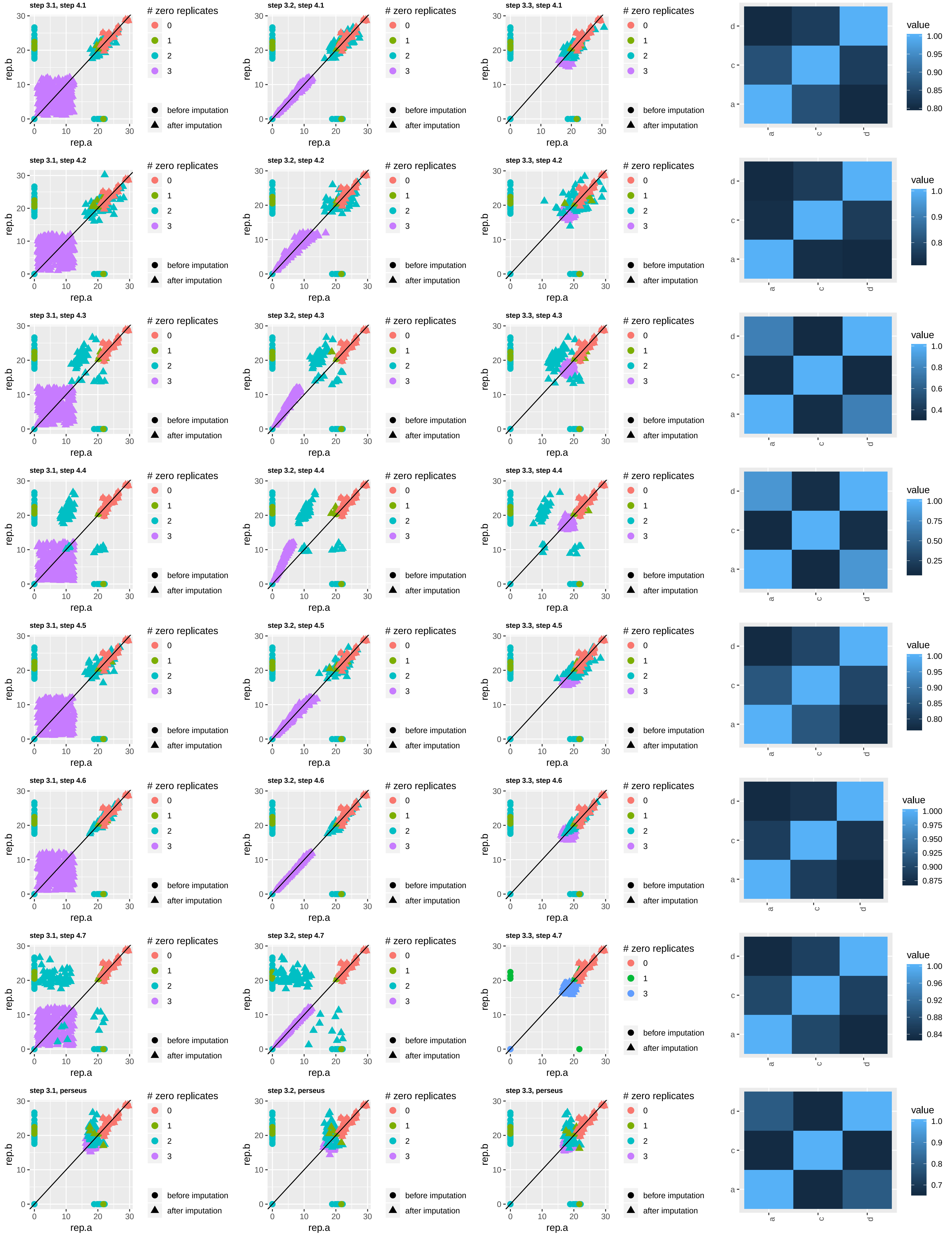

NCBP1−LAP, multi−well, condition 14 cases

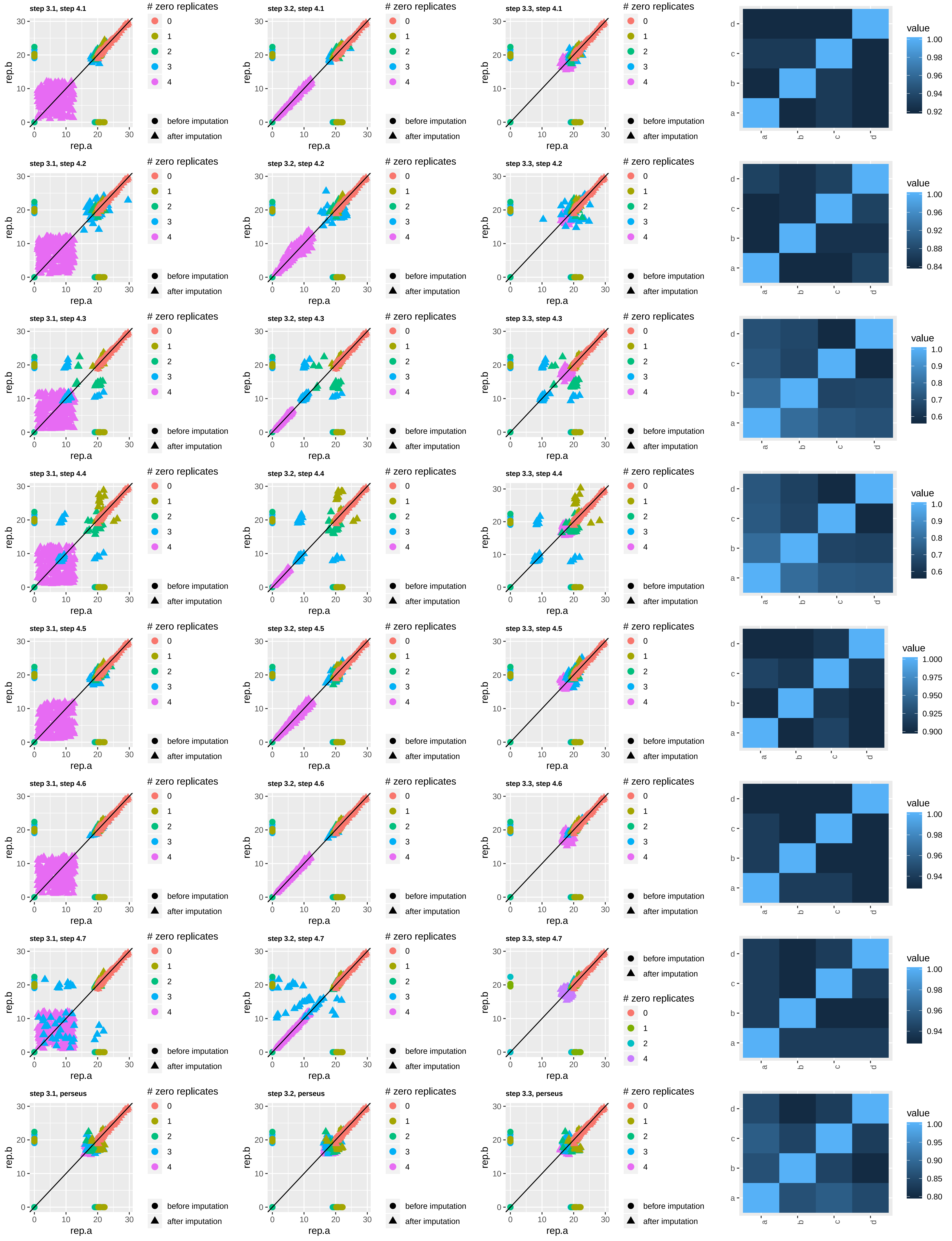

NCBP1−LAP, multi-well, condition 14

controls

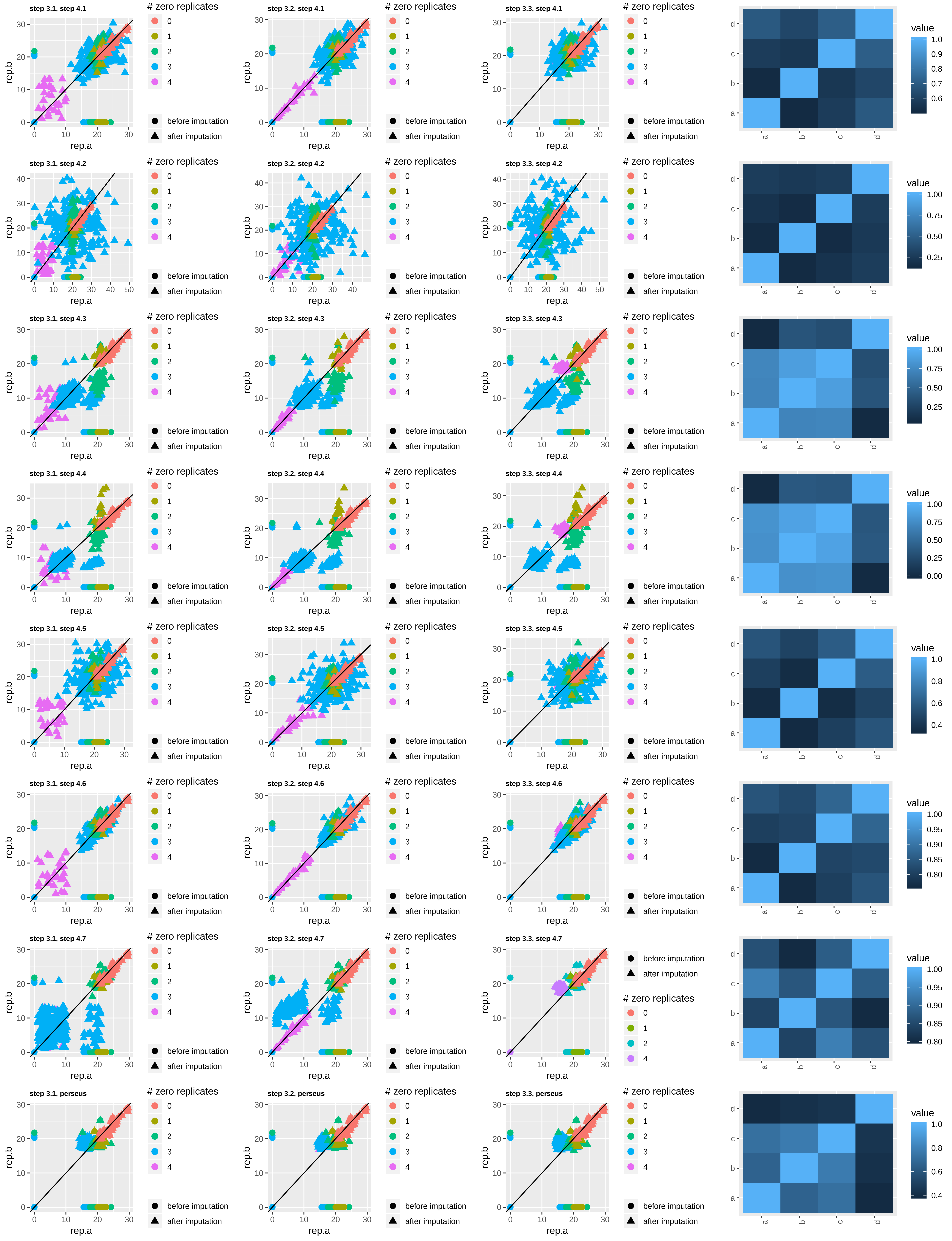

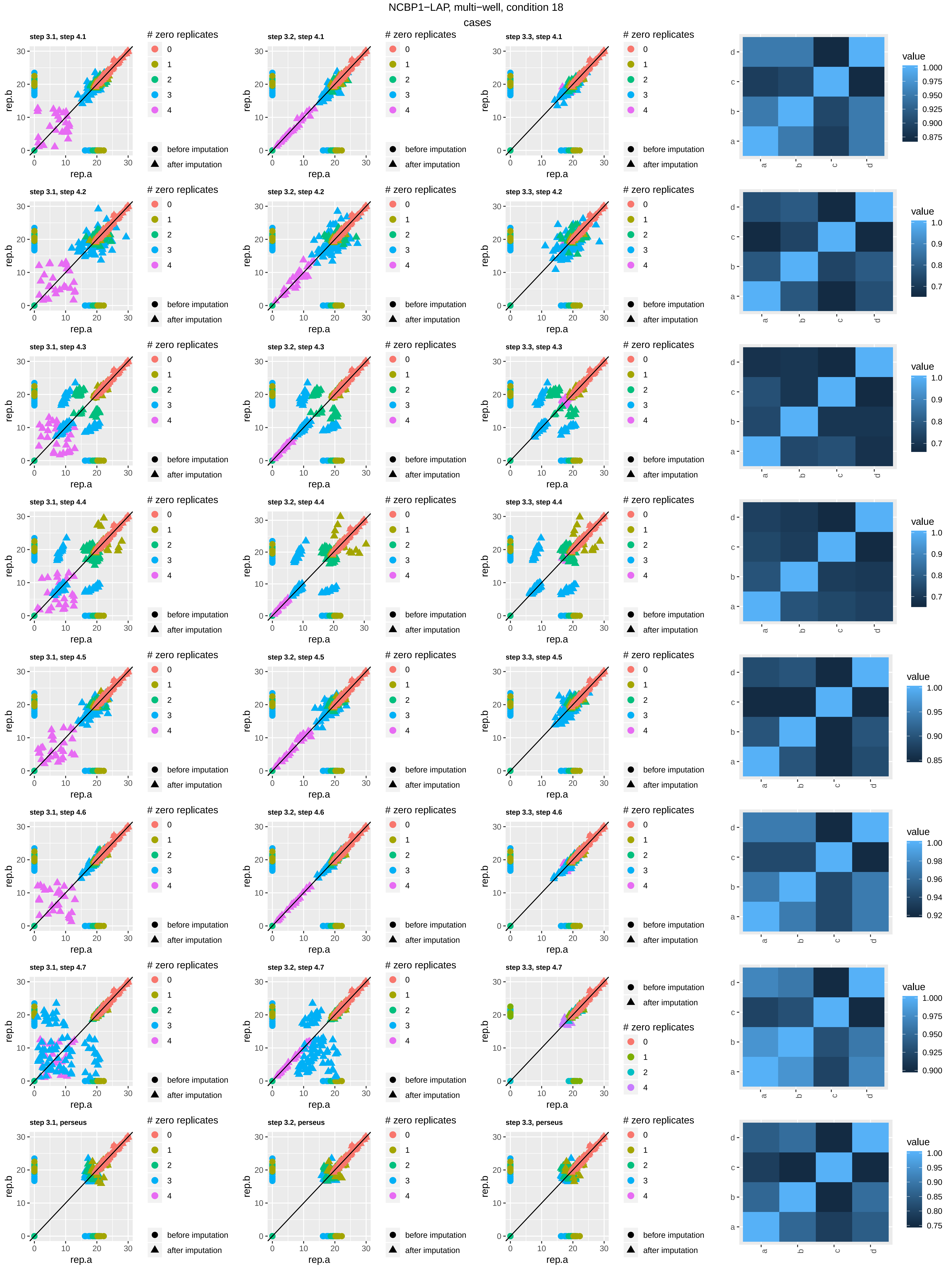
NCBP1−LAP, multi-well, condition 18
cases
step 3.1, step 4.1
step 3.2, step 4.1
step 3.3, step 4.1
step 3.1, step 4.2
step 3.2, step 4.2
step 3.3, step 4.2
step 3.1, step 4.3
step 3.2, step 4.3
step 3.3, step 4.3
step 3.1, step 4.4
step 3.2, step 4.4
step 3.3, step 4.4
step 3.1, step 4.5
step 3.2, step 4.5
step 3.3, step 4.5
step 3.1, step 4.6
step 3.2, step 4.6
step 3.3, step 4.6
step 3.1, step 4.7
step 3.2, step 4.7
step 3.3, step 4.7
step 3.1, perseus
step 3.2, perseus
step 3.3, perseus
rep.a
rep.b
### zero replicates
0
1
2
3
4
before imputation
after imputation
value
a
b
c
d

NCBP1−LAP, multi-well, condition 18

controls

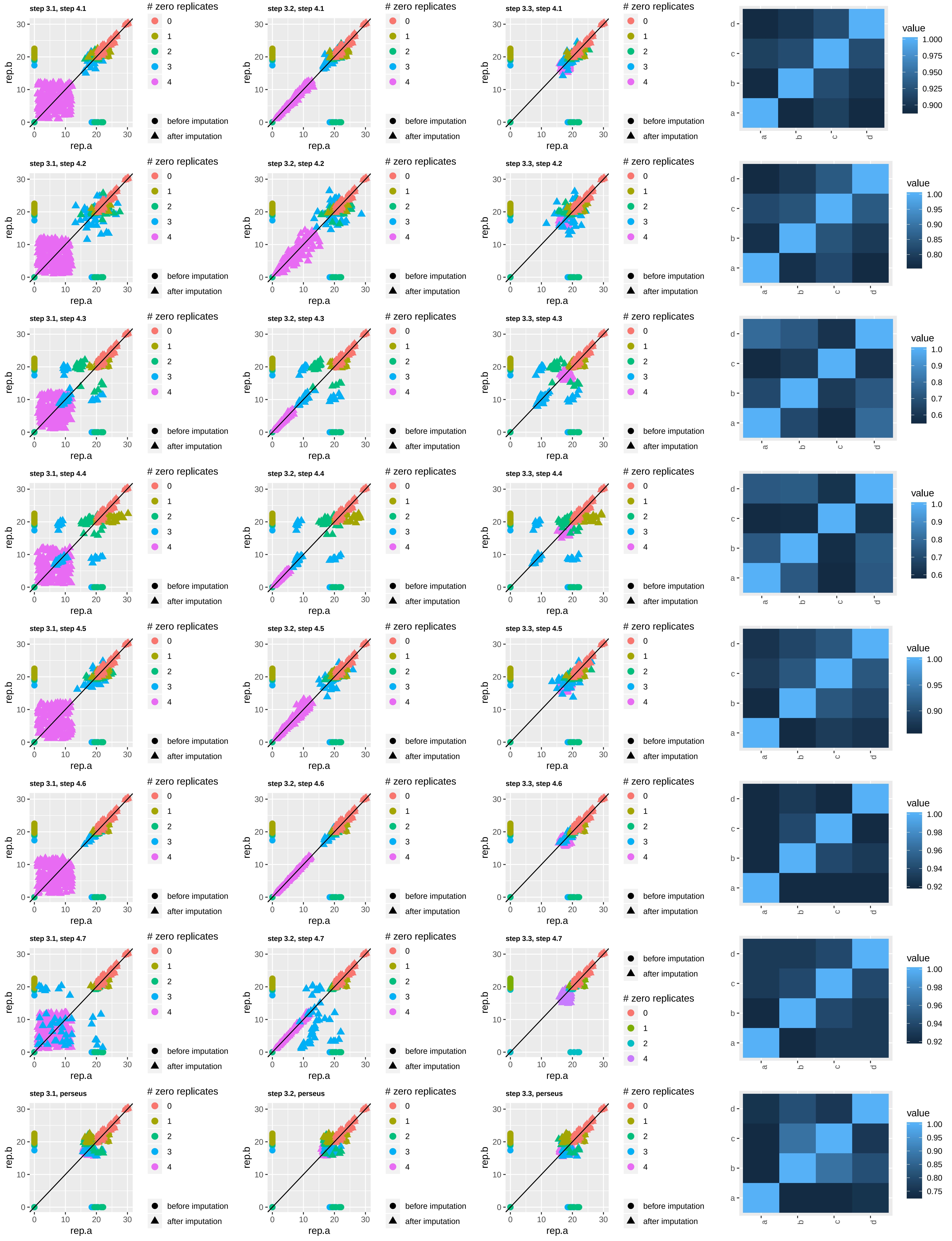

NCBP1−LAP, multi-well, condition 20 cases

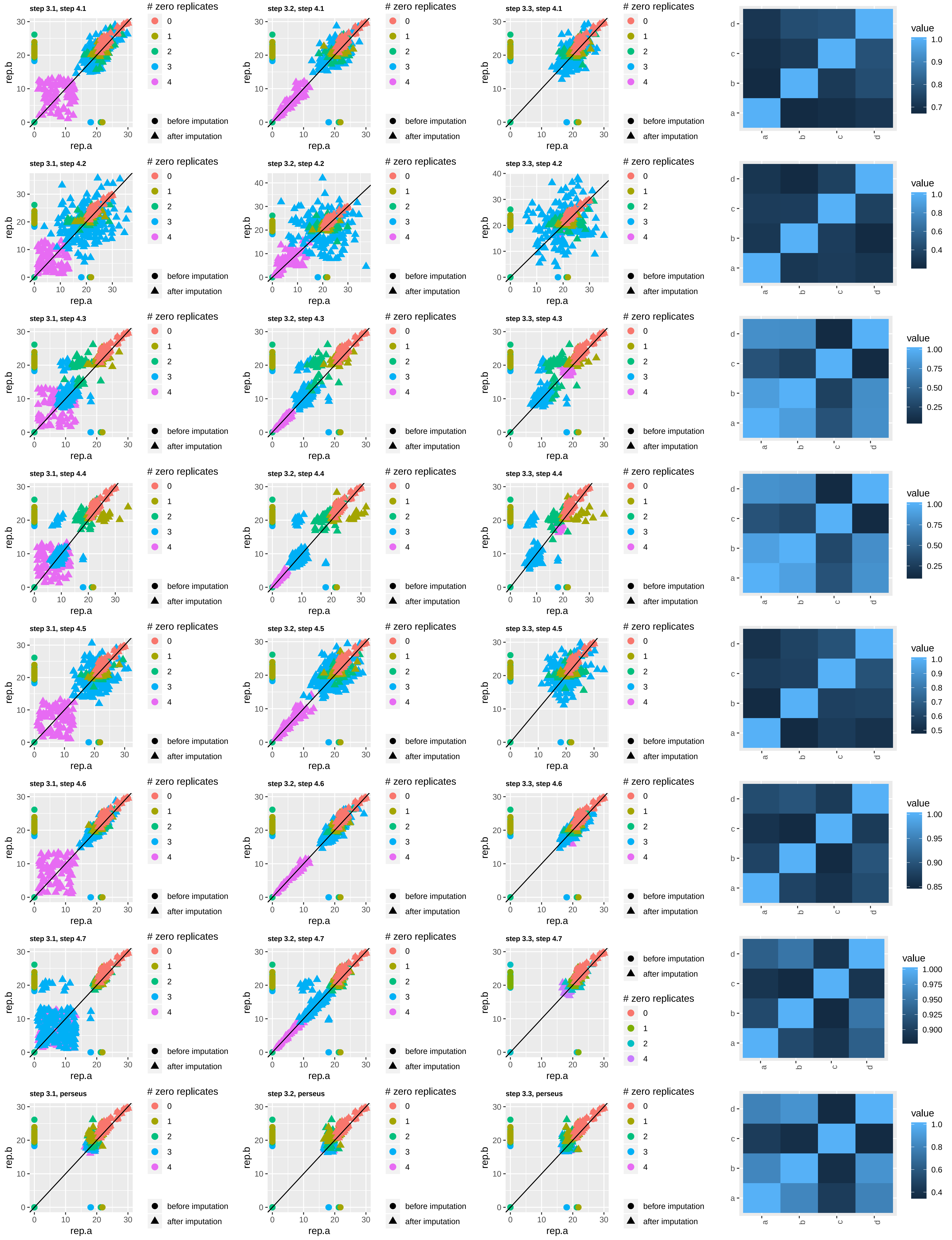

NCBP1−LAP, multi-well, condition 20

controls

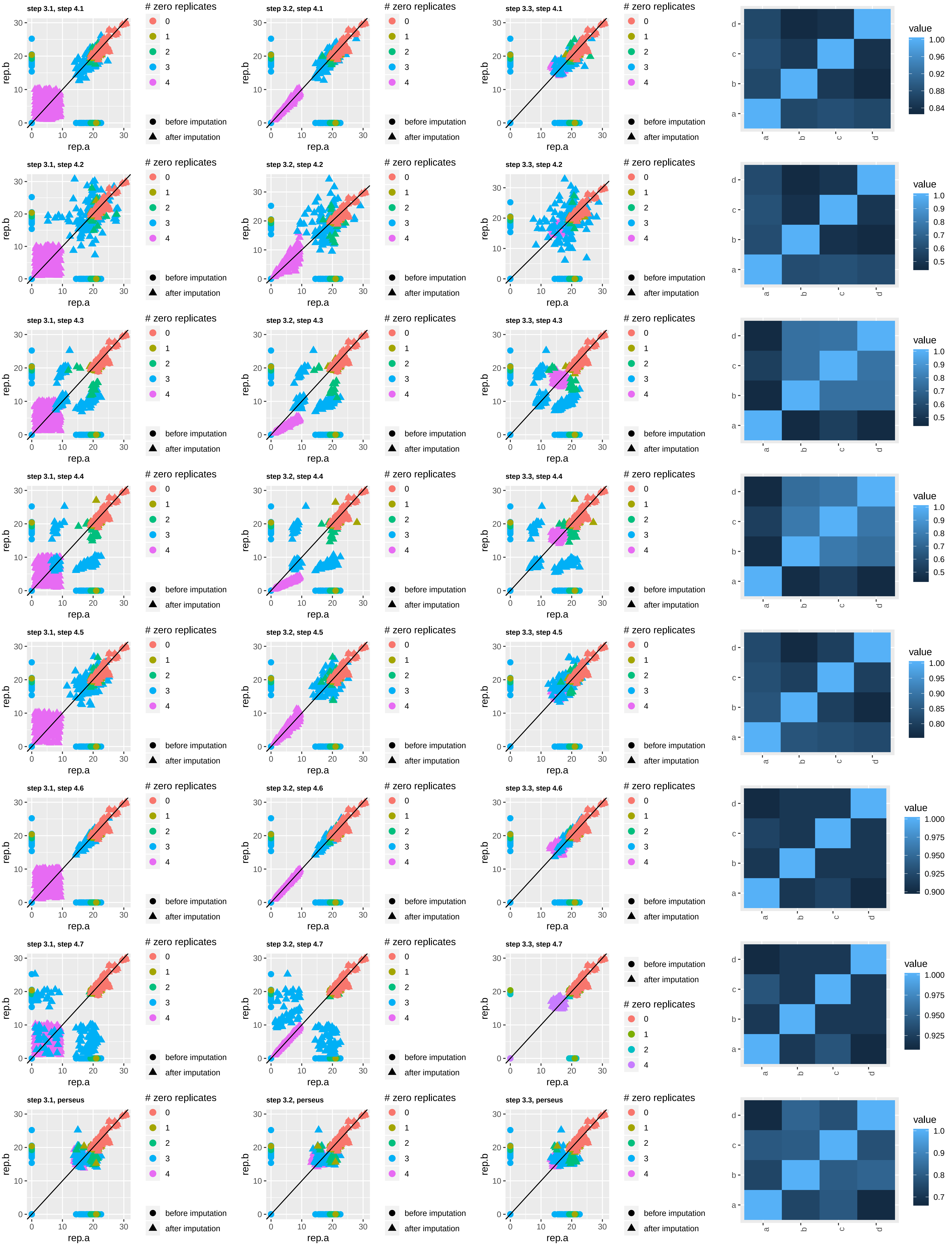

NCBP2−LAP, multi−well, condition 7

cases

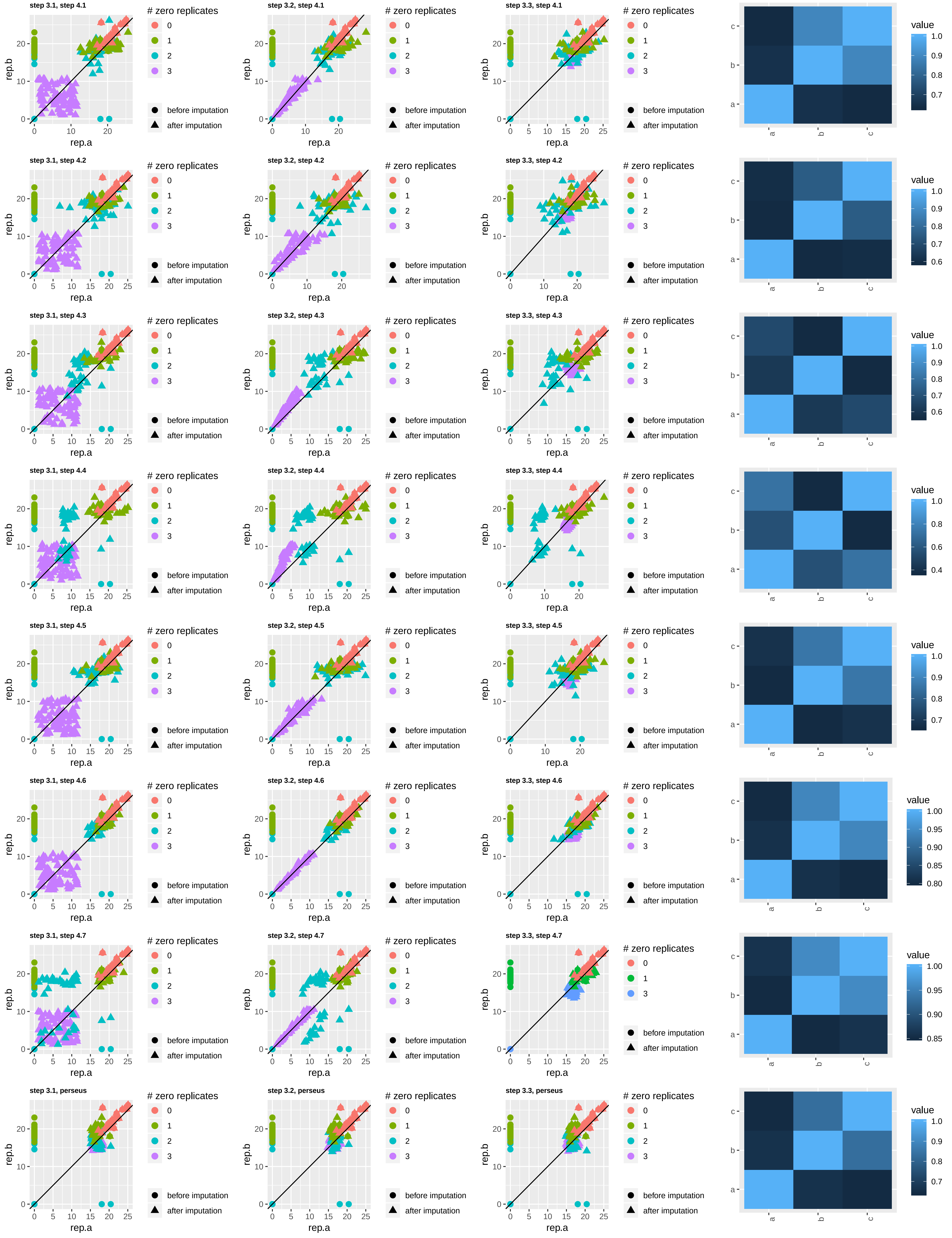

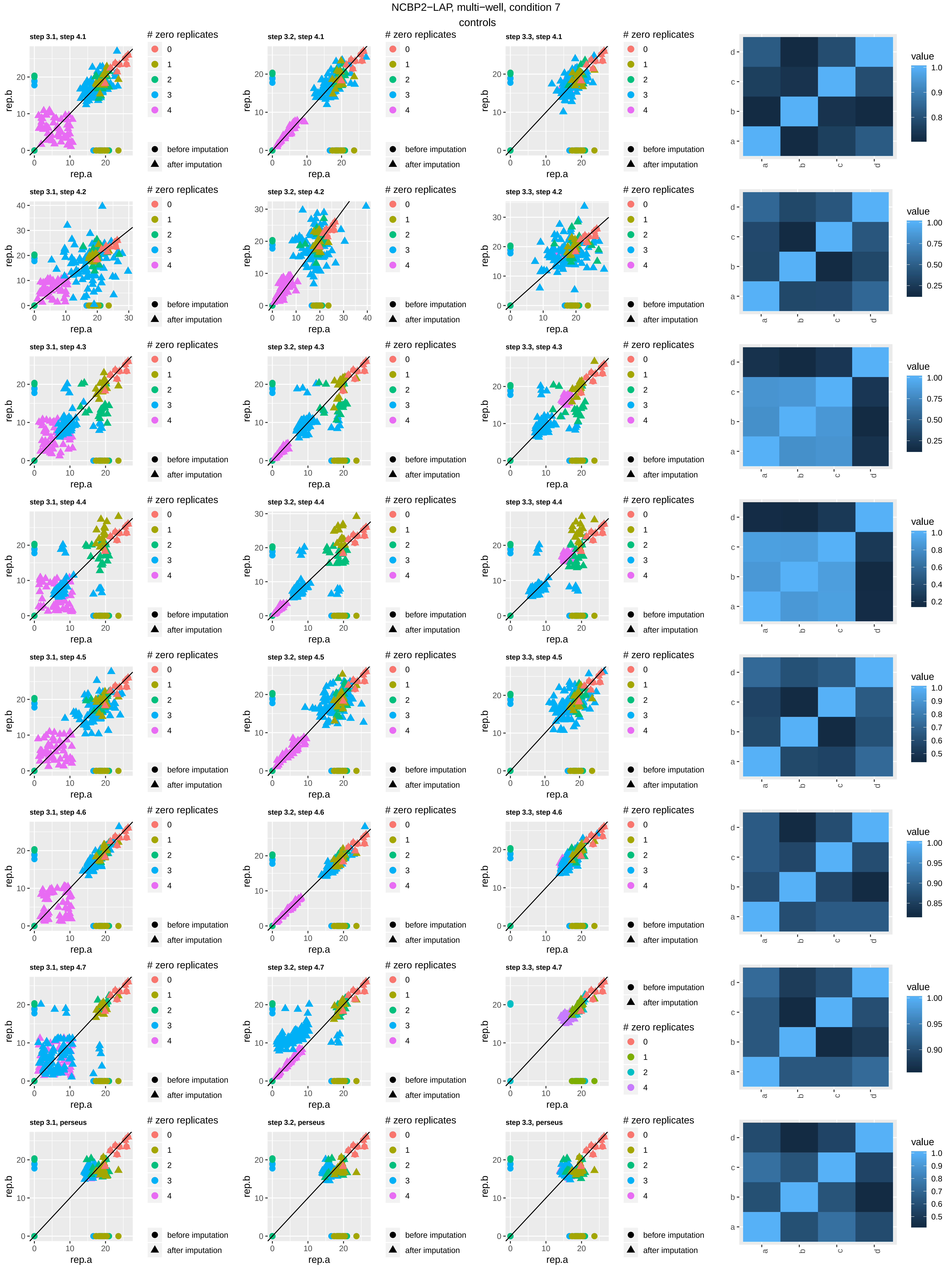

NCBP2−LAP, multi−well, condition 7
controls
step 3.1, step 4.1
step 3.2, step 4.1
step 3.3, step 4.1
step 3.1, step 4.2
step 3.2, step 4.2
step 3.3, step 4.2
step 3.1, step 4.3
step 3.2, step 4.3
step 3.3, step 4.3
step 3.1, step 4.4
step 3.2, step 4.4
step 3.3, step 4.4
step 3.1, step 4.5
step 3.2, step 4.5
step 3.3, step 4.5
step 3.1, step 4.6
step 3.2, step 4.6
step 3.3, step 4.6
step 3.1, step 4.7
step 3.2, step 4.7
step 3.3, step 4.7
step 3.1, perseus
step 3.2, perseus
step 3.3, perseus
rep.a
rep.b
### zero replicates
0
1
2
3
4
before imputation
after imputation
value
a
b
c
d

NCBP2−LAP, multi−well, condition 10
cases

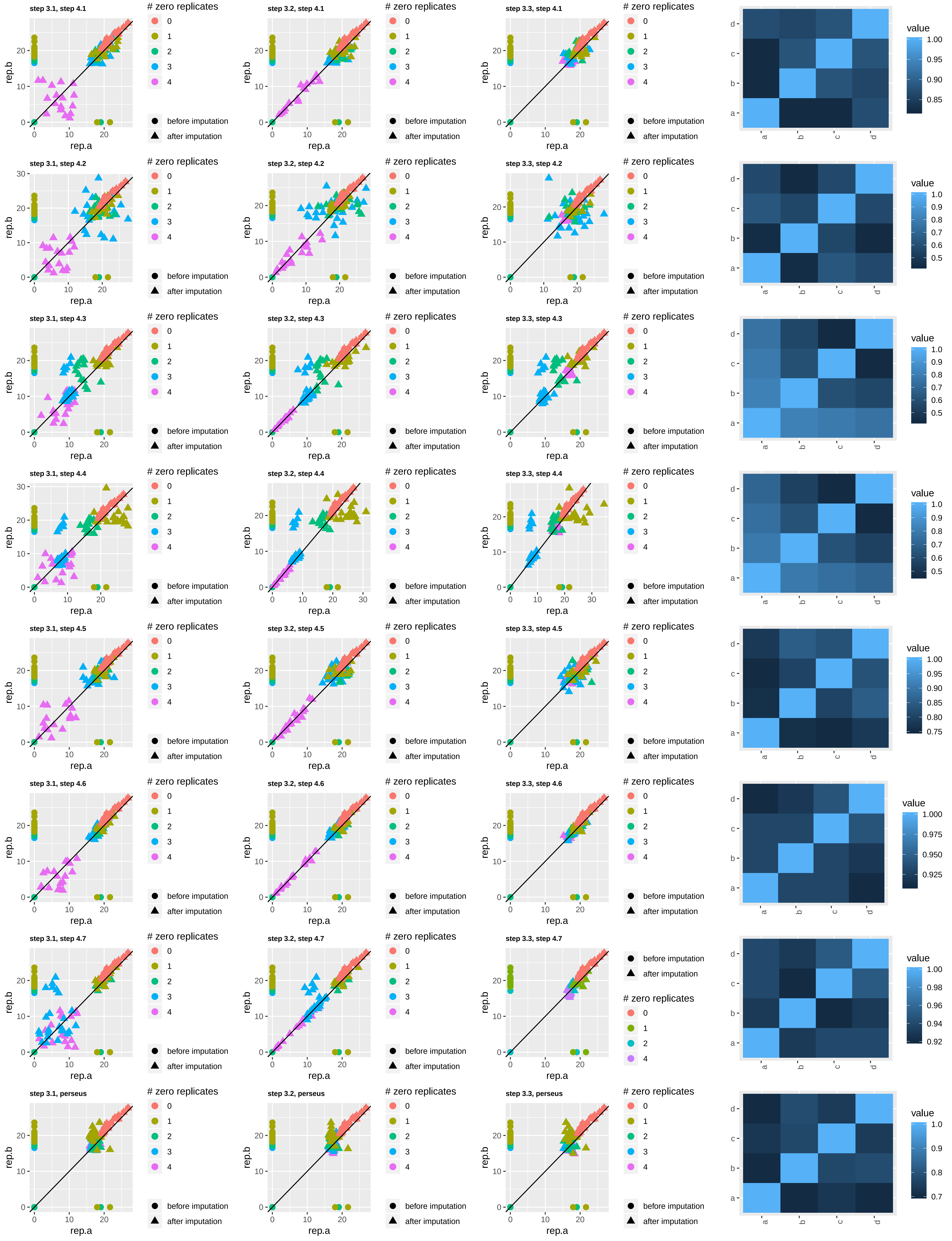

NCBP2−LAP, multi-well, condition 10

controls

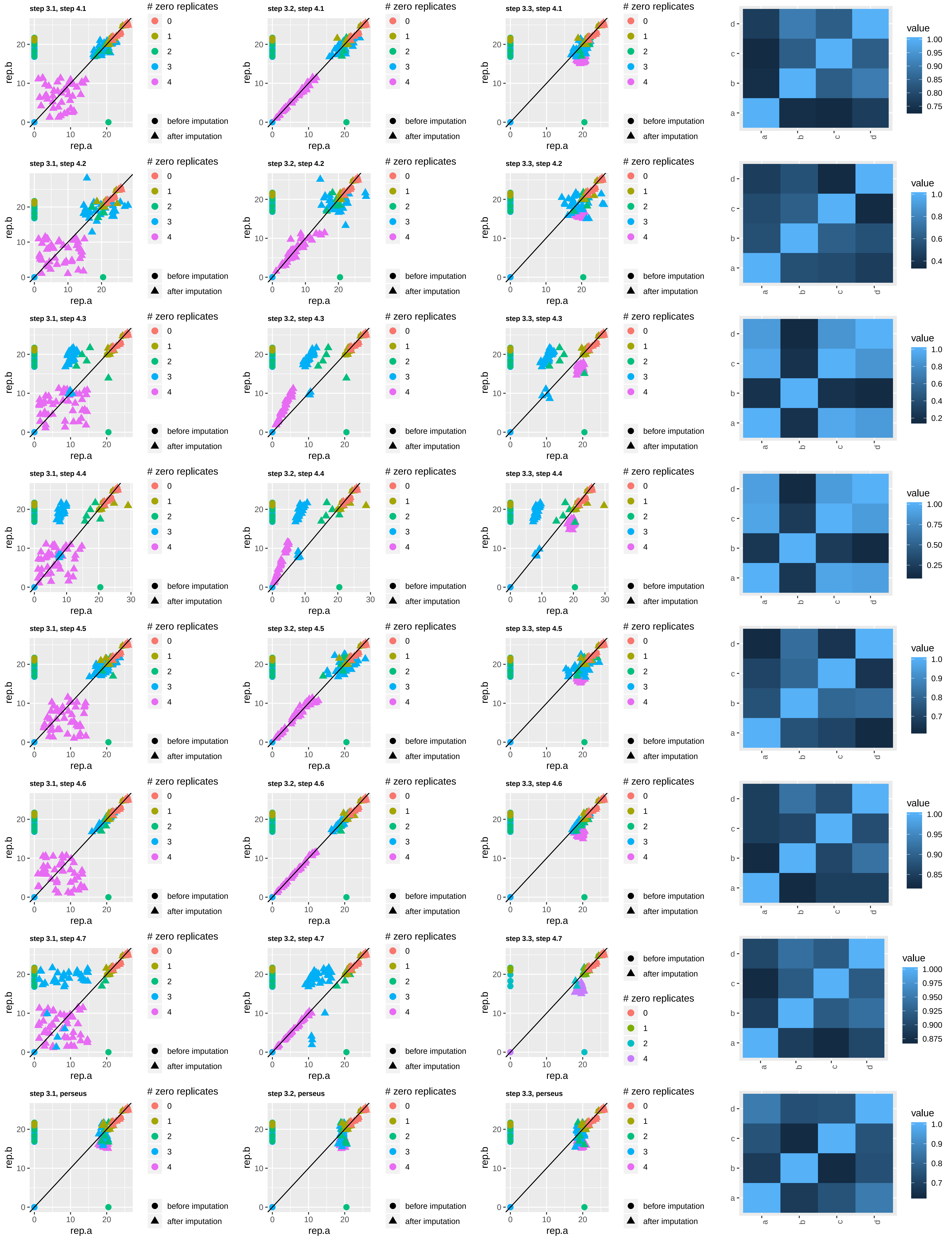

NCBP2−LAP, multi-well, condition 12
cases

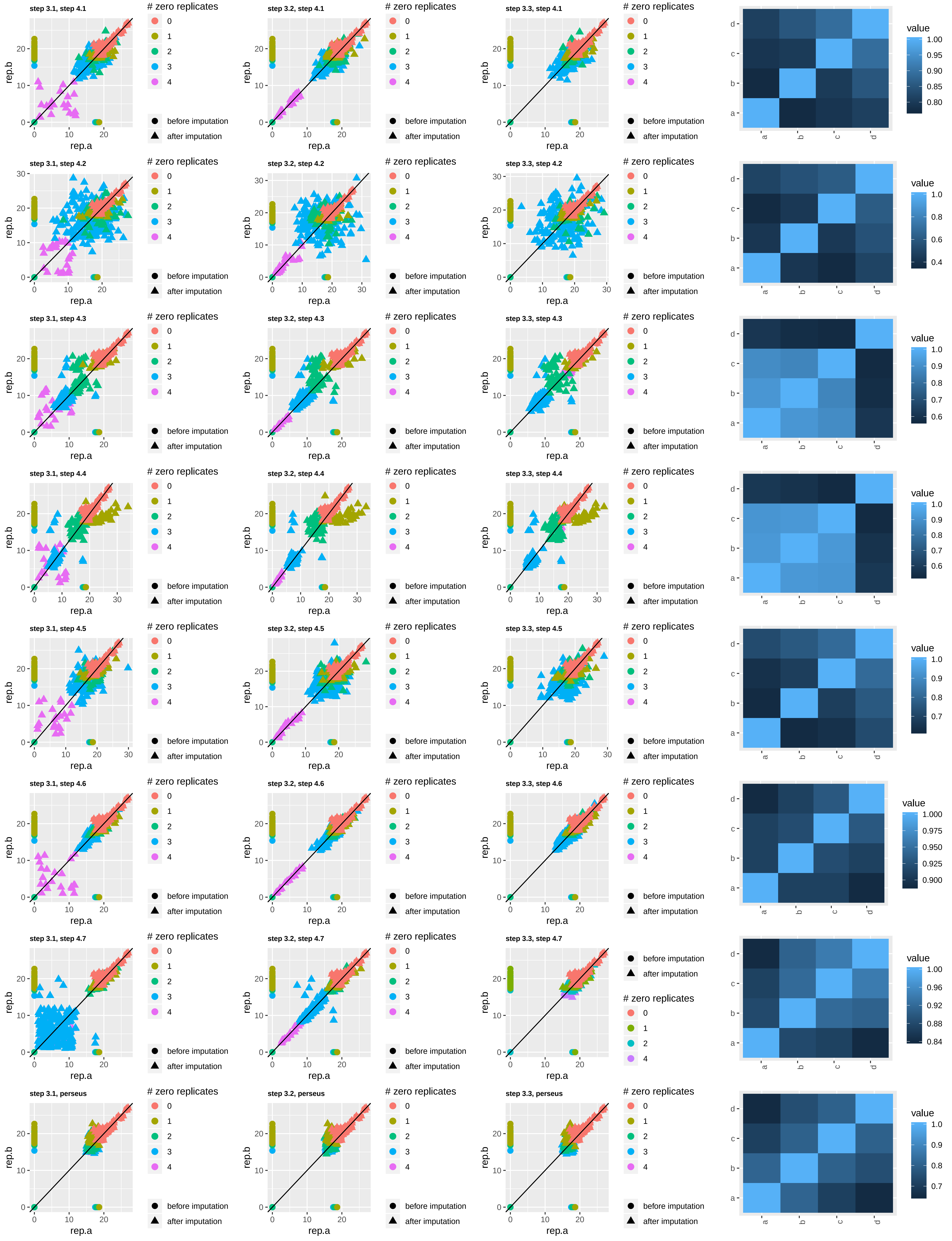

NCBP2−LAP, multi−well, condition 12

controls

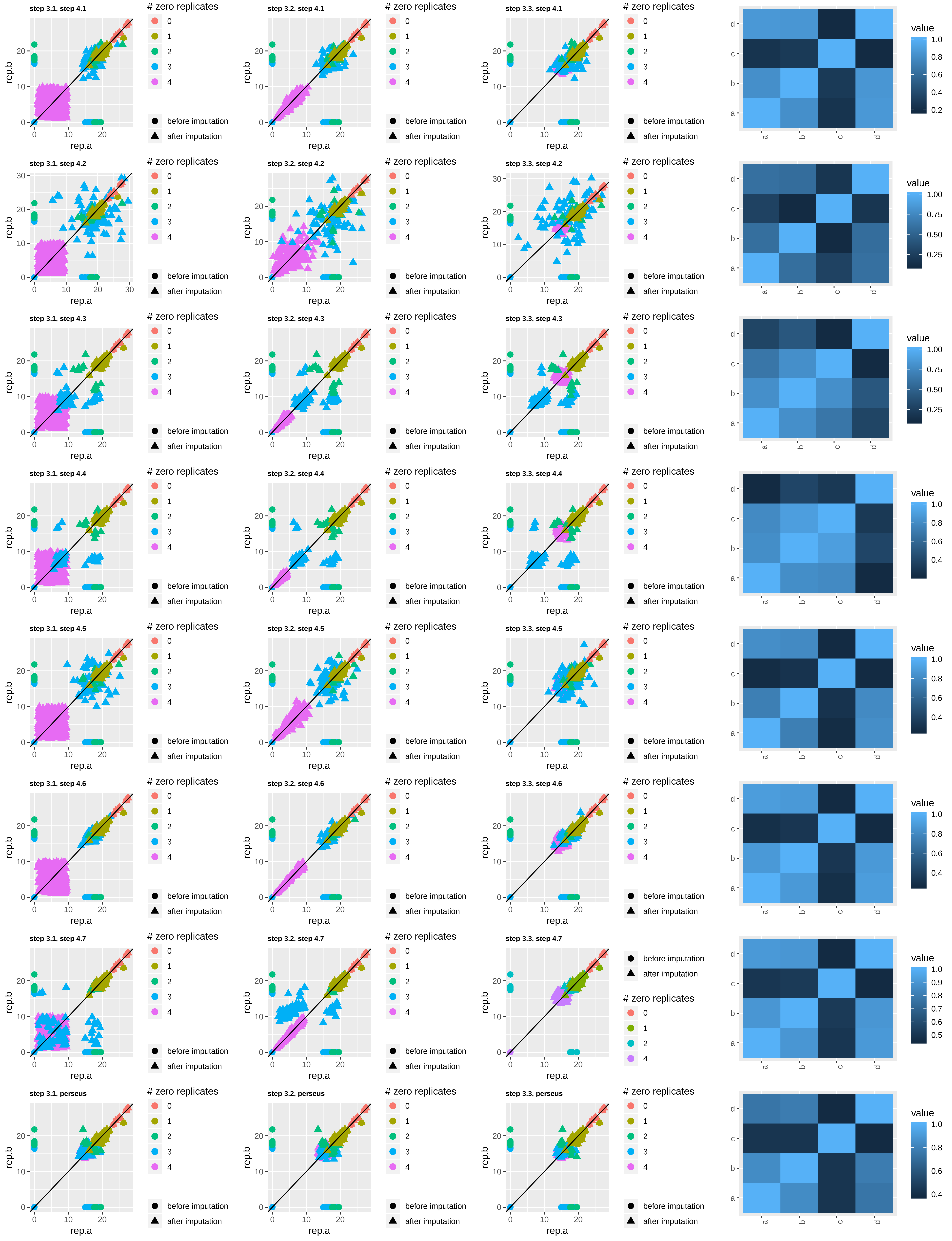

NCBP2−LAP, multi-well, condition 14 cases

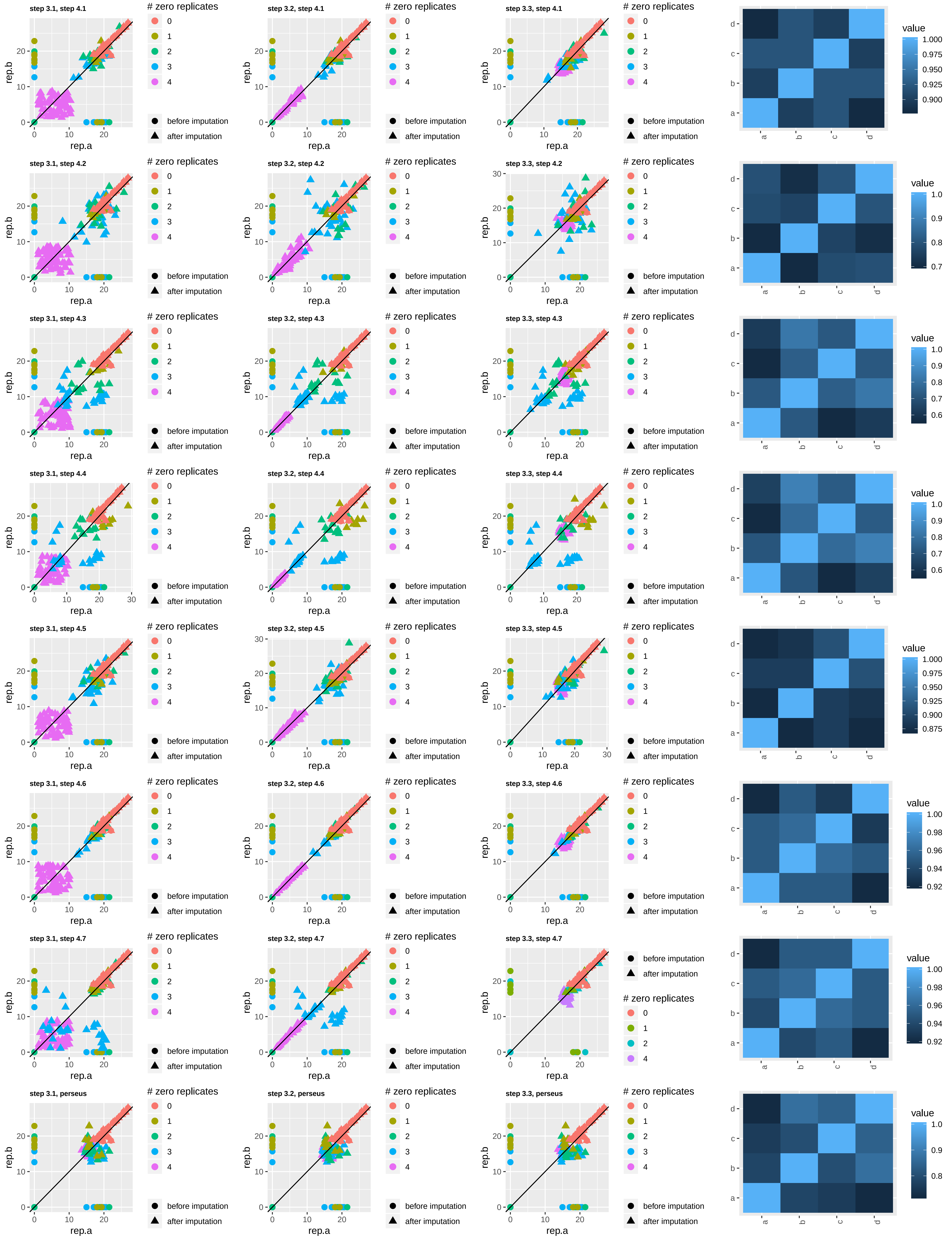

NCBP2−LAP, multi−well, condition 14

controls

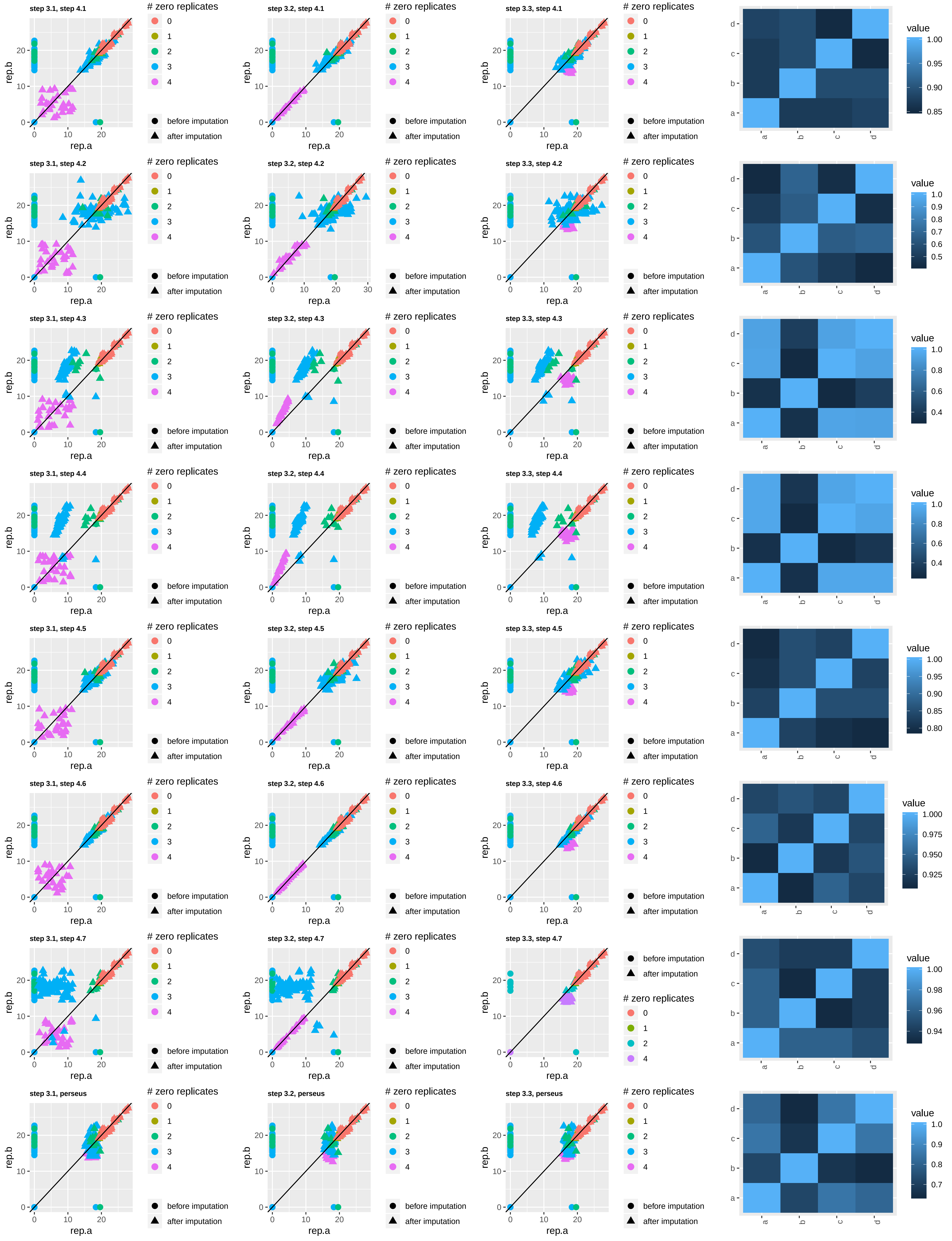

NCBP2−LAP, multi−well, condition 18

cases

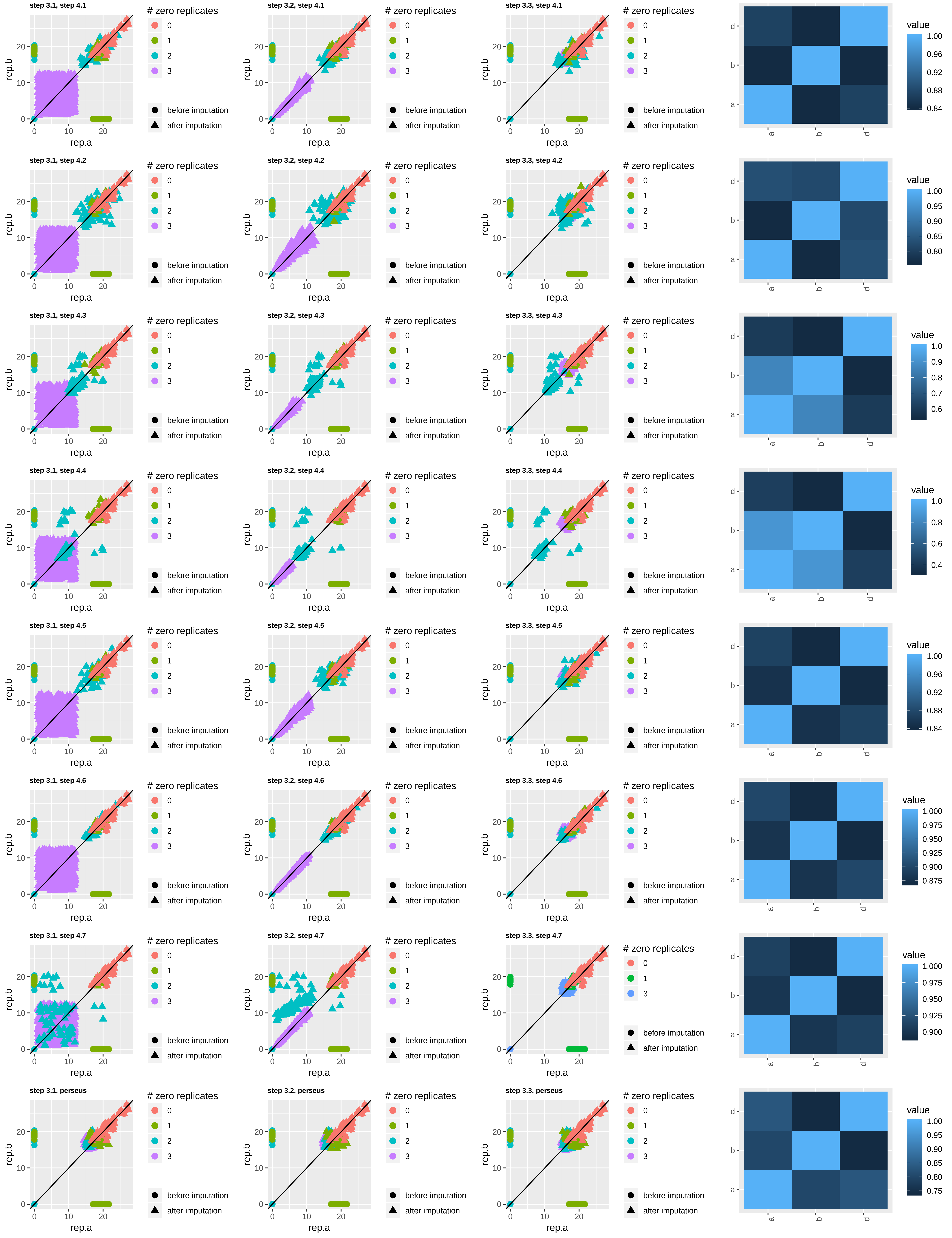

NCBP2−LAP, multi−well, condition 18

controls

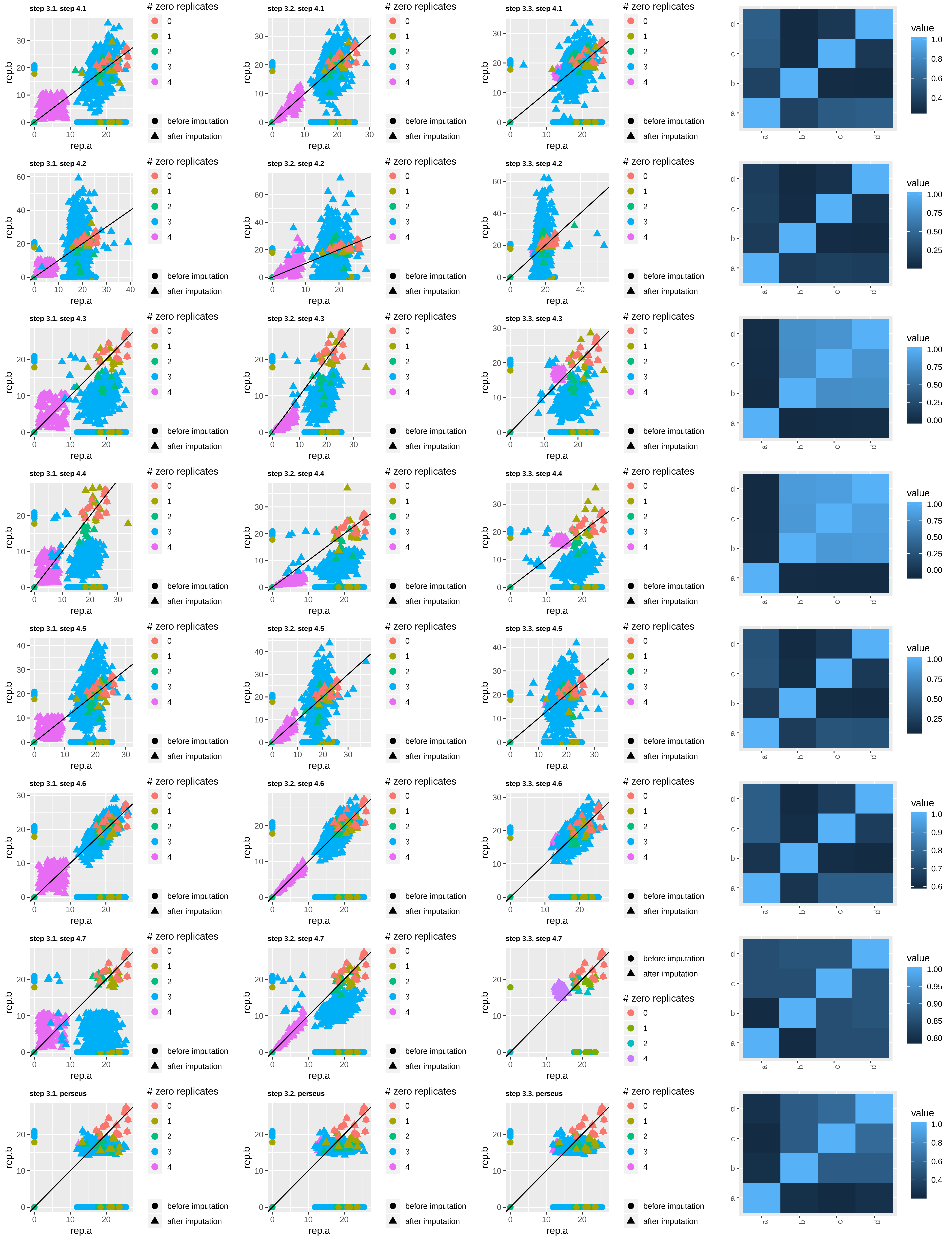

NCBP2−LAP, multi-well, condition 20

cases

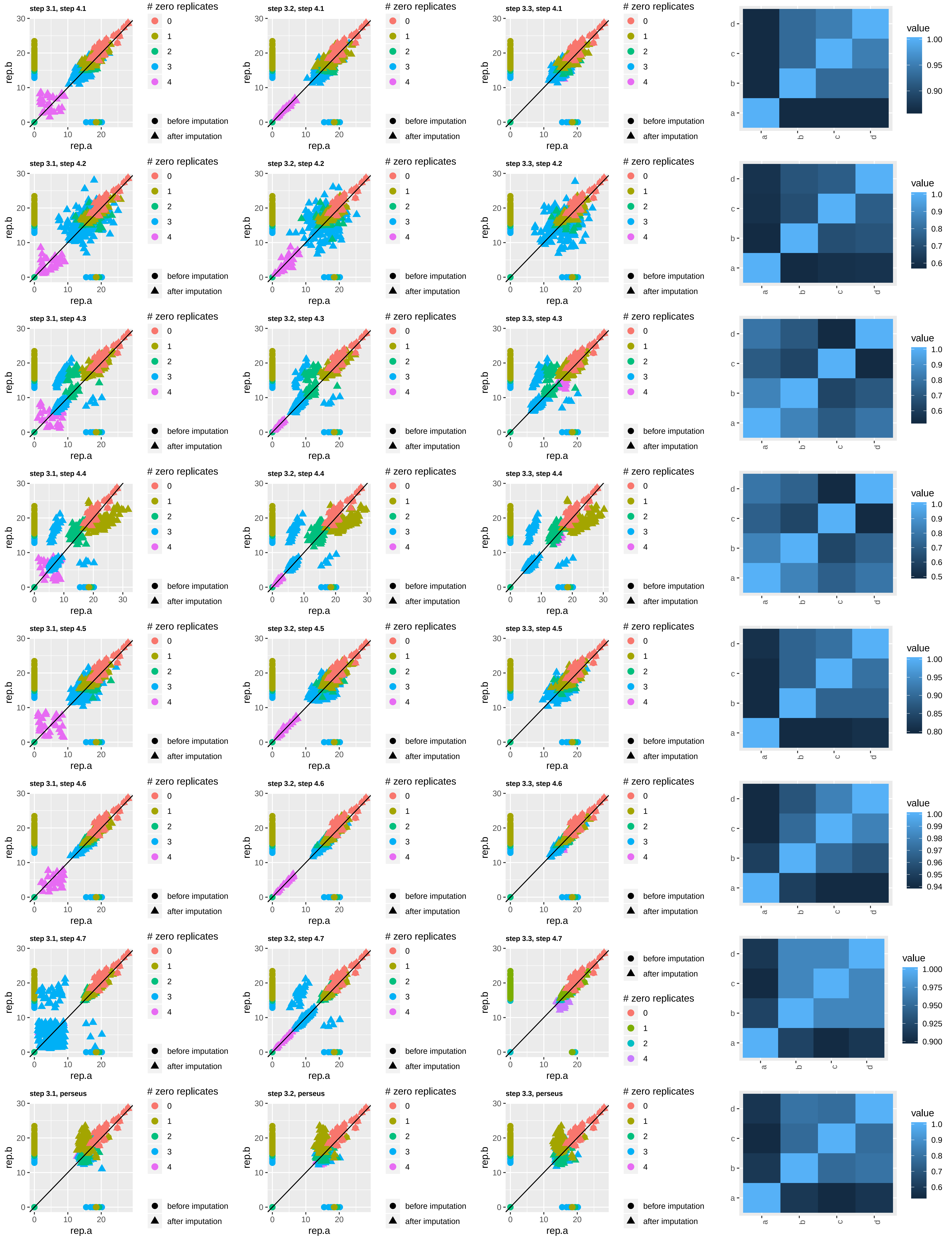

NCBP2−LAP, multi−well, condition 20

controls

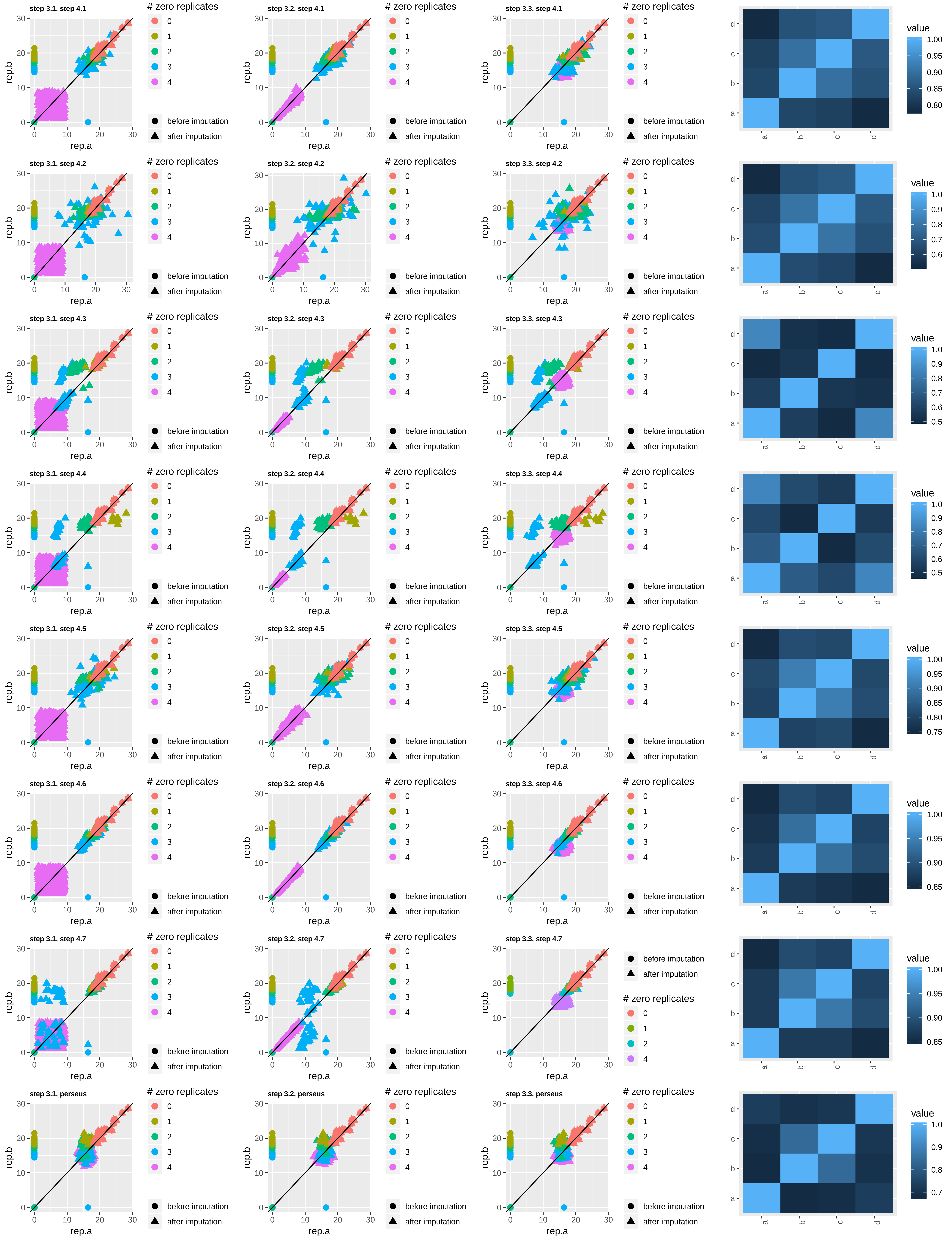

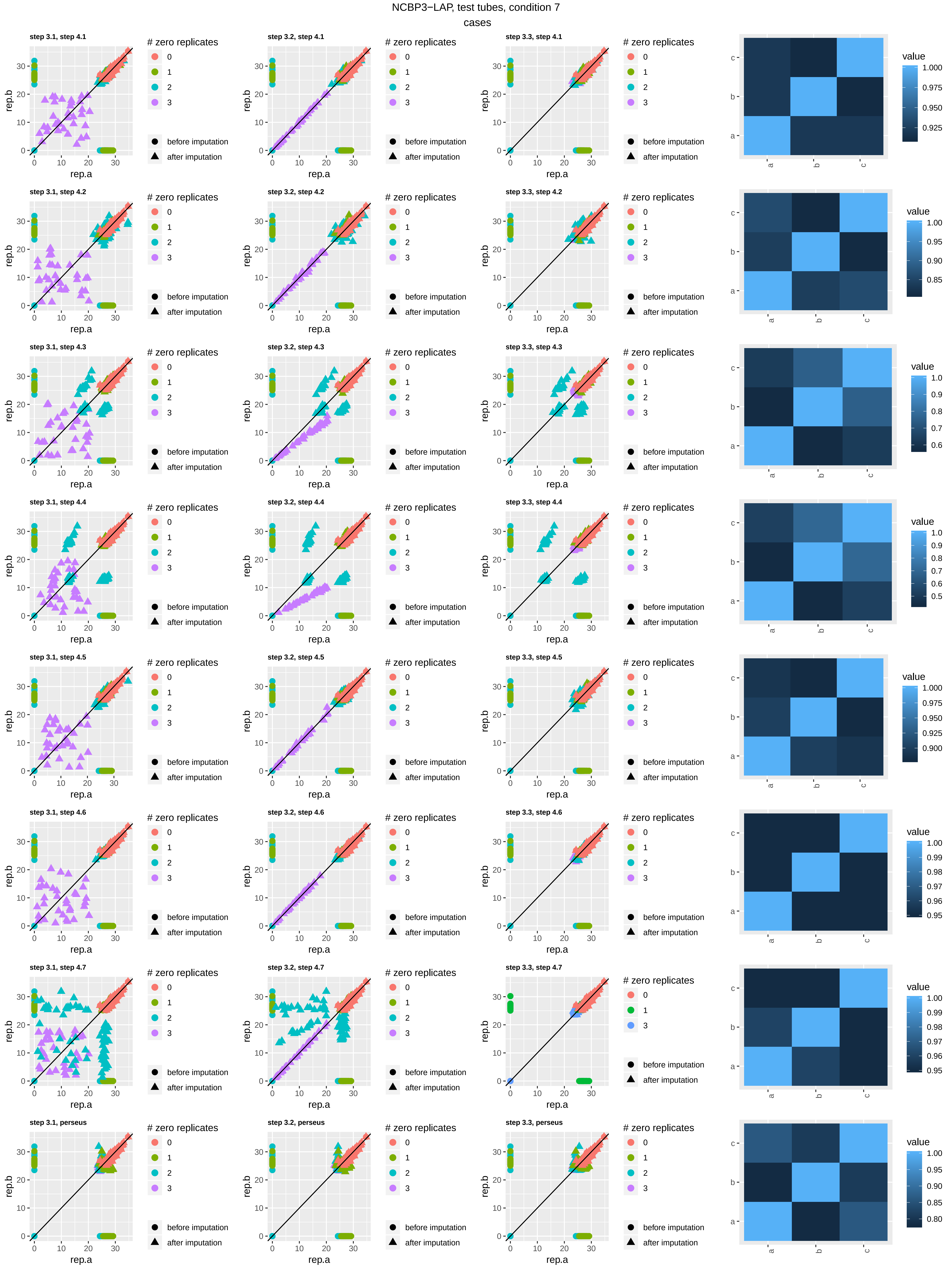
NCBP3−LAP, test tubes, condition 7
cases
step 3.1, step 4.1
step 3.2, step 4.1
step 3.3, step 4.1
step 3.1, step 4.2
step 3.2, step 4.2
step 3.3, step 4.2
step 3.1, step 4.3
step 3.2, step 4.3
step 3.3, step 4.3
step 3.1, step 4.4
step 3.2, step 4.4
step 3.3, step 4.4
step 3.1, step 4.5
step 3.2, step 4.5
step 3.3, step 4.5
step 3.1, step 4.6
step 3.2, step 4.6
step 3.3, step 4.6
step 3.1, step 4.7
step 3.2, step 4.7
step 3.3, step 4.7
step 3.1, perseus
step 3.2, perseus
step 3.3, perseus
rep.a
rep.b
### zero replicates
0
1
2
3
before imputation
after imputation
value

NCBP3−LAP, test tubes, condition 7

controls

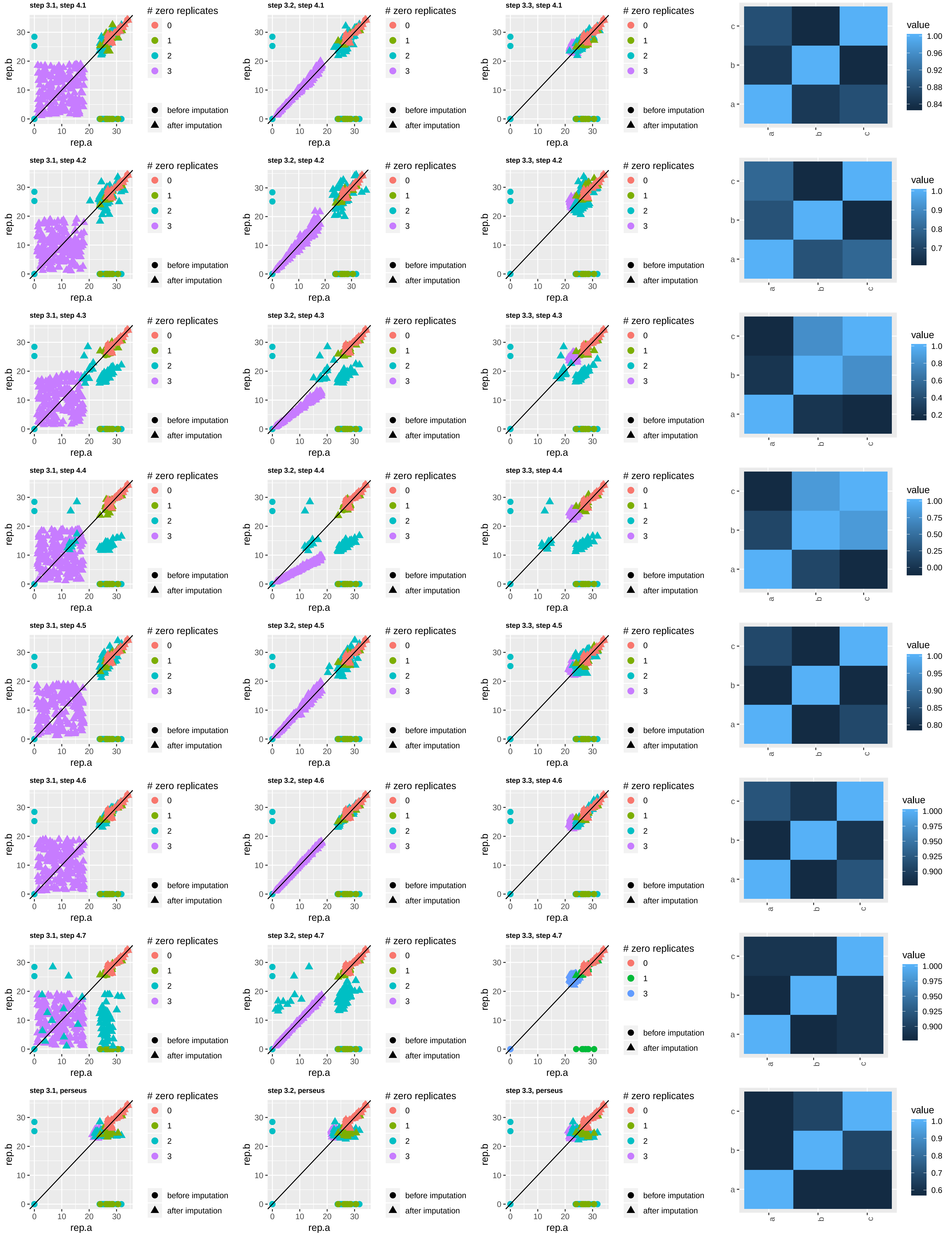

NCBP3−LAP, test tubes, condition 10

cases

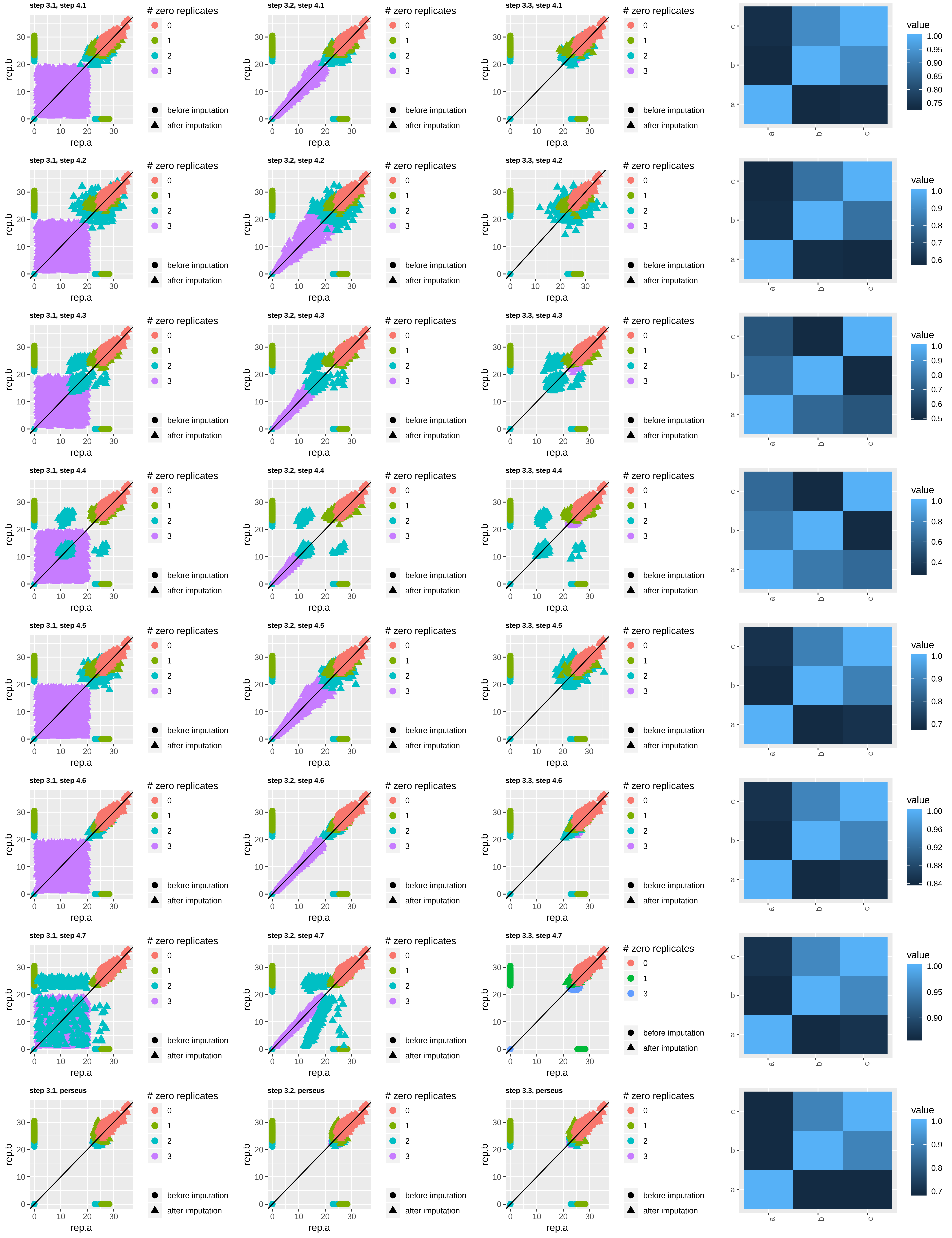

NCBP3−LAP, test tubes, condition 10

controls

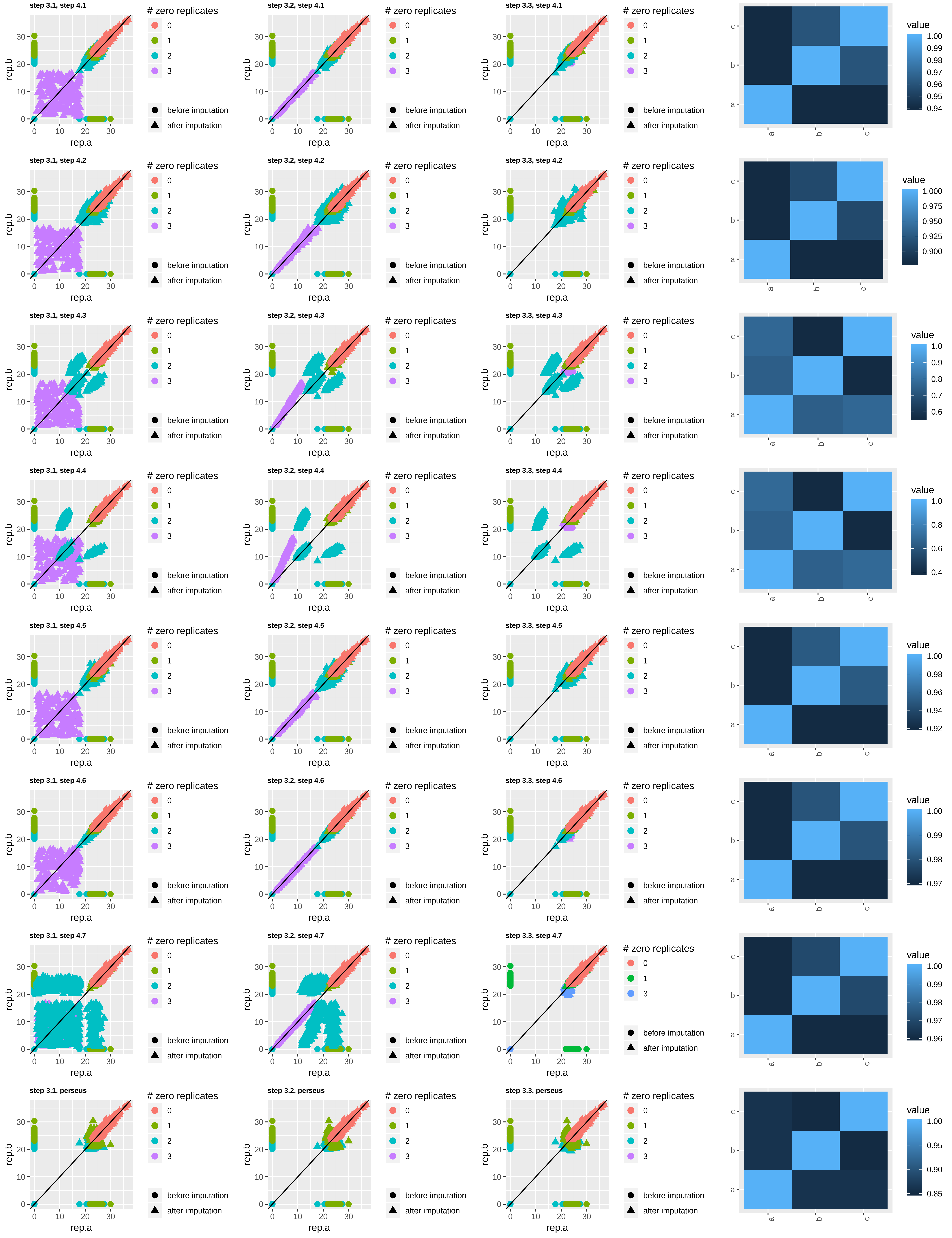

NCBP3−LAP, test tubes, condition 12

cases

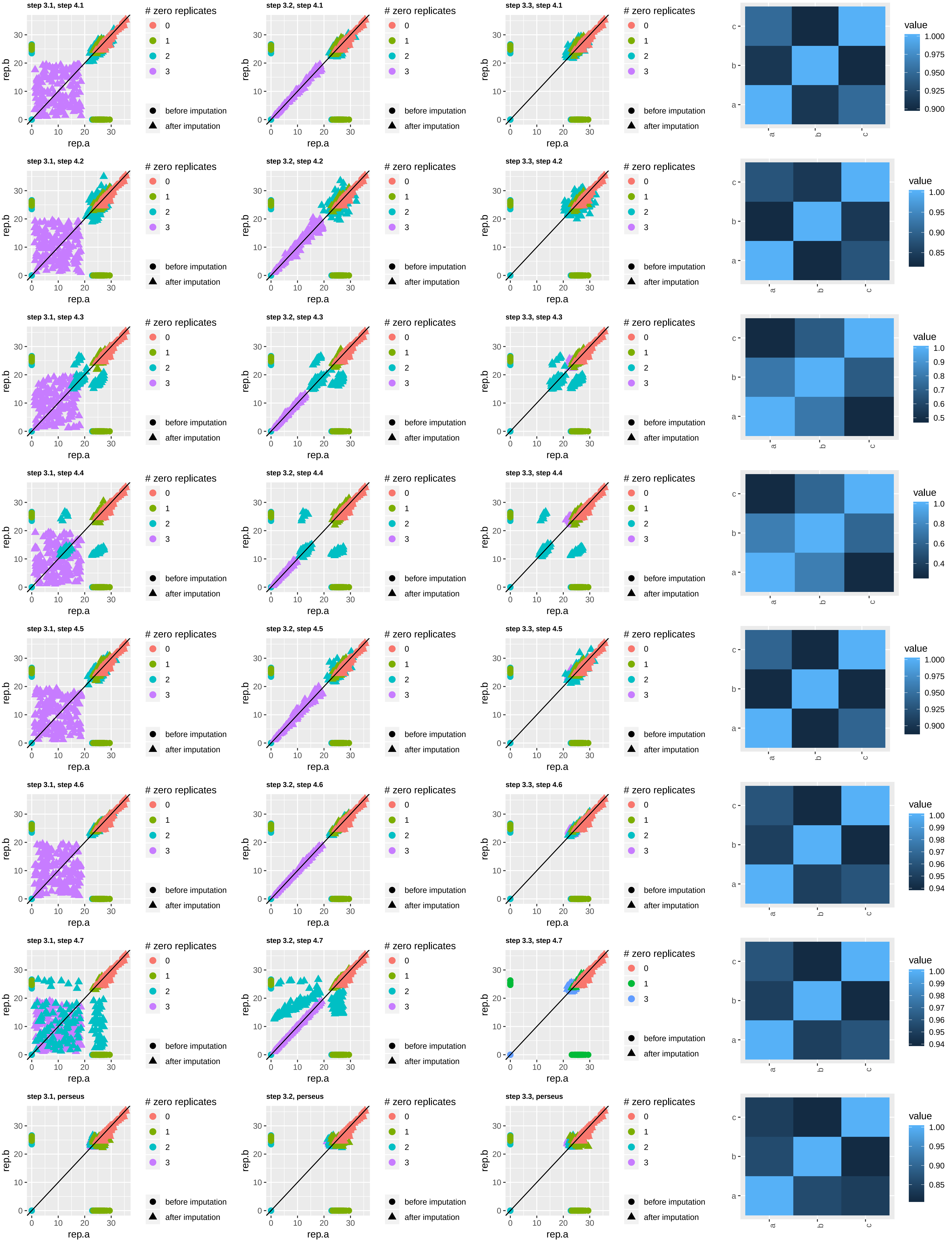

NCBP3−LAP, test tubes, condition 12

controls

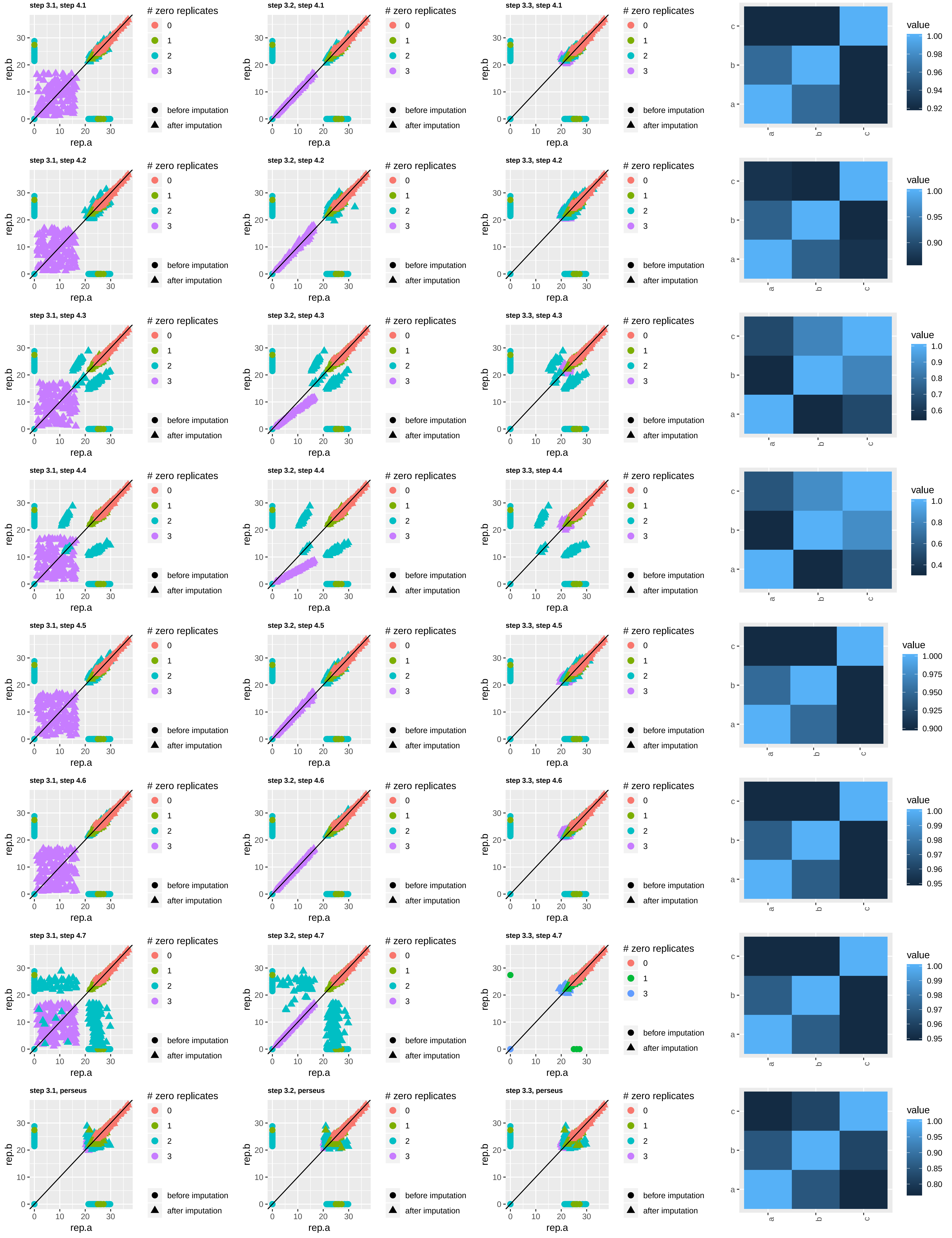

NCBP3−LAP, test tubes, condition 18

cases

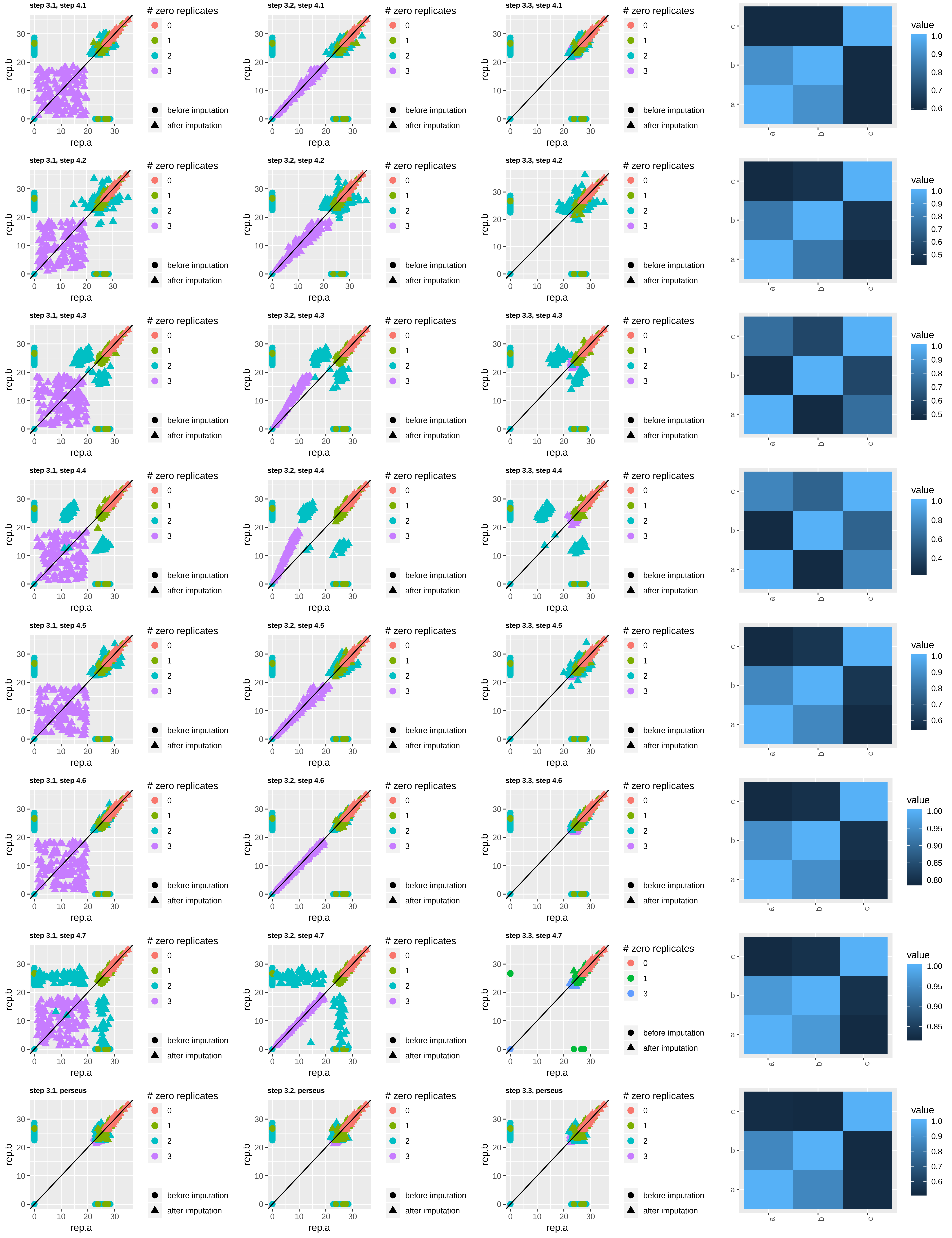

NCBP3−LAP, test tubes, condition 18

controls

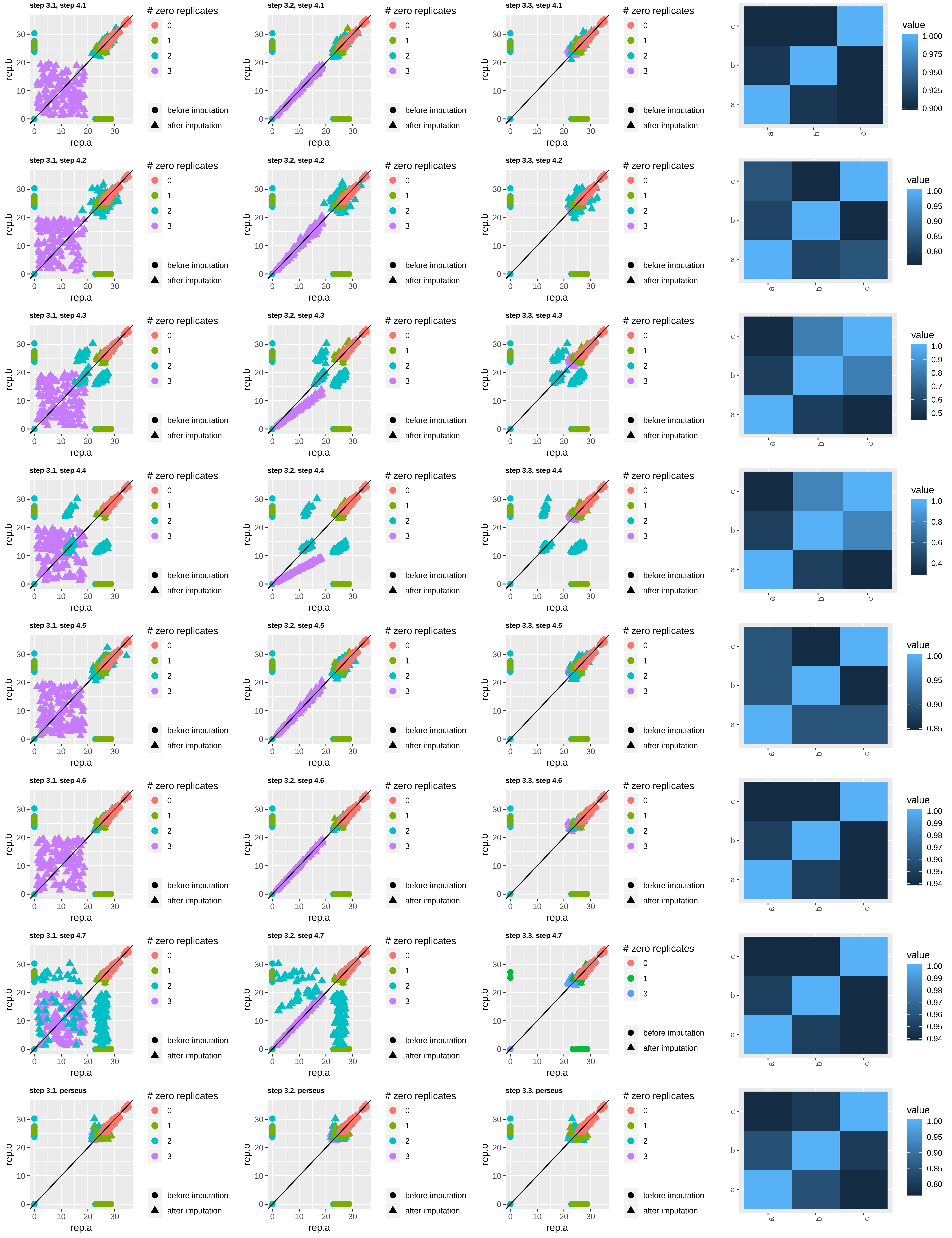
