## Supplementary figures and images for "Affinity proteomic dissection of the human nuclear cap-binding complex interactome"

### 3D_Animated_MDS.gif

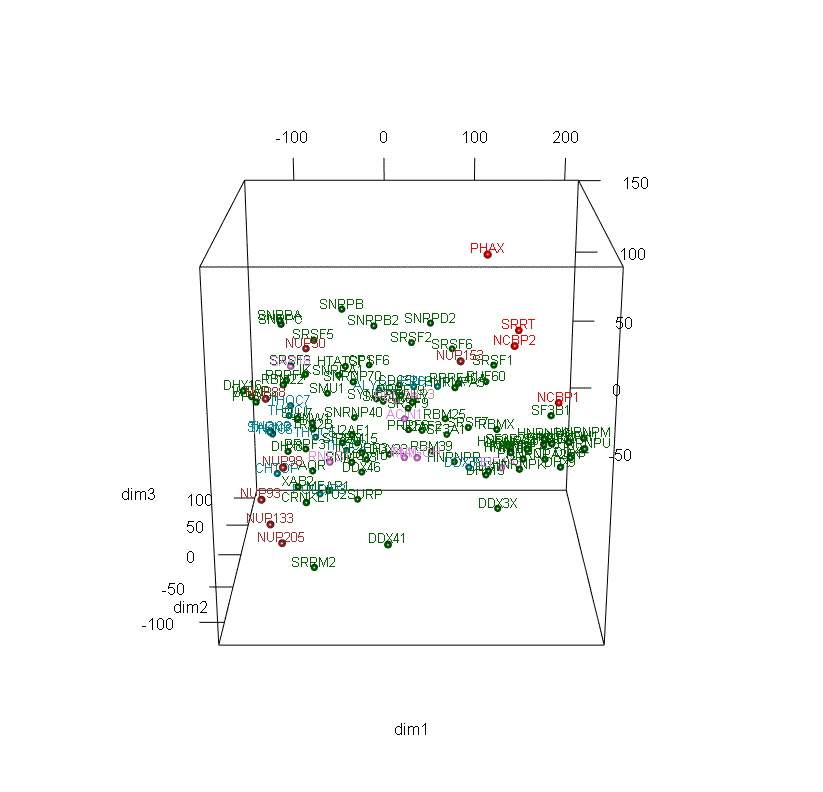

### GFP_normalization.pdf

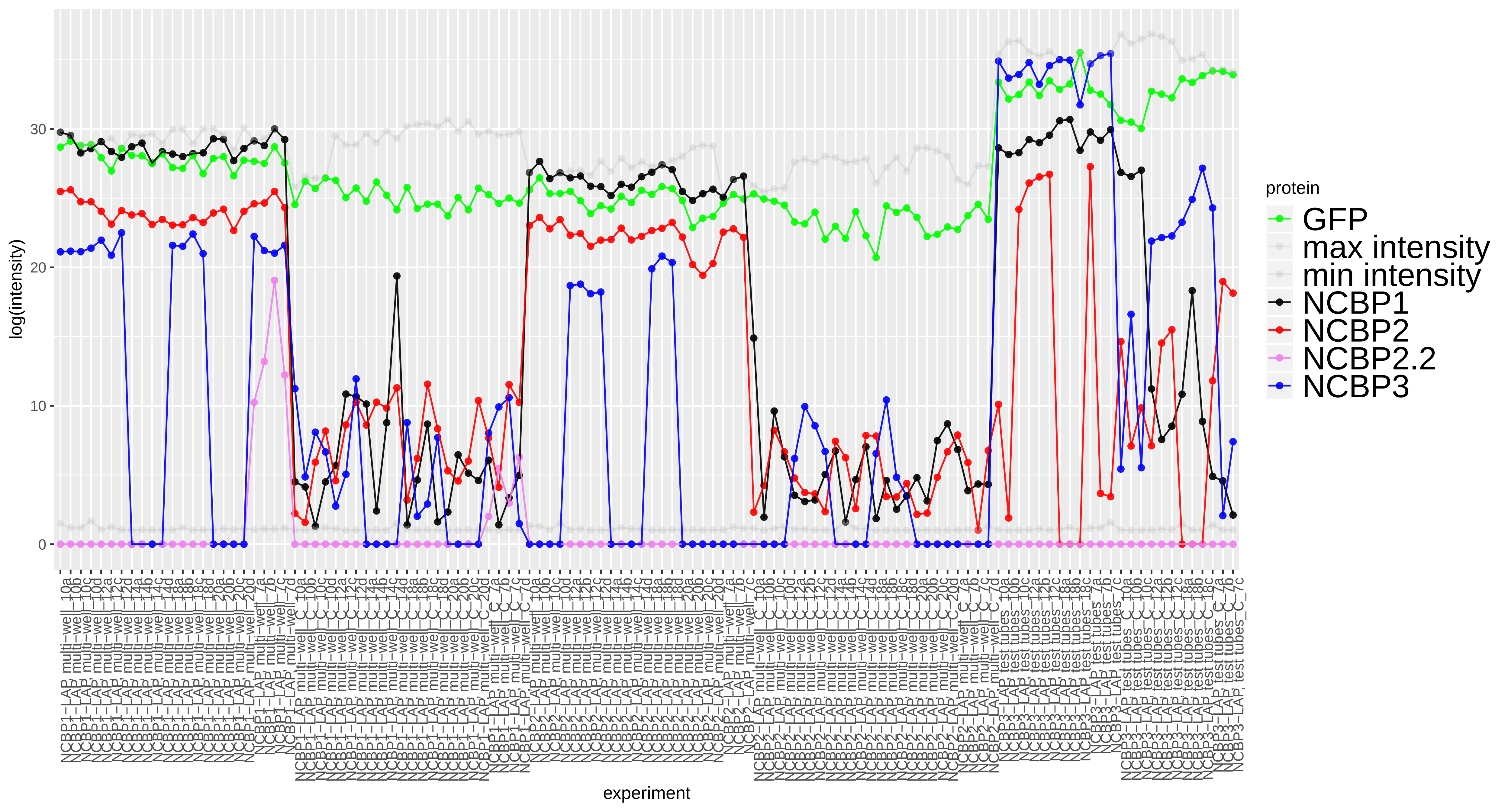

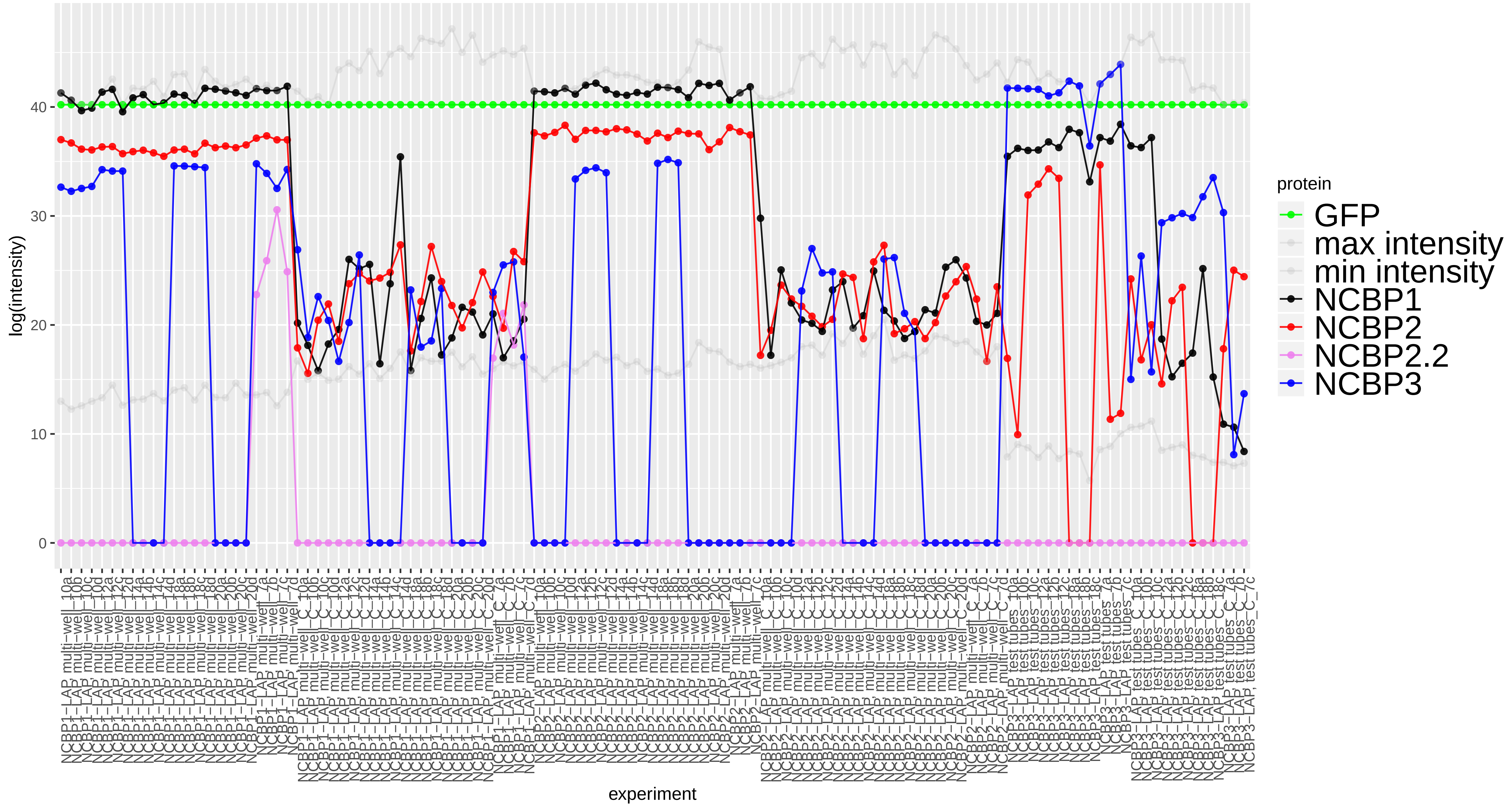

### Supplemental_heatmap.pdf

#### Supplemental Heatmap

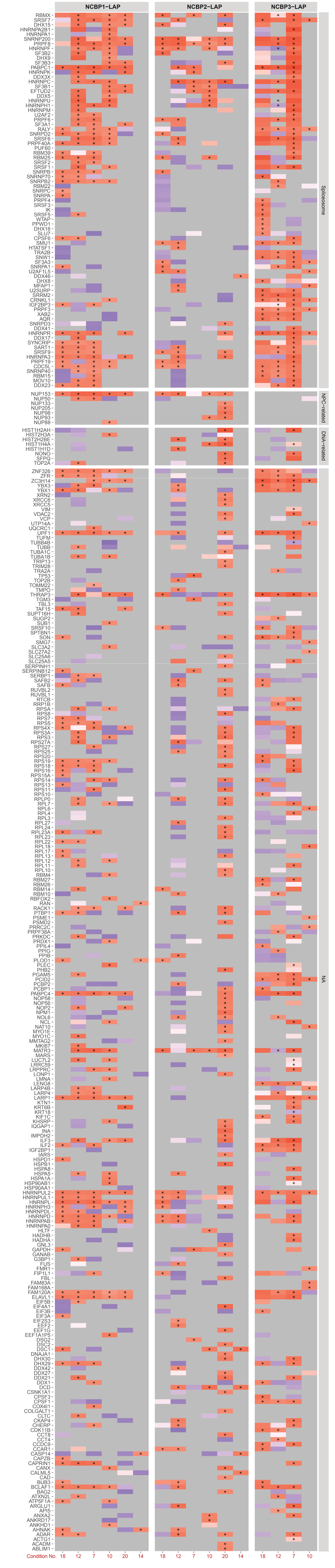
